# A Large Scale Joint Analysis of Flowering Time Reveals Independent Temperate Adaptations in Maize

**DOI:** 10.1101/086082

**Authors:** Kelly Swarts, Eva Bauer, Jeffrey C. Glaubitz, Tiffany Ho, Lynn Johnson, Yongxiang Li, Yu Li, Zachary Miller, Cinta Romay, Chris-Carolin Schöen, Tianyu Wang, Zhiwu Zhang, Edward S. Buckler, Peter Bradbury

## Abstract

Modulating days to flowering is a key mechanism in plants for adapting to new environments, and variation in days to flowering drives population structure by limiting mating. To elucidate the genetic architecture of flowering across maize, a quantitative trait, we mapped flowering in five global populations, a diversity panel (Ames) and four half-sib mapping designs, Chinese (CNNAM), US (USNAM), and European Dent (EUNAM-Dent) and Flint (EUNAM-Flint). Using whole-genome projected SNPs, we tested for joint association using GWAS, resampling GWAS and two regional approaches; Regional Heritability Mapping (RHM) (*1*, *2*) and a novel method, Boosted Regional Heritability Mapping (BRHM). Direct overlap in significant regions detected between populations and flowering candidate genes was limited, but whole-genome cross-population predictive abilities were ≤0.78. Poor predictive ability correlated with increased population differentiation (r = 0.41), unless the parents were broadly sampled from across the North American temperate-tropical germplasm gradient; uncorrected GWAS results from populations with broadly sampled parents were well predicted by temperate-tropical F_ST_s in machine learning. Machine learning between GWAS results also suggested shared architecture between the American panels and, more distantly, the European panels, but not the Chinese panel. Machine learning approaches can reconcile non-linear relationships, but the combined predictive ability of all of the populations did not significantly enhance prediction of physiological candidates. While the North American-European temperate adaption is well studied, this study suggest independent temperate adaptation evolved in the Chinese panel, most likely in China after 1500, a finding supported by differential gene ontology term enrichment between populations.

## Background

This study sought to refine our understanding of the genetic architecture underlying maize flowering by combining diverse populations genotyped using whole genome resequencing data. One of the easiest ways for a plant to adapt to a new environment is to modulate the time until flowering, avoiding obstacles to reproductive success. However, changes in flowering lead to reproductive isolation, structuring populations and generating strong pleiotropic associations with other traits (*3*, *4*). This makes days to flowering an excellent trait to study the biological mechanism of adaptation, but confounds association mapping approaches.

### Flowering traits and population structure

Flowering time traits have received much attention in maize due to the quantitative nature of inheritance, high heritabilities (up to 0.96 for days to silking in the USNAM (*5*) population) and ease of scoring (*5*–*18*). Only a handful of loci, mostly in the autonomous flowering pathway, have confirmed effects for modulating days to flowering either through mutagenesis or positional cloning, including Rap2.7 and the MITE insertion at the Vgt1 locus (*7*, *19*), ZCN8 (*20*), ID1 (*21*, *22*), conz1 (*23*), df1 (*24*), zfl1 and zfl2 (*25*) and the major photoperiod locus, ZmCCT (*10*). However, independent days to flowering QTL have been mapped in tens of biparental, RIL, and other structured population designs (*8*, *12*), including four NAM populations (*5*, *26*, *27*) and in diversity populations (the largest being the Ames Diversity Panel (*11*)); and of the list above, only Vgt1, ZCN8, and ZmCCT have common standing variation and are routinely detected in mapping populations (*5*, *8*, *13*). Flowering time is also of interest for breeding, as shifting days to flowering is one of the easiest ways to combat the effects of climate change (*28*–*31*), and is important for moving traits such as disease resistance or yield between tropical and temperate germplasm.

Previous studies (*5*, *8*, *12*, *15*, *26*, *32*, *33*) have confirmed that days to flowering is quantitative and highly additive in maize, and that many of the effects target common loci, with allelic series present at many loci; a meta-analysis of 22 linkage mapping and ANOVA studies record a total of 313 QTL that collapsed in 62 consensus regions of the genome (*12*). Synteny mapping to rice and Arabidopsis resulted in 19 overlapping associations (*12*), and synteny mapping with sorghum revealed that 92.5% of QTL found in sorghum were less than 10Mbp from a corresponding QTL in USNAM (*33*). A later meta-analysis including 29 studies (*8*), found 441 significant QTL could be collapsed into 59 genomic regions. Similarly, days to flowering in the first NAM population, USNAM, found over 50 genomic regions with significance, and additionally confirmed the existence of allelic series at common loci. A recent, large multi-environment evaluation of maize flowering in the USNAM, Ames diversity panel and a newly developed Chinese NAM (CNNAM) (*27*) population using low-density GBS markers identified 130 QTL from linkage mapping in USNAM and CNNAM, of which 40 overlapped between the two populations (*32*). However, little overlap has been found in QTL between the EUNAM-Dent and EUNAM-Flint populations (*16*). Additionally, cross heterotic pool predictive accuracy in the two EUNAM panels was typically close to zero or even negative in most families (*26*) and Unterseer et al (*34*) found evidence for selection on different pathways between the two panels. This suggests that genetic control for flowering is at least partially differentiated across populations.

The quantitative basis for days to flowering in maize results from the long and complex history of adaptation to new environments. Adaptation to climate was under selection pre-domestication, as the wild progenitor teosinte contended with restricted range during the last glacial period before subsequent expansion as temperatures started to rise 12,000 years ago (*35*, *36*). Altitudinal variation also preadapted maize to range expansion; the direct progenitor, *Z. mays* ssp. *parviglumis*, ranges from 400-1,500m elevation in central Mexico, and the closely related upland teosinte, *Z. mays* ssp. *mexicana* specializes in the higher and drier uplands, 1,600-2,700m (*37*), and has contributed substantial variation to domesticated maize(*38*, *39*).

Domestication of maize took place early during the mid-Holocene maximum (9,000-5,000 BP) (*40*, *41*), where annual temperatures were higher than any time before the industrial era (*36*). After domestication, humans moved maize across the Americas, reaching the southern Andes by at least 3,600 BP (*42*) and moving northward to southern Ontario, Canada by 1,500 BP (*43*). After Spanish contact with the Americas in the late 1,400s, maize was quickly established across the world (*44*–*48*). Modern breeding in the last 100 years has further complicated these histories with the development of inbred lines, heterotic groups, and the intentional introgression of exotic germplasm into global maize, particularly from the historical US Dent germplasm (*49*). The resulting global modern maize germplasm is highly diverse, and days to flowering in inbred lines varies from 35-120 days (*50*).

### Meta-analysis and population structure in human and plant literature

Meta-analysis of GWAS results, or combining p-values from independent studies to increase the significance of common, marginally significant loci, has been routinely applied in human studies since the 1980s, namely because of the problems associated with direct access to human disease data (*51*). Numerous methods exist for combining results (*51*), both Bayesian and Frequentist, but all balance generalizability against power to control for between dataset heterogeneity with respect to study size, phenotypic measurements, or population structure (e.g., MANTRA (*52*)). Human studies tend to be underpowered compared with crop plants due to the inability to replicate phenotypes, and meta-analysis approaches are very effective at increasing power to detect variants in human studies (*1*, *53*, *54*). One of the reasons for these successes is the reasonably high resolution in human studies (*55*), due to the outcrossing nature of human populations and researchers ability to capitalize on this advantage due to the widespread availability of (imputed) high-density markers (*56*). Additionally, the recent and exponential expansion of human populations has generated an overabundance of rare alleles (*57*), which mapping studies have low power to detect alone, but can be boosted to significance in aggregate (*51*).

Historically in crop plants, researchers have focused more on biparental (*14*, *58*, *59*) or more recently Nested Association Mapping (NAM) populations (*60*). This is 59) because, unlike in humans, inbreeding is tolerated and ethics allow for structured mating designs and highly replicated phenotypic estimates. Structured populations require only sparse markers because recombination events are limited, decreasing genotyping costs but also limiting resolution. In return for limited resolution, structured populations control for historical population structure, allowing for accurate effect estimates of even rare variants, which can be a source of novel breeding variation. NAM designs link multiple biparental populations with a common parent, allowing for the estimation of allelic effect estimates across the parents. Most meta-analyses to date in structured populations combine linkage mapping studies rather than GWAS (*8*, *12*). In this analysis, we combined the high resolution of diversity panels, whole-genome SNP coverage, and decreased effects of population structure and accurate effect estimates of structured designs to further resolve the genetic basis for flowering.

The motivating purpose of this study is whether combining populations at whole-genome coverage can enable us to detect more associations, with more resolution, in a complex trait like days to flowering. We reanalyzed five mapping populations for association with flowering traits, days to silking and days to anthesis, using both single marker tests and regional approaches. We introduce a novel method, Boosted Regional Heritability Mapping or BRHM, for regional mapping that controls for population structure and extended linkage disequlibrium in mapping populations and that easily integrates results from populations. Days to flowering is confounded with population structure and adaptation to new environments, so we also tested adaptive relatedness between populations based on the ability of the populations to cross-predict in a genomic prediction framework, and used a Random Forest Classifier machine learning framework to understand the basis for predictions. We test for differences in the mechanisms for adaptation in different environments by comparing GO term enrichment across populations, and testing mapping results against genome annotations and F_ST_ estimates in a machine learning framework.

## Methods

### Datasets

Five publically available maize datasets were reanalyzed for this analysis, four Nested Association Mapping (NAM) designs (US (*60*), Chinese (*27*), and two European panels based on the major European heterotic groups, Flint and Dent (*16*, *26*)) and one diversity panel (Ames) (*11*) (Table 1). Phenotypes used were genotypic estimates reported in the original studies; spatially corrected BLUPs in the Ames, CNNAM, and USNAM populations and spatially corrected BLUEs from testcrosses between the EUNAM-Flint and EUNAM-Dent panels. All of the populations were genotyped at low density in their original study, which we used to anchor whole genome projection for all individuals. The EUNAM panels were genotyped using an Illumina MaizeSNP50 BeadChip (*61*), and the other panels were genotyped with Genotyping-By-Sequencing (GBS) (*62*). Whole genome genotypes from maize Hapmap3.2.1 (*63*) were imputed using K-Nearest Neighbor imputation (KNNi) (*64*) with an overall accuracy of 0.988 and a minor allele imputation accuracy of 0.94 for imputed genotypes, then haplotypes projected onto all populations. Hapmap 3.21(*63*) was called on 1,268 inbred genotypes from across the world, with highly variable depth of coverage, and paralogous sites were retained, as they provide signal in GWAS. Because paralogous sites were retained in Hapmap 3.21, we used KNNi (*64*) to impute, which was robust to high error rates in genotype calling, but KNNi over-imputes missing data to the major allele.

GBS-genotyped populations were projected using FILLIN (*65*), and EUNAM was projected using a custom implementation of FSFHap (*65*). KNNi, FILLIN, and FSFHap all use implementations in TASSEL (*66*). For GBS populations, projection was anchored by 465,085 consensus sites – where the physical positions match and the major/minor alleles are shared – between Hapmap3 and GBS. Projection accuracy as calculated by the correlation between masked known and imputed consensus SNPs was r = 0.99 overall between masked and subsequently imputed genotypes (0.96 for minor alleles). For EUNAM, the parental haplotype breakpoints were imputed for each of the progeny using FSFHap. TASSEL used those breakpoints to project the Hapmap 3.21 genotypes of the parents onto the progeny. While most of the parents of these NAM populations were completely inbred, there were a minority that had residual heterozygosity, which can produce families with three haplotypes segregating. Projected datasets were then filtered using appropriate parameters for the family structure of each population, to ensure a minimum of at least 10 minor alleles for any given site in a population; NAM populations were filtered so that one family must have a minimum minor allele frequency of 0.1 (which controls for the parental residual heterozygosity), giving a minimum minor allele frequency of 0.02 for the population as a whole, and Ames was filtered for minimum MAF of 0.015. Any sites with a maximum heterozygosity above 0.02 or coverage below 0.3 were removed. Before calculating kinships, any residual missing genotypes were assigned a homozygous genotype randomly drawn from the genotypic frequency distribution at each site, by family if appropriate.

**Table 1.** Population statistics for KNN imputed-projected and filtered genotypes

| Population | Pop Type<br>(N families) | N Filt KNNi<br>Sites | Ntaxa<br>DTA | Ntaxa<br>DTS |
| --- | --- | --- | --- | --- |
| USNAM | NAM (25) | 39,302,149 | 4,758 | 4,758 |
| Ames | Diversity | 34,967,601 | 2,312 | 2,279 |
| CNNAM | NAM (11) | 27,047,718 | 1,928 | 1,928 |
| EUNAM-Dent | NAM (10) | 23,766,799 | 841 | 841 |
| EUNAM-Flint | NAM (11) | 25,388,592 | 811 | 811 |

### Multidimensional Scaling (MDS) of American landraces and NAM parents

It is often unclear how inbred lines fit within an adaptive evolutionary context, since breeding programs cross and select on progeny from unrelated individuals. We performed joint MDS analysis with American landraces from Takuno et al (*67*) spanning the two American temperate-tropical gradients with the parents of the four NAM populations to understand the distributions of the parents across American temperate adaptation. Each landrace individual was included ten times in the IBS distance matrix used to calculate the MDS coordinates so that the first two coordinates reflect the relatedness between landrace individuals. Landraces and parents for CNNAM were natively genotyped using GBS, and EUNAM and USNAM projected in Hapmap3.21 coordinates. MDS was based on 465,085 consensus sites between Hapmap3 and GBS (cmdscale () in R, using an IBS distance matrix generated in TASSEL)

### Cross population prediction

Cross population prediction were performed to better understand how populations were related to each other with respect to shared genetic architecture for days to flowering. Cross population prediction was performed using GBLUP as implemented in TASSEL (*66*). The training and test populations were combined in a single kinship (similarity) matrix, calculated using the Centered-IBS implementation in TASSEL (Endelman and Jannink (*68*), after Van Raden (*69*)) and phenotypes for the test population masked so that the model was trained solely from the training population phenotypes. This method scales such that the mean diagonal elements equal 1Ȼf, accommodating the divergence from Hardy-Weinberg equilibrium inherent in structured populations. Predictions from the resulting model for the test phenotypes were then correlated with the true phenotypes (the “predictive ability”) in R using the Pearson method in cor.test() in the stats package. We use predictive ability, rather than prediction accuracy, the predictive ability divided by the square of the heritability, because heritabilities for flowering traits are quite high (all greater than 0.8) and it is more conservative. Predictive ability based on genomic subsets presented in Figure S6 – coding sequence, three and five prime regions, introns and intergenic – were calculated following Rodgers-Melnick et al (*70*).

### Genome-wide Association Study (GWAS)

To determine associations between individual SNPs and flowering traits a GLM was conducted in TASSEL (*66*) on two sets of phenotypes. One was the unmodified phenotypes from the original publications, uncontrolled for population structure, because flowering is not only correlated with population structure, but it acts to differentiate populations. Thus, many of the regions that differentiate populations may also control regions important for temperate adaptation and we captured these in the uncorrected model. We also controlled for population structure by fitting 5 MDS coordinates calculated from an IBS distance matrix in TASSEL for the Ames panel, or fitting a family term for the structured populations in a TASSEL GLM for the NAM populations. We additionally tested a two-step modified mixed linear model framework for computational efficiency. We first calculated residuals in the TASSEL MLM from models incorporating 1) only a kinship matrix calculated in TASSEL using the Centered-IBS method (*68*, *69*) as a random effect, or 2) both a kinship matrix as random and 5 MDS coordinates (for Ames) or a family term (for NAM populations) as fixed effects. The residuals from these models were used as the response in a TASSEL GLM. We do not include the two-step GLM results because we found that, due to the high correlation between population structure and flowering time, fully controlling for population structure reduced power to detect even well-known flowering loci such as Vgt1 or ZmCCT.

### Resample Marker Inclusion Probability (RMIP)

The RMIP (*71*) approach is a resampling model selection procedure that can help identify the most informative SNPs. RMIP was first applied to maize to analyze leaf architecture traits by calculating residuals for each chromosome from a model that included terms from a joint linkage model for all other chromosomes, following Tian et al. (*3*); this was the method used to calculate residuals for Ames and USNAM. For CNNAM and the EUNAMs, instead of using joint linkage, the residuals were calculated from a mixed model fit with a leave-one-out relationship matrix for each chromosome, which was based on the markers from all other chromosomes except the target chromosome. The residuals were then used to fit a stepwise model to 100 random subsamples of 80% of the marker data. The RMIP for a SNP was set to the number of times it appeared in any model divided by 100. Tests found that RMIP is insensitive to rare alleles, and only works well when an allele is present in at least three families; because of the limited number of parents in the EUNAM and CNNAM populations, we found that this approach had limited power to identify SNPs and did not focus on the RMIP results in subsequent analyses (but see Figures S9–S18 for RMIP results in context of the other mapping methods).

### Regional Heritability Mapping (RHM)

Regional genomic mapping approaches should be better at detecting globally rare alleles present in parents of mapping populations, as those alleles are brought up in frequency in the progeny. Because of the variability in population structure and consequently MAF and SNP overlap between the populations, we first used a regional variance partitioning approach to identify regions that could be jointly analyzed between the populations. We follow a similar approach to that introduced in Nagamine et al (*2*), testing each region of the genome using a model with two kinship matrices, one derived from the region of interest and the other from the rest of the genome, fit as random effects. Additionally, family is fit as a fixed effect for the NAM populations (Figure 1). Our implementation differed from the Nagamine et al method in that the genomic kinship was not constructed using markers from the region tested. The same model is also fit without the regional kinship, and the likelihood of the reduced model is divided by the likelihood of the full model to produce a likelihood ratio test statistic for the region. Following Nagamine et al, the test statistic is compared against a mixed chi-squared (0,1) distribution for significance. For this study, we used 10,024-site kinships, because preliminary tests indicated that this was the smallest number of SNPs to generate reasonably stable variance estimates in this model, with respect to convergence. Before metaanalysis of RHM significant regions across populations, the resulting p-values are adjusted for multiple testing using the Benjamini-Hochberg method in the p.adjust method in the R package “stats”.

Preliminary tests found the RHM mapping approach was only successful in identifying associations in the larger populations (Figures S1–2). While this method has been successful in humans, and located numerous regions of the genome in the Ames diversity panel, and, oddly, reported more regions for DTS in the large USNAM panel than the original study (*5*), very few regions were identified in the smaller NAM designs, and meta-analysis did not identify new regions of the genome. These results suggest that RHM is especially sensitive to population size. Additionally, reduced recombination inherent to especially the EUNAM designs (*72*) is theoretically problematic for RHM. In RHM, if LD extends beyond the target region, even if the causal signal is located in the region, including the region may not increase the likelihood of the model since the signal is also represented in the “rest of the genome” kinship. These observations stimulated development of the BRHM method, and RHM results were not used in further analyses.

**Figure 1.**
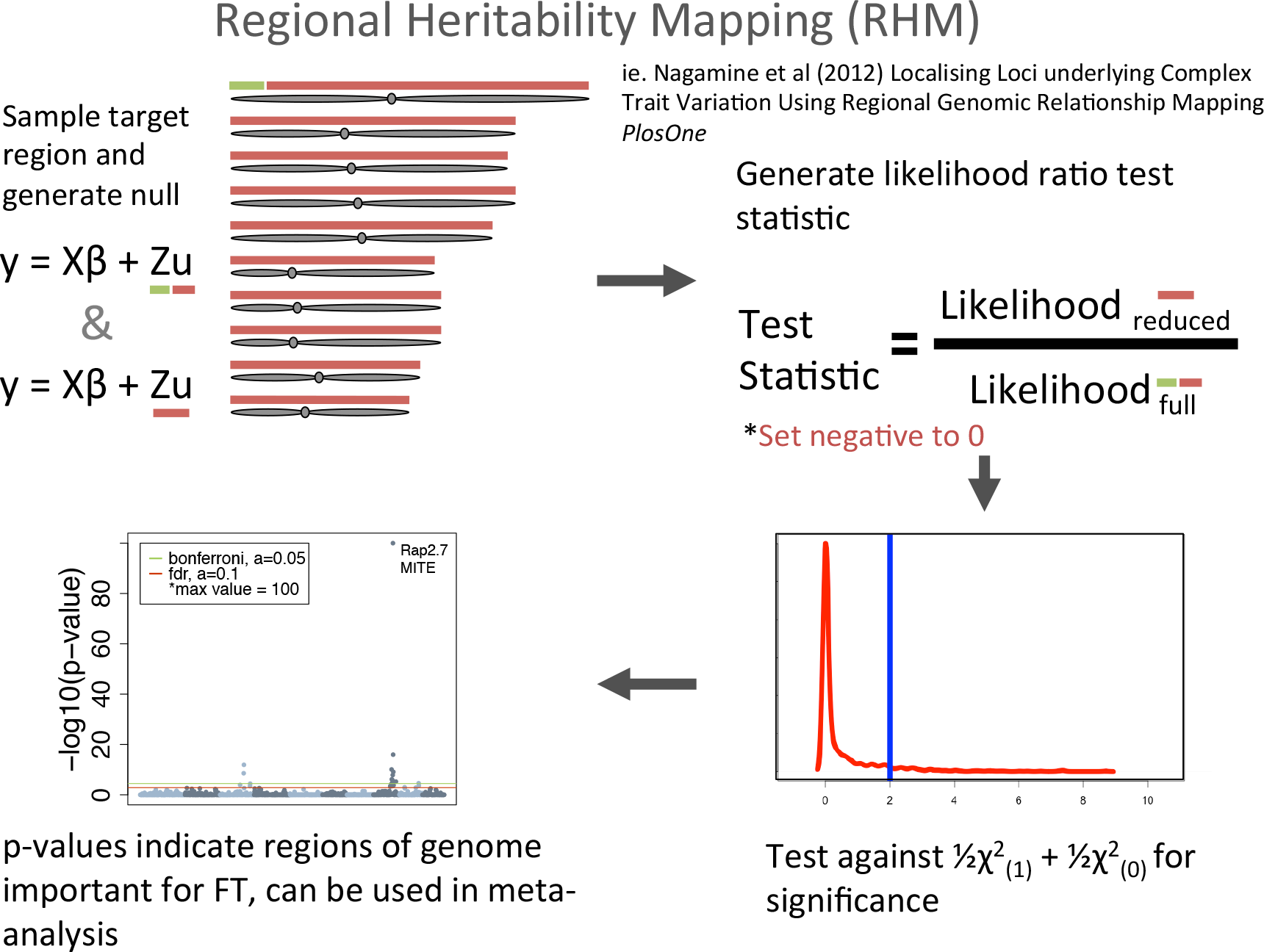
Regional Heritability Mapping (RHM)

### Boosted Regional Heritability Mapping (BRHM)

Because the rest of the genome kinship can capture signal from the target region when LD is high, as is common in NAM designs, we developed an extension of RHM, BRHM, to increase power to detect associations, especially in the smaller NAM panels. The BRHM algorithm iteratively samples a scaled number of randomly placed regions on the target chromosome as random effects, controlling for the rest of the genome, in a mixed linear model framework using the REML solver implemented in LDAK against response phenotypes (Figure 2) (*73*). Each iteration randomizes the distance between each of the regions in a model, and thus the degree of LD shared by any two kinship matrices, generating a distribution of estimated heritabilities by any given region in the genome. We used 30,000 site kinships computed using the “Centered-IBS”(*68*, *69*) method for structured populations or “Normalized-IBS”(*74*) for the Ames diversity panel as implemented in TASSEL(*62*). The value thirty thousand was determined empirically to maximize resolution while maximizing model convergence. The number of chromosomal kinships was scaled for one kinship per 50Mbp, to ensure a consistent probability that any two kinships would be in close physical proximity. For each iteration of the model, kinships were established at random start sites (in increments of 512 site blocks for computational efficiency in the final models), using a uniform random generator, with the caveat that they cannot overlap and any kinship established less than 30,000 sites from the end of the chromosome was adjusted to 30,000 sites before the end.

BRHM determines significance for regions by comparing the resulting distribution to an empirical null distribution of estimated heritabilities, representing the signal for the region resulting from global population structure. The null heritabilities are estimated using the same genomic kinships and models, but replace the true phenotypes with a null phenotype as the response. The null phenotypes randomly scramble the residuals from the model *y* = *Xβ* + *Zv*, where the random effects include the kinship calculated from the untested chromosomes and the error term, by adding the error estimate from another randomly chosen individual to the true BLUP estimate for the kinship for each taxon, or genotype. This generates a null to test the hypothesis that the heritability estimated by the target kinships from the chromosome tested are identical to that expected from population structure as detected from the rest of the genome. Models were run with random starts for all kinships for the number of iterations expected to cover the genome to 40X coverage; after these iterations, the start site of the first kinship assigned to each model was chosen to ensure that a minimum of 40 models covered each SNP.

Paired real and null results were aggregated into regions, 512 site regions in the final model, based on unique model coverage. Models that did not converge (ran more than 25 iterations, or had a difference in the model likelihoods greater than 0.1 between the last 2 iterations) were excluded, rendering a small number of regions with less than coverage. Because the resulting distributions of null and estimated results were not always normally distributed, significance for each region was evaluated using the non-parametric Wilcoxon signed rank paired two-sample one-sided test implemented in the R method “wilcox.test()”. The null hypothesis for this test is that the median of the population of estimated results is not greater than the median of the population of null results. To control for differences in power across regions, model results were randomly downsampled to 30 models per test, and the average p-value for thirty tests reported. If the distributions were identical – usually in the case that the null and estimated values were all zero – the p-value was set to 1. Before meta-analysis, the resulting p-values are adjusted for multiple testing using the Bonferroni method in the p.adjust() method in the R package “stats”.

**Figure 2.**
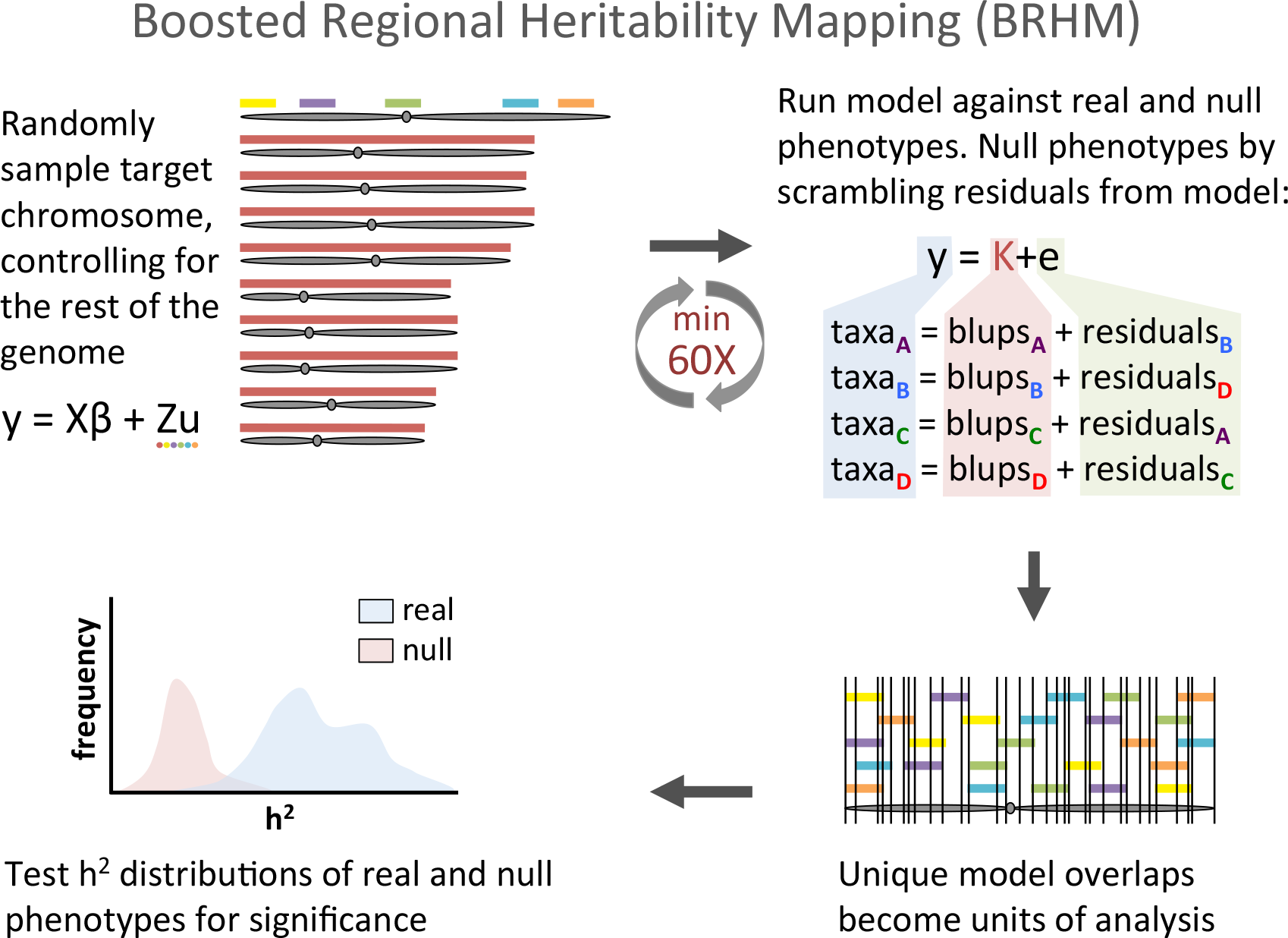
Boosted Regional Heritability Mapping (BRHM)

### Meta-analysis for variance component methods

We combine p-values for regions of the genome with unique region overlap across populations using Fisher's method for combining p-values (*75*), using the formula, 
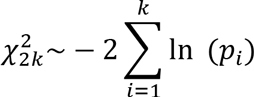
where p_*i*_ is the p-value for *i*th hypothesis in 1…*k* populations. The resulting test statistics are *χ*^2^_(2k)_ distributed, and the mean false discovery rate is calculated as α = (k+1)/2k. While this method is not appropriate for combining GWAS results because it does not account for the direction of effects or heterogeneity between populations, this is not a constraint in variance mapping approaches because p-values are generated against internal controls and heritability is unidirectional. Meta-analysis for resampling GWAS entails directly combining results from each population; a meta-analysis for GWAS results was not conducted because of limited shared SNPs and because initial tests indicated that the structure of the populations and extended LD in the NAMs generated no positive associations for the *Vgt1* locus.

### Machine learning analyses with GWAS results as the response

We employ the RandomForestClassifier method in Spark (*76*) in a Databricks environment, classifying additive p-values from GWAS results such that the smallest 1% are classified as 1, or true positives, the next 4% of results are skipped, and 2 million random results from the remaining 95% are classed as 0, or true negatives. We employ a classifier rather than a regression based approach, because we are only interested in how the top regions rank relative to all the others, and a regression approach gives more weight to the insignificant results because they vastly outnumber the top hits. P-values for SNPs not reaching filtering criteria in a given dataset were assigned to 1, before ranking results, because a SNP that does not meet MAF filtering criteria will not be highly significant, even if biologically causative. We cross-validated by using the other nine chromosomes to predict the tenth. Accuracy is reported as area under the curve (AUC), the integration of the ROC curve (for Receiver Operating Characteristic). ROC curves display the momentary false positive rate on the x-axis against the true positive rate for each unique raw predicted value reported by the classifier model, as the prediction for each classified instance is reported on a continuous random variable. An AUC of 0.5 is equivalent to guessing. Overall AUC is simply the average AUC for the 10 chromosomal tests. Top predictors were reported as the average mean ranking across all 10 tests. Within a given test, predictors are ranked by which splits they participate in for a given tree, averaged across all trees.

### Combined GWAS

Genotypes and phenotypes were directly combined together for combined GWAS analysis to test the efficacy of directly testing phenotypes derived from non-overlapping research designs together. Genotypes were only minimally filtered for polymorphic sites, leaving 70 million genotypes in the dataset. To control for differences in heat units across different studies, phenotypic variances for each population were shrunk using a mixed linear model (the lmer package in R) where dataset was fit as a random effect. This shrinks variances due to the different environments in which each populations were grown but also has the less desirable effect of standardizing true phenotypic differences between the populations. Shifting the mean of each individual by the difference between the value predicted for B73 for the appropriate dataset and the value for B73 predicted by the Ames diversity panel centers the resulting phenotypes. All populations were run as a single analysis in a GLM framework using the FixedEffectsLMPlugin in TASSEL, controlling for population structure using a dataset term and 5 MDS coordinates calculated from an IBS distance matrix in TASSEL(*66*).

### Candidate gene lists

We extend the time to flowering candidate gene list from Dong et al(*77*), adding all of the genes from the CENTRORADIALIS (ZCN) gene family, after Danilevskya et al (*78*) (Table S1), as the 79 candidate gene list. We additionally use unmodified the 918 candidate gene list from Li et al (*32*). We also generate five new lists of candidate genes, one from the combined GWAS DTA results and two each from Ames GWAS results, with and without population structure correction for DTS and DTA, based on adding unique genes from a 100kbp window around top GWAS results sorted by p-value (Table S2). The Ames candidate lists, while confounded with autonomous flowering pathway genes, should be more enriched for temperate adaptation alleles than the physiological candidate lists.

### Machine learning analyses for candidate genes

We also use Random Forest classifiers to provide insight into the attributes of variants that explain top gene regions based on candidate genes for days to flowering and GWAS results from this study. SNPs from within candidate gene regions and 100kbp on either side are set as class 1, and set the rest to class 0. Tests are done the same way as with GWAS as the response. Mapping results, per-site F_ST_ estimates and power considerations such as MAF and coverage for Hapmap 3.21 SNPs provide machine learning variables (the total list of predictors used in models is provided in Table S3). F_ST_ was calculated on a per-site basis between populations in vcftools (*79*), missing and negative values set to 0. Additional F_ST_ calculations were derived from sets of thirty Hapmap 3.21 taxa representing temperate US germplasm, tropical germplasm and Northern Flint germplasm (Table S4).

### GO term comparison between populations

To better understand which classes of gene function are enriched in the BRHM mapping results for the five populations, we calculated enrichment across the top 1,000 significant genes. The top genes were chosen by iterating through the BHRM results ordered by the downsampled Wilcoxon-signed rank p-values for each population, adding genes from these regions until 1,000 genes were returned so that the genes surveyed for each population were balanced. The top 1,000 gene list was submitted to AgriGO (*80*) for singular enrichment analysis (SEA) against the complete GO database using Fisher's test for significance, no FDR correction, and a minimum of 5 mapping entries, using the Zea mays AGPv3.30 reference. The resulting GO terms were aggregated by the authors based on GO definitions (Table S5). A chi-square goodness of fit was calculated in Excel (CHITEST) from the sum for each category.

## Results

This study combines five populations, four NAM designs and a diversity panel, totaling over ten thousand individuals (Table 1). These populations were previously genotyped using GBS or an Illumina array and imputed with whole genome resequencing using FILLIN and FSFHap, respectively. This resulted in 70 million segregating markers across all populations (Table 1). The populations have variable genetic overlap based on the first two MDS coordinates, where Ames, as expected of a diversity panel, shows the greatest genetic diversity and CNNAM and EUNAM-Flint are the most isolated (Figure 3). Only half of the SNPs that survive filtering are shared between study populations (Figure S3), and the allele frequencies at those loci are highly variable (Figure S4).

### Evaluation of relatedness between populations by MDS analysis and phenotypic comparison

**Figure 3.**
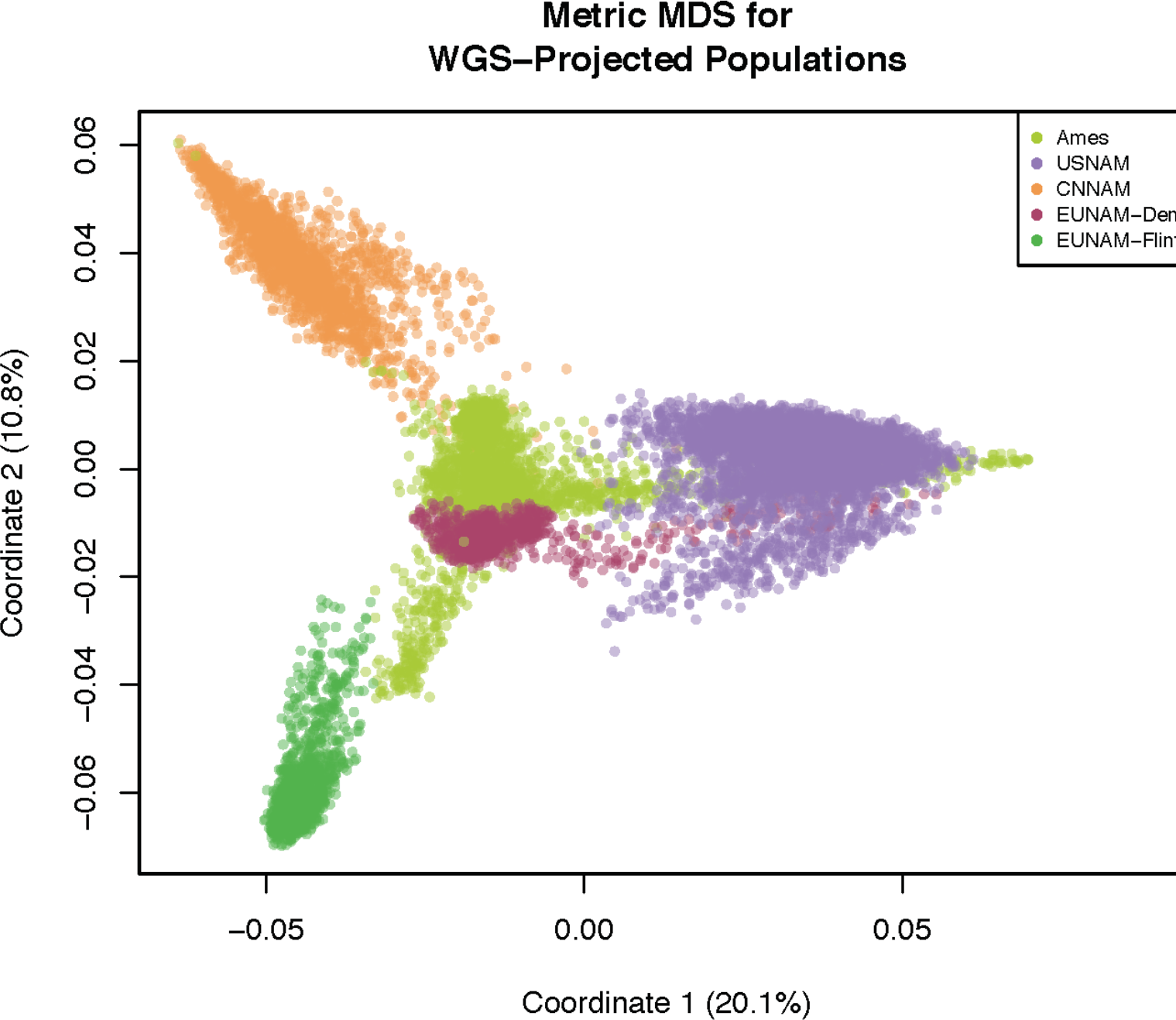
MDS of study populations from 70 million projected Hapmap 3.21 segregating SNPs (cmdscale () in R, using an IBS distance matrix generated in TASSEL).

To better understand the global genetic relationships between the founders of the four NAM populations, Figure 4 shows the genetic relatedness of the NAM-type population parents to a panel of GBS-genotyped landraces from across the Americas (Ames includes all of the USNAM parents). In this context, the NAM founders overwhelmingly cluster on the North American temperate-tropical gradient, rather than the South American gradient. CNNAM and EUNAM-Dent panel show a more restricted geographic origin for the founder germplasm, while the other populations contain parents with a greater mix of American tropical and temperate origins.

**Figure 4.**
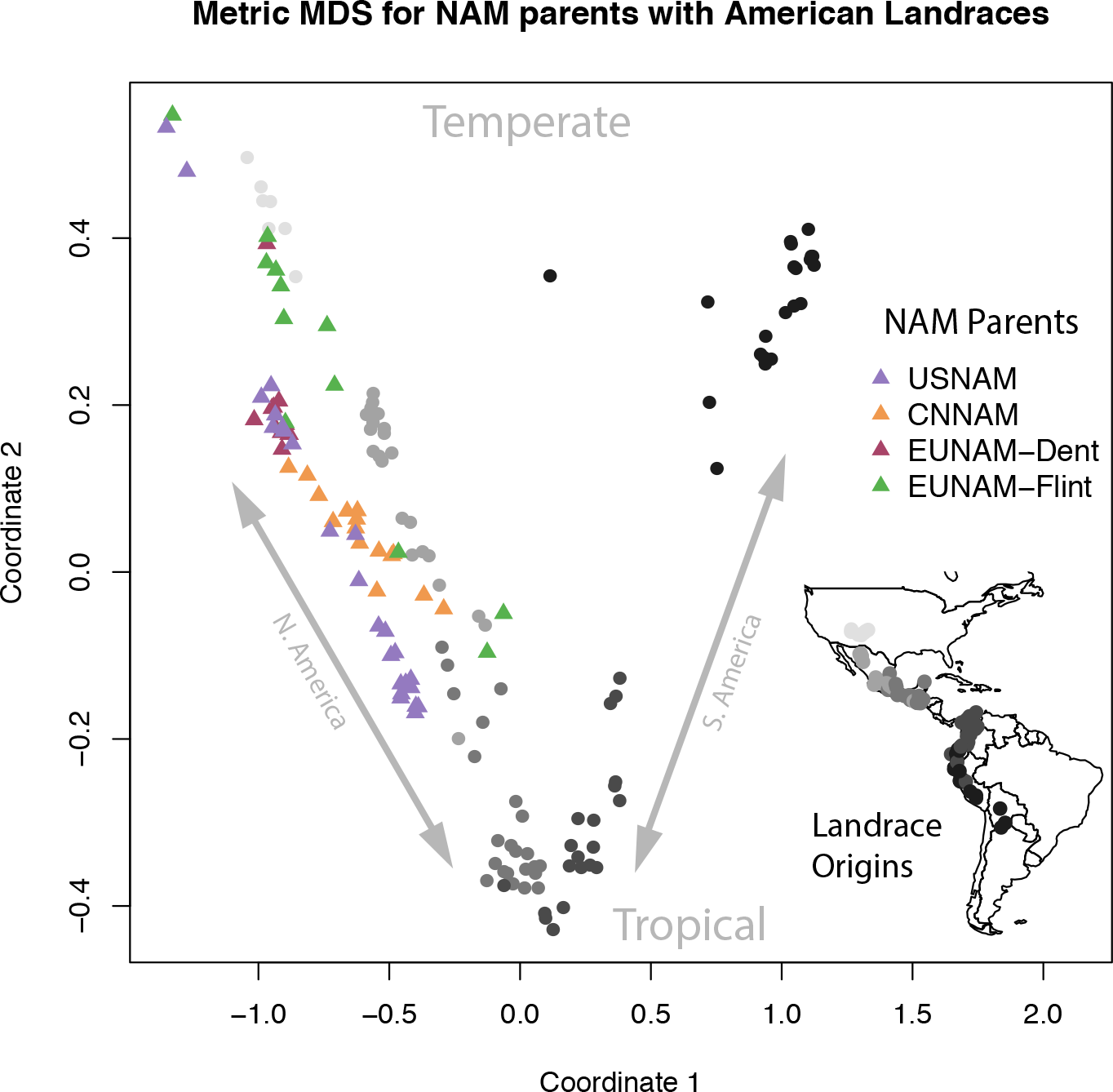
MDS of GBS genotypes American landraces from Takuno et al (*67*), replicated 10X each to drive the first two coordinates, and NAM population parents. The closest US-NAM parents to EUNAM-Dent are the temperate US Dents, including B73, B97, Oh7B, Ky21, and MS71, while the nearest US inbreds to CNNAM parents are South African inbreds M162W, M37W, and the Texas US line, Tx303. All of these are more proximal to the tropical Mexican landraces. The EUNAM-Flint recurrent parent, the Iodent/BSSS EUNAM-Dent line UH007, clusters with the majority of the EUNAM-Flint parents. The Spanish lines EP44, EZ5 and the French line F64 are the most tropically associated lines in the EUNAM-Dent population.

The five populations vary with respect to both the mean and the total variance for flowering (Figure 5). This is the result of both genetic variance within the populations, and environmental effects of the growing regions. Shifting the populations by the difference between the predicted value of B73 in the population relative to the measured value of B73 in the Ames population allows for comparison of the days to flowering of the different populations in the same environmental context. The EUNAM populations, which were grown in exclusively European locations, shift to the short flowering end of the distribution, and CNNAM shifts to a later flowering average relative to Ames. Problematically, the EUNAM-Dent population shifts to the earliest flowering position, earlier than EUNAM-Flint, which flowered slightly earlier in similar European trial locations. This is likely due the increased relatedness between the EUNAM-Dent and B73, relative to EUNAM-Flint, giving a better estimate of flowering. Lehermeier et al. (26) ind that the Flint panel has low predictive ability for the B73 testcross in leave-one-out cross validation.

**Figure 5.**
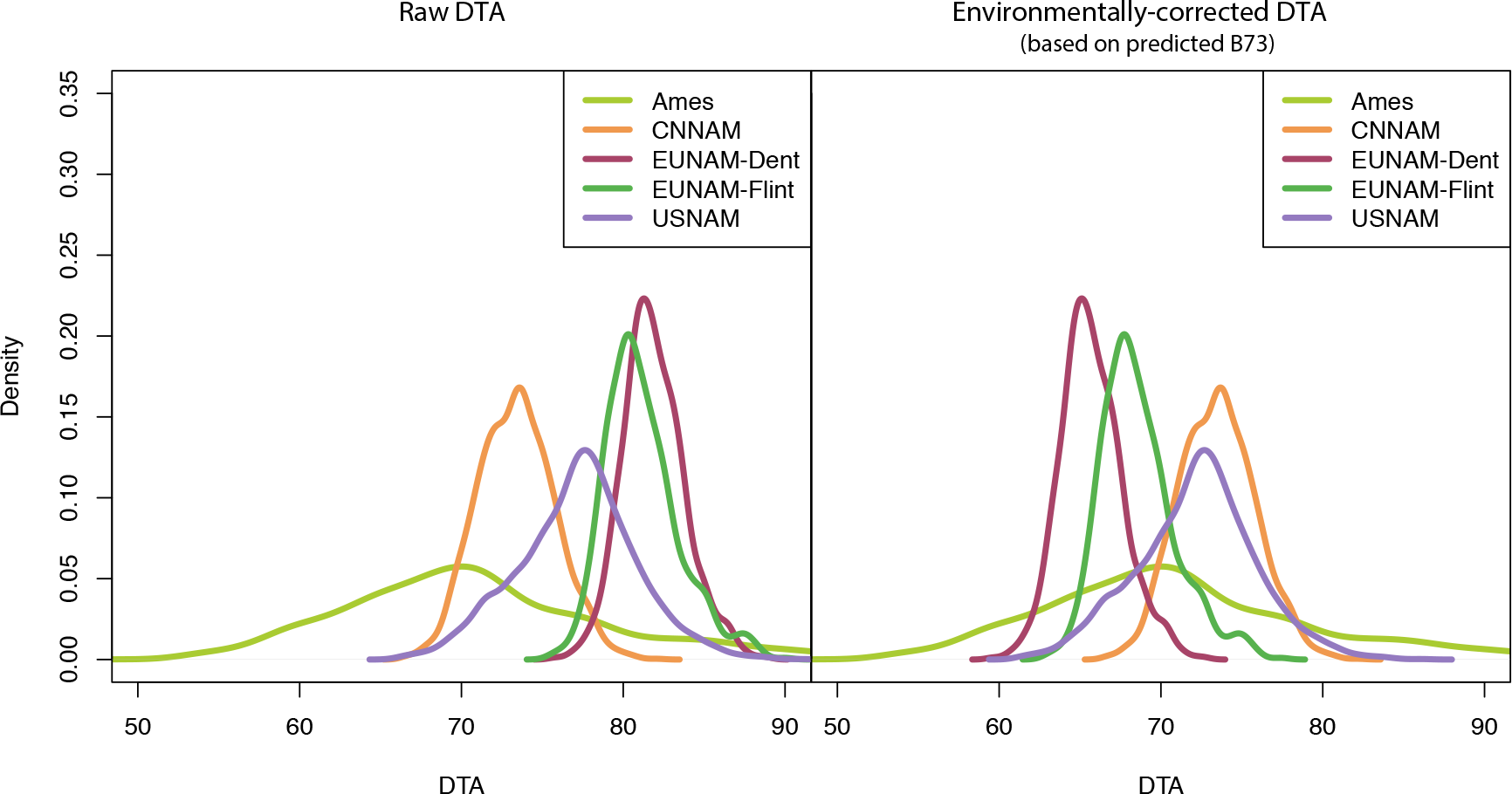
Distribution of reported spatially-corrected phenotypes for days to anthesis, before and after B73-centered correction for dataset.

### Cross population prediction of days to flowering

We performed cross-population prediction to better understand how the genetic basis for flowering is structured across these populations. Cross-population predictions were generated pairwise with GBLUP in TASSEL across all genomic SNPs (Figure 6). Ames has the best cross population predictive abilities (given throughout as the Pearson correlation between observed and predicted, conservatively) overall, and unsurprisingly given the close relationship between Ames and USNAM, the best predictive ability is when the model is trained on Ames and predicting DTA in USNAM (r = 0.78; 0.67 for the reverse). Ames predicted all of the populations better than the others, with the exceptions of CNNAM, which might be expected since Ames is a diversity panel and suggests that Ames is a superset to everything but CNNAM (Chinese lines are present, but poorly represented in Ames (*11*)). CNNAM, uniquely, not only was predicted poorly by the other population but also did not predict any population well. The EUNAM-Dent population also had poor cross-population predictive ability, but was well predicted by Ames. Both of these populations also show a marked non-linear correlation in predicting Ames, suggesting that a large subset of the Ames population is not represented in these panels. The EUNAM-Flint population is well predicted by Ames and USNAM, but especially when predicted by USNAM, the high overall correlation is especially driven by a relatively small set of lines in the top right of the plot descended from primarily three families descending from the Spanish lines EP44, EZ5 and the French line F64. These three parents are the most “tropical” of the EUNAM-Flint lines on the N. American gradient (Figure 4).

We also looked at predictions using only SNPs significant in mapping, and using SNPs from the functionally annotated regions of the genome, and found that subsets can sometimes improve predictions if genomic predictive ability was low, but never if the accuracy was already high (Figures S5 and S6). An exception where a well predicted population is further improved by a subset is when the EUNAM-Flint population is predicting Ames or USNAM; in both of these cases predictive ability is already high, and genic or open chromatin subsets further increased accuracy.

**Figure 6.**
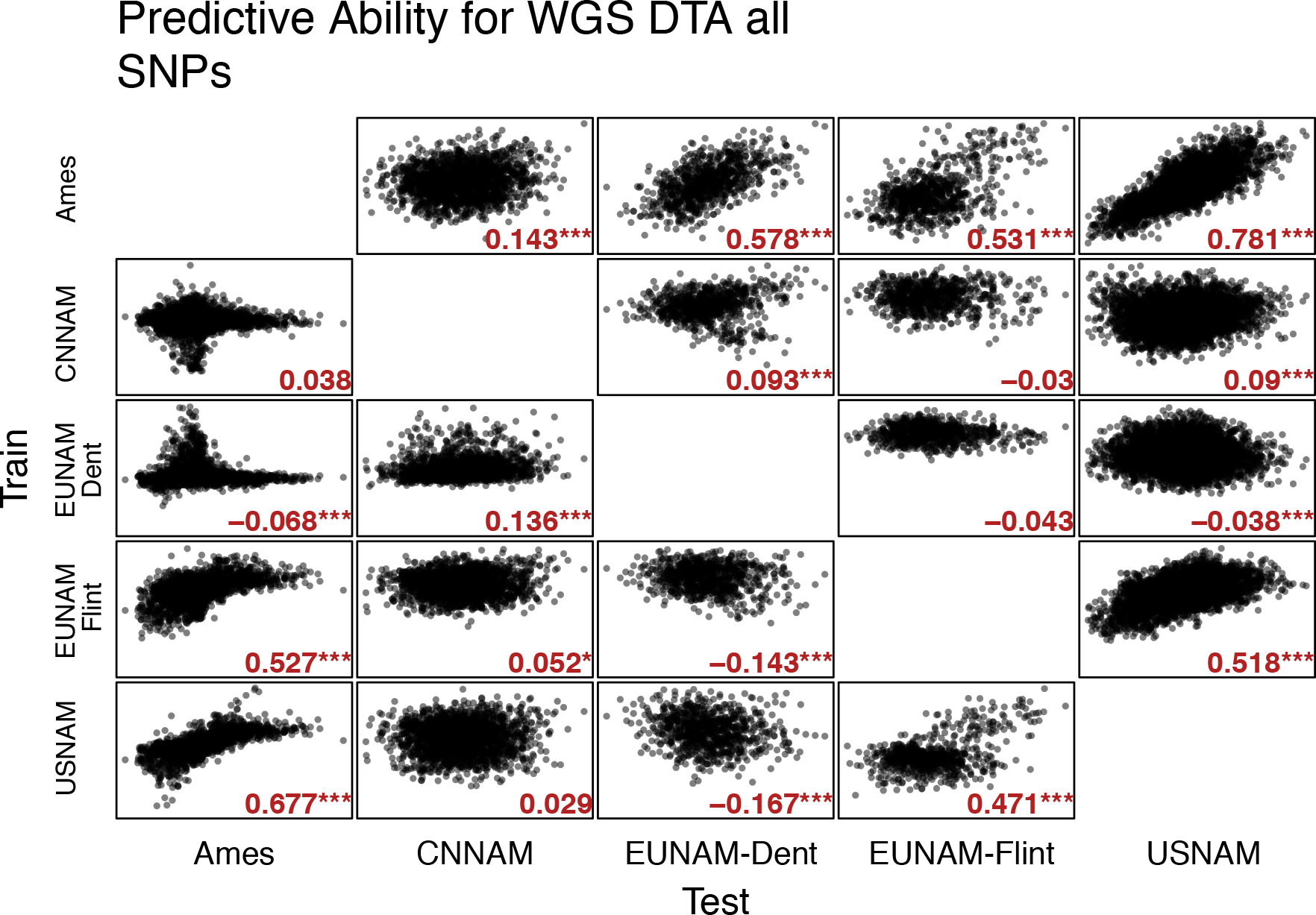
GBLUP cross population predictive abilities for DTA using all 70 million segregating SNPs in Hapmap3.21. Genetic similarity matrix generated using the Centered-IBS method in TASSEL.

Because predictive ability is a function of relatedness, we calculated F_ST_ statistics for all pairs of populations. Figure 7 shows a correlation of −0.41 for F_ST_ and predictive ability for DTA across all markers, confirming a relationship between close relatedness and high predictive ability, but there are outliers that do not fit the expected pattern. Figure 7 shows that CNNAM and EUNAM-Dent generally have lower cross-population predictions than expected by population relatedness, with the exception that EUNAM-Dent is predicted similar to expectation by Ames. In contrast, EUNAM-Flint has higher than expected predictive ability with USNAM.

**Figure 7.**
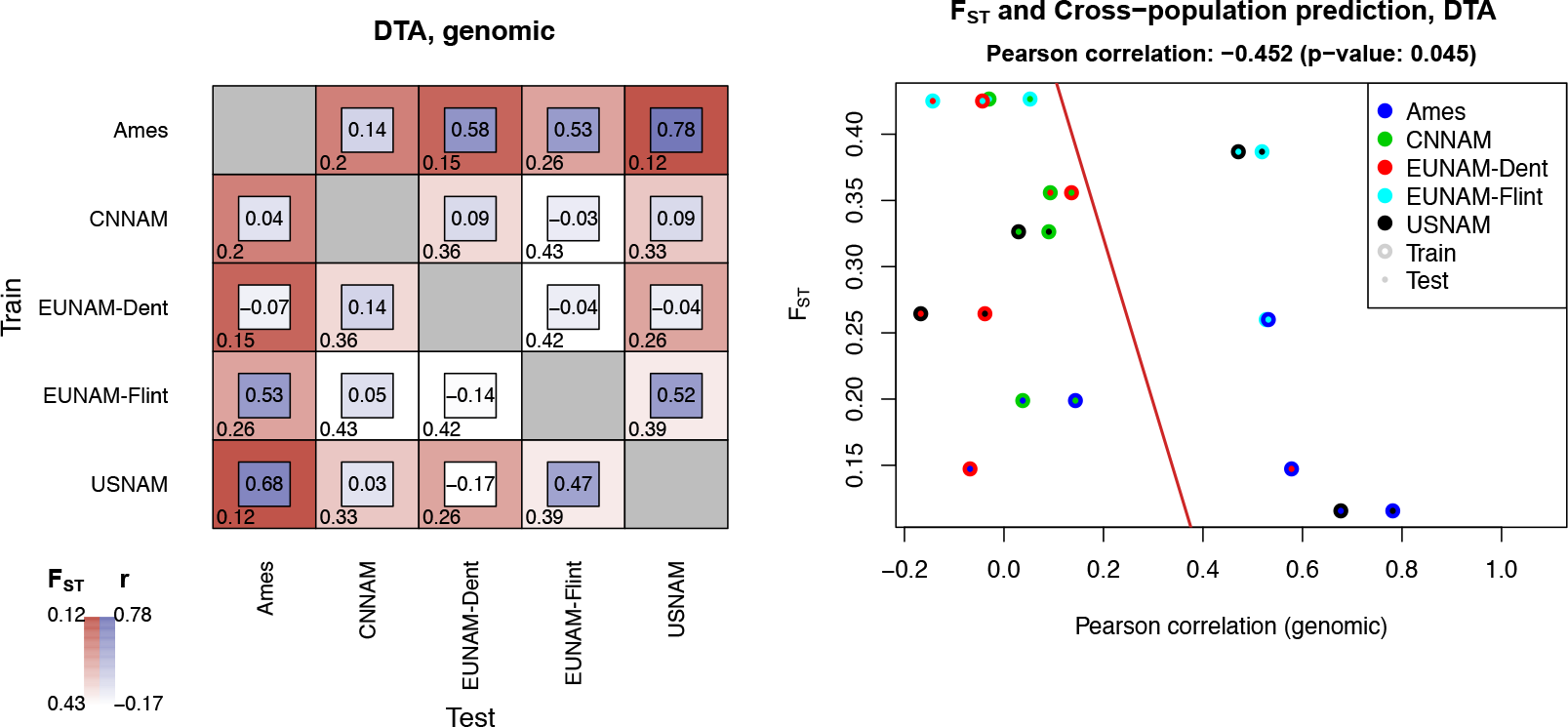
F_ST_ and cross-population predictive ability, DTA.

### Mapping days to flowering loci across the genome by population, and combining results in meta-analysis

We mapped SNPs and regions of the genome for populations individually, comparing overlap of significant mapping results between the populations, and with published candidate gene lists. We performed GWAS and resampling GWAS (RMIP) to identify important SNPs, and two regional variance component approaches, RHM and the novel BRHM extension, so that populations with minimal SNP overlap can be compared on a regional basis. We found the RMIP and RHM approaches to be inappropriate, especially for the smaller NAM populations, and did not pursue further analyses.

GWAS results were run for each population, and because changes in flowering time generate population structure, we performed GWAS with no population structure correction and minimal population structure correction (a family term for NAM populations, and 5 MDS coordinates for Ames). GWAS results generally tracked with major findings from other methods (Figure S7, S9–S18), with the NAM populations tracking large regions of the genome, consistent with lower rates of recombination reducing resolution. Family control in the NAM populations exacerbated this pattern, removing peaks from all but the highest variance regions by BRHM or RHM, especially in the EUNAMs, which have less recombination (Figure S8). In Ames, the only population with resolution for GWAS, population structure control changes the significance of regions, but still identifies regions as significant that BRHM does not.

GWAS results were not comparable across populations in meta-analysis due to lack of shared segregating SNPs (Figures S3 and S4). As expected, the NAM population datasets did not gain resolution on their own from marker-level testing, due to high local LD resulting from minimal recombination inherent to the NAM designs (Figure S7). To assess overlapping significant loci, despite lack of overlap of SNPs between filtered sets in each population, we used a random forest classifier machine learning approach, where additive p-values for excluded SNPs for a given population were imputed to 1 (Figures 8, S19). Without population structure control the results are similar to the cross-population predictions, Ames and USNAM are well predicted, mostly by each other. EUNAM-Flint is predicted to a lesser extent by Ames, followed by USNAM, and EUNAM-Dent is predicted equally well, but by EUNAM-Flint. CNNAM is predicted less well than by chance. After accounting for population structure, no population is well predicted by the others, and only Ames has a positive predictability.

**Figure 8.**
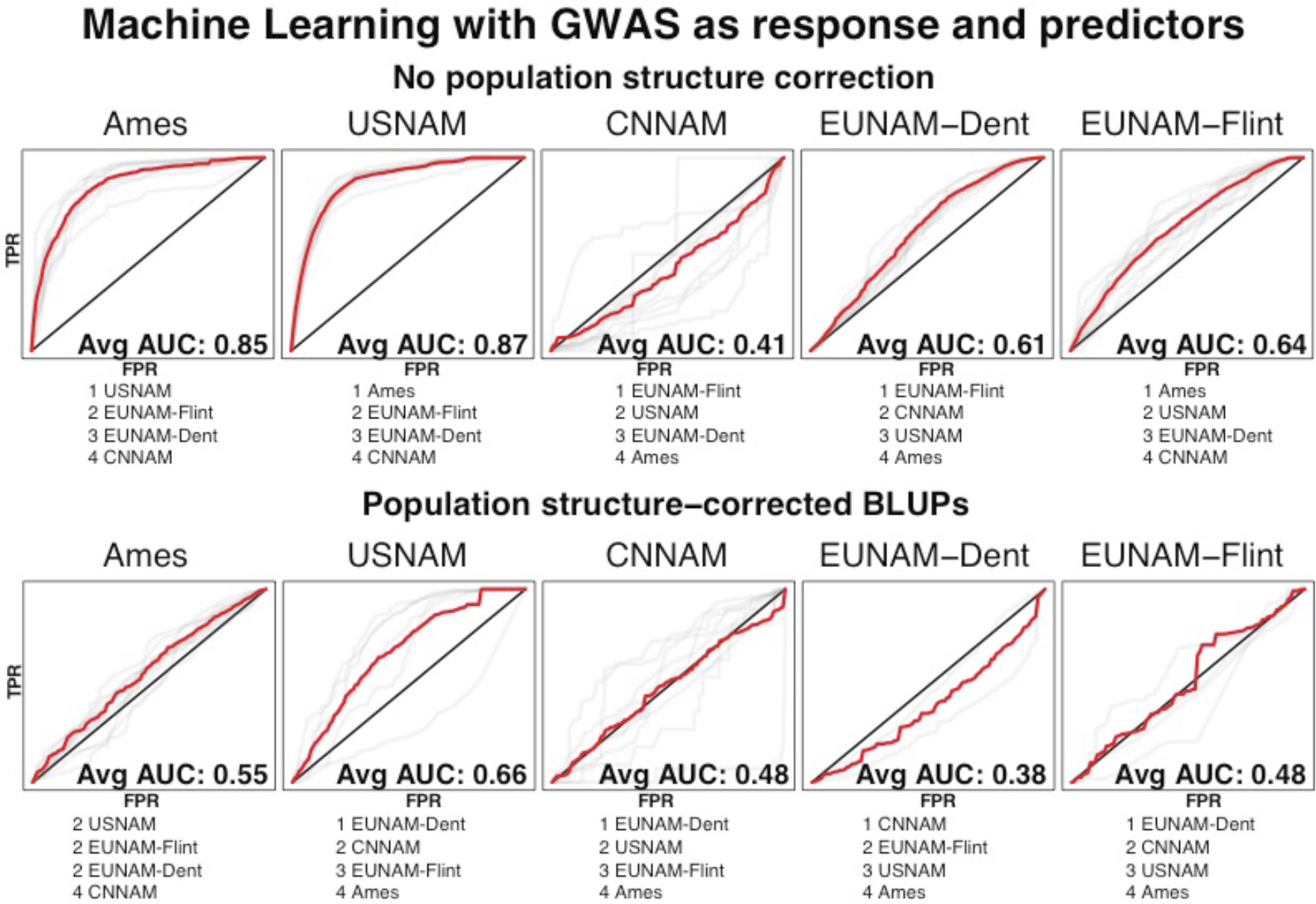
Average AUC (in bold) and predictor rankings across all chromosomes from random forest classifier for GWAS results between populations to evaluate overlap in GWAS results between populations. The red line represents the mean of the chromosomal ROC curves, sampled at 0.01 intervals. Predictors are equivalent GWAS (with or without population structure correction) for the other populations. Trained on nine chromosomes and tested on the 10^th^. Not all chromosomes for the population structure corrected results have top predictors, and are excluded from overall AUC calculations.

Because the GWAS results contain signal from population structure as well as from underlying flowering loci, we also tested the GWAS results against measures of population differentiation between three different N. American germplasm pools, tropical, temperate, and Northern Flint; the temperate germplasm results from the admixture between the southern Dent and N. Flint pools (Figures 9, S20). Confirming the inference from the cross population predictions, Ames is predicted very well (overall AUC of 0.89) by the tropical contrasts, followed closely by USNAM, with EUNAM-Flint well predicted at 0.67, EUNAM-Dent rather poorly predicted at 0.59 and CNNAM slightly negatively predicted with an AUC score of 0.47. Population structure control with a family term for the NAMs and 5 MDS coordinates for Ames reduced predictive ability by population differentiation, but did not eliminate the effects, confirming that flowering time is closely tied to population structure.

**Figure 9.**
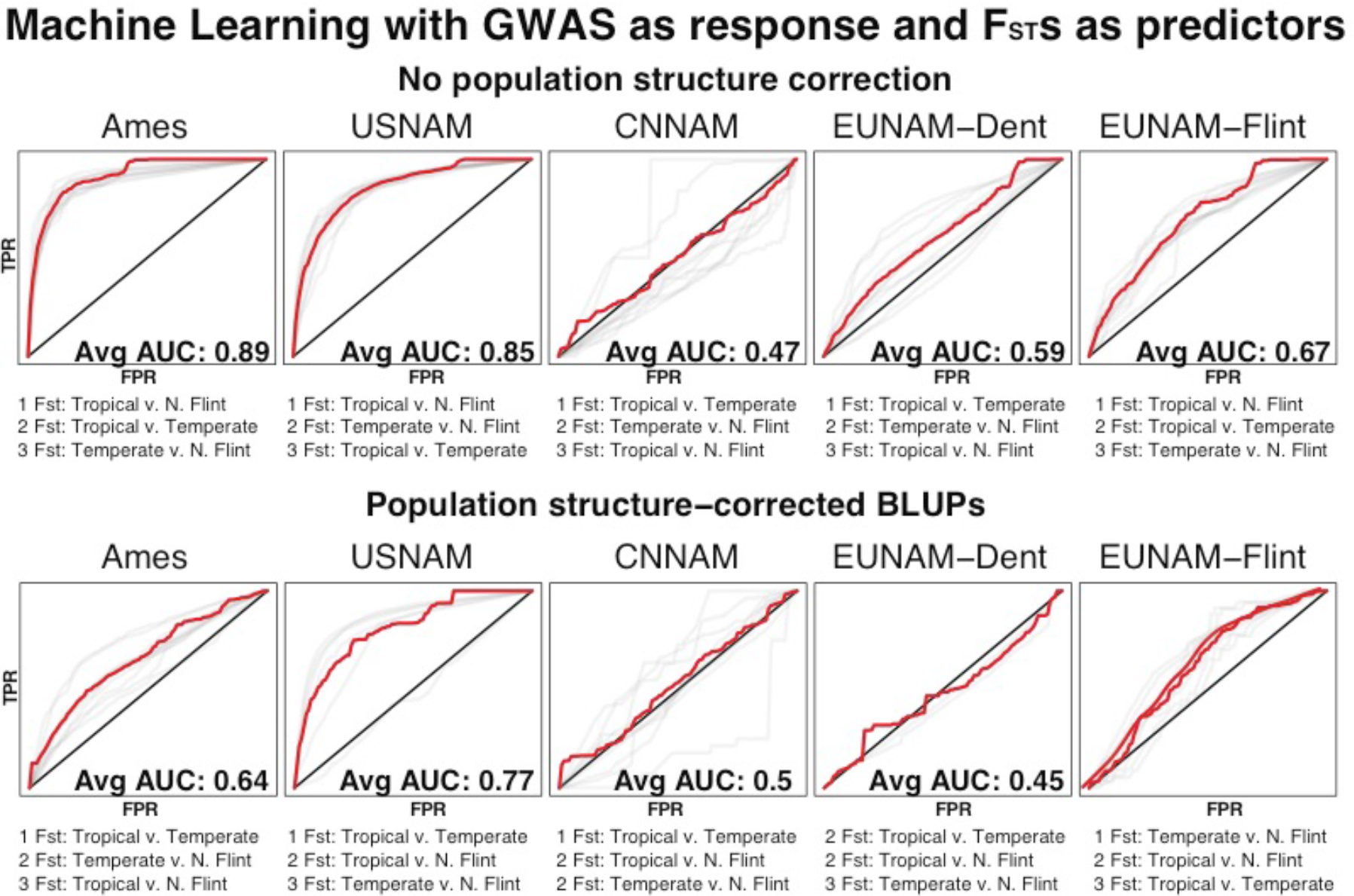
Average AUC and predictor rankings across all chromosomes from random forest classifier with GWAS additive p-values as the response, and N. American F_ST_s as the predictors. Trained on nine chromosomes and tested on the 10^th^. The red line represents the mean of the chromosomal ROC curves, sampled at 0.01 intervals. Not all chromosomes for the population structure corrected results have top predictors, and are excluded from overall AUC calculations.

BRHM succeeds in detecting associations, even when LD is high, while still controlling for false associations. This is more important in the Ames diversity panel, in which the population design does not control for historical population structuring, than in the NAM population design – the null heritabilities, which represent the variance due to global population structure alone, are typically less than the heritabilities for the real phenotypes in the NAM populations, but vary nearly independently in the Ames diversity panel (Figures S7–18). Figure 10 shows extended LD around ZmCCT (the ZmCCT region for all populations is highlighted in supplemental figures S7 and S8), as indicated by a bimodal distribution of estimated heritabilities for a linked region upstream, where heritability is high when no kinships are more proximal to ZmCCT, and near zero when they are. Both the RHM and BRHM methods successfully control for this downstream linked region, as it is not significant in either (although it is possible there is real signal from this region, as a candidate gene is nearby, but it is not detectable due to genome structure near ZmCCT). Although false association is well controlled in both, the RHM method is more conservative than BRHM, with lower resolution; BRHM is significant for a smaller region than RHM around ZmCCT, and identifies additional regions across the chromosome, many nearby flowering candidates. BRHM also controls for false association due to population structure, which is a problem for association mapping in diversity panels when the trait is highly correlated with population structure (Figure 3).

**Figure 10.**
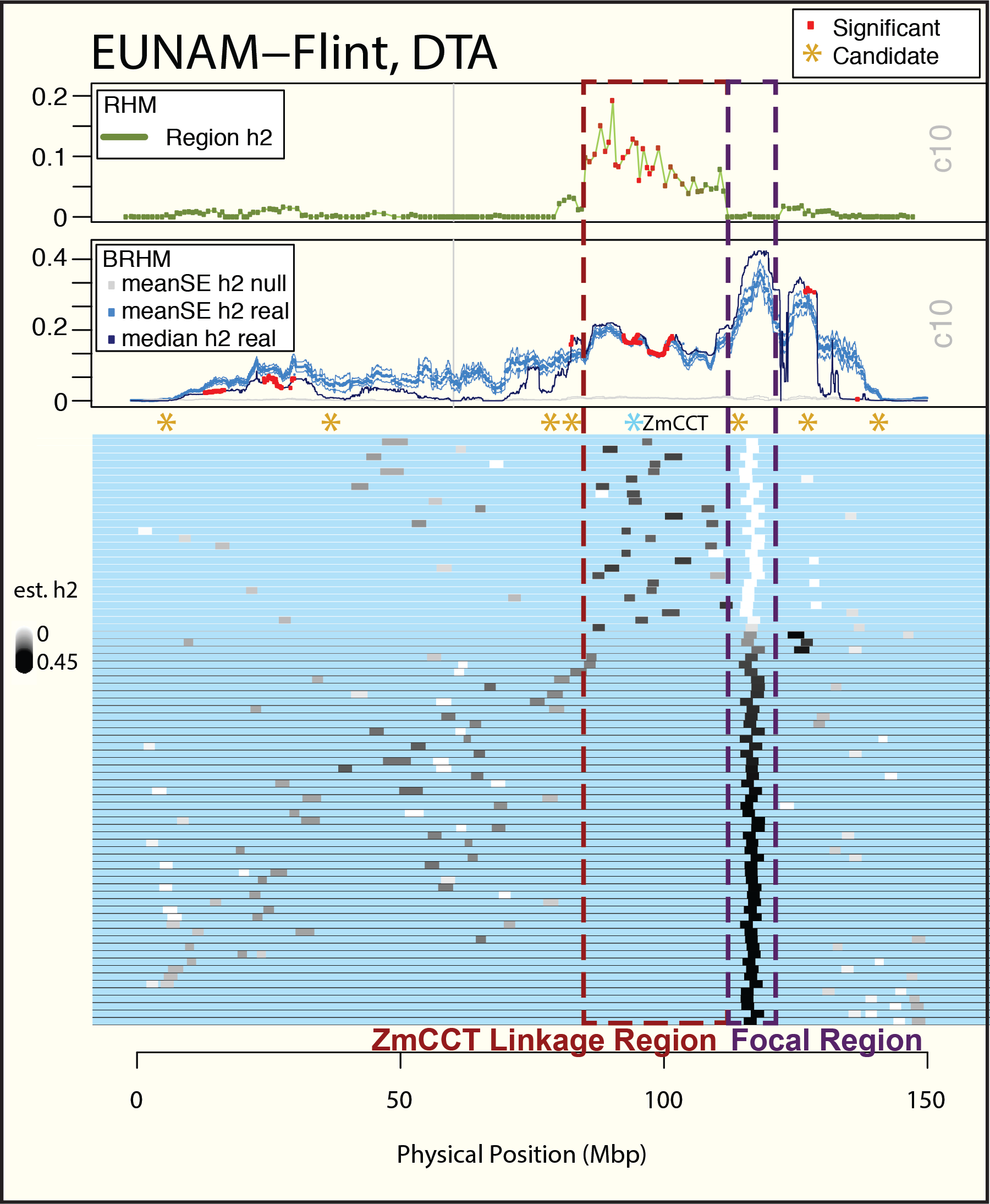
RHM and BRHM results for chromosome 10, and the variances explained for all models in BRHM that contain a kinship that falls on the region downstream of ZmCCT, and has the greatest difference between the median and mean variance estimated. This figure illustrates how large haplotype blocks can generate high variances explained based on variance in sampling, and also how the BHRM model correctly determines significance for such regions, with more power and resolution than RHM to detect candidates.

Overlap in significant regions between populations is low in BRHM. The greatest overlap within significant regions for both populations is between the EUNAM-Flint and USNAM populations, followed by Ames and USNAM (Table 4, Figures S6–7).

**Table 4.** Overlap between significant BRHM results at Bonferroni corrected alpha = 0.01 between populations. Calculated as the shared overlap/total number significant in either population. The top diagonal is for DTS, and the bottom is for DTA.

|  | Ames | CNNAM | EUNAM-Dent | EUNAM-Flint | USNAM |
| --- | --- | --- | --- | --- | --- |
| Ames |  | 0.07 | 0.038 | 0.085 | 0.116 |
| CNNAM | 0.062 |  | 0.064 | 0.079 | 0.096 |
| EUNAM-Dent | 0.048 | 0.061 |  | 0.079 | 0.089 |
| EUNAM-Flint | 0.083 | 0.101 | 0.04 |  | 0.134 |
| USNAM | 0.109 | 0.091 | 0.104 | 0.131 |  |

### Combined GWAS

Directly combining populations for joint GWAS analysis, bypassing the need for meta-analysis, is desirable to avoid the loss of power inherent in meta-analysis. Combining populations was challenging, however, because the phenotypic variances between populations are due to both the different environments in which the samples were grown, but also due to differences in genetic diversity of the datasets (Figure 5). We shrunk population variances using dataset as a random effect, then predicted the value of the reference line, B73, in each population and shifted each value for the population by the difference between B73 in Ames and the predicted B73 for the population. This was done because the heat units in the different studies vary, so this method centers the data on genotypes rather than a grand mean. This shift better standardizes taxa genetically similar to B73 than those more distant, creating another bias reflected in the rank change between EUNAM-Flint and EUNAM-Dent before and after correction, a known error because the populations were grown in similar environments (Figure 5). The biases from combining datasets add to known biases for GWAS conducted for flowering time, and GWAS for populations with extended linkage such as found in the NAMs, resulting in extreme deviation from uniform for p-values (Figure 11). Despite this, the well-known Vgt1 locus and ZmCCT, which is significant in 4 of the 5 populations (Figure 10), shows clear and locally dominant signals (Figure 11, inset). However, genome-wide these signals are not especially notable, inconsistent with previous studies (the highest signal is on chromosome 1, in a region not noted for high effect days to flowering loci, but that is a high-linkage region of high effect in many of the populations (Figure 11)).

**Figure 11.**
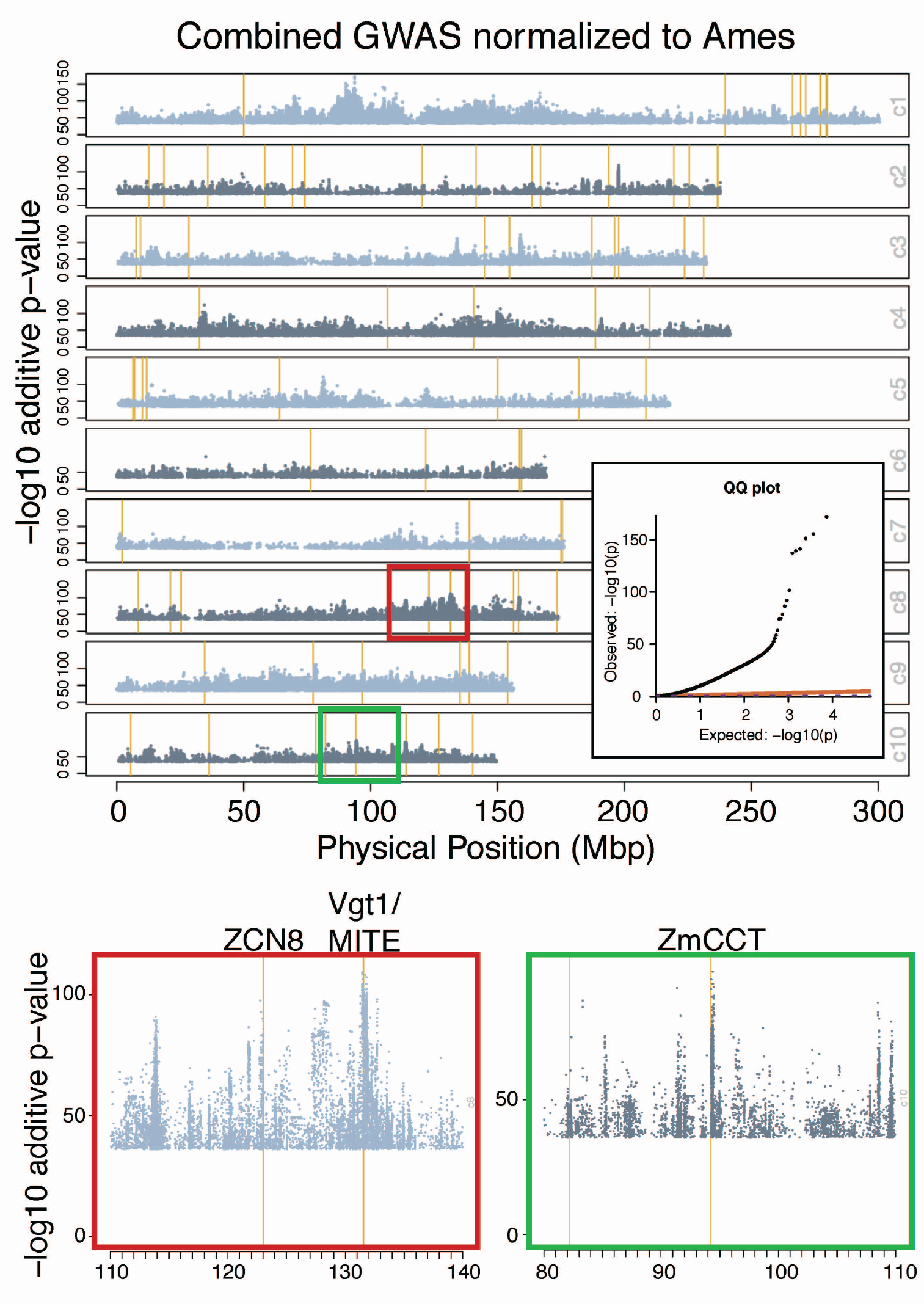
Combined GWAS analysis where traits are shifted by the differences between the predicted value of B73 in the population and the value of B73 in Ames, and the populations shrunken to account for variable GDD in the different study ennvironments. Population structure is controlled by 5PCs and a dataset term. Top 300,000 points by additive p-value plotted. Y-axis tick marks every 50. Vertical yellow lines represent Dong et al/Danilevskya et al candidate genes (Table S2). QQ-plot for combined GWAS results downsampled to 0.0001 (but including the top 10 values). The top flowering time loci in Ames and USNAM are highlighted.

### Candidate gene enrichment

We looked at enrichment for the Dong et al/Danilevskya et al days to flowering candidate genes focusing on the BRHM method. Most of the populations show slight enrichment for flowering candidate genes, between 1-2X (Table 5). Interestingly, CNNAM shows the most enrichment for candidate genes. Candidate overlap between populations is also concentrated among the top few, previously identified, flowering loci, namely ZCN8 and the Vgt1 locus on chromosome 8 and ZmCCT on chromosome 10 (Table S1).

**Table 5.** Proportion of Dong et al candidate genes in significant regions relative to random genes within significant regions. Significance based on Bonferroni corrected significance for BRHM method, Benjamini-Hochberg for RHM, and for SNPs selected in at least a certain number of models for the RMIP method. Random genes in significant regions are the average of 100 samples.

|  | <u>DTA</u> |  |  | <u>DTS</u> |  |  |
| --- | --- | --- | --- | --- | --- | --- |
| **BRHM** | 0.001 | 0.01 | 0.05 | 0.001 | 0.01 | 0.05 |
| Ames | 0.968 (6) | 1.316 (10) | 1.22 (11) | 1.297 (8) | 1.282 (9) | 1.424 (13) |
| CNNAM | 1.579 (6) | 1.777 (11) | 1.31 (11) | 2.174 (8) | 2.609 (15) | 1.724 (16) |
| EUNAM-Dent | 1.019 (7) | 1.046 (8) | 1.012 (12) | 1.372 (9) | 1.171 (10) | 1.033 (14) |
| EUNAM-Flint | 1.156 (8) | 0.937 (8) | 0.915 (13) | 1.441 (10) | 1.202 (12) | 0.904 (13) |
| USNAM | 1.146 (28) | 1.095 (32) | 1.044 (38) | 1.139 (29) | 1.16 (36) | 1.164 (42) |

\*Candidates in significant regions/Average over 100 samples of non-candidate genes from the maize v3 gene models in significant regions (Number of candidates in significant regions)

Comparison of GWAS results for SNPs around physiological candidate genes to random matched subsets shows only enrichment for EUNAM-Dent and, marginally, USNAM without population structure correction in the short candidate list, but in the long list is enriched for Ames, EUNAM-Flint and USNAM (Table 6). With population structure correction, only Ames is significantly enriched for p-values in the short list, but all populations are enriched for genes in the long list. Candidate lists were also generated for the top Ames results and combined GWAS results, based on the sizes of the flowering time candidate lists both with and without population structure correction. Ames is the population with the highest resolution and for which top hits should capture temperate adaptation loci that may or may not be related to flowering explicitly. P-values from around the short 79 Ames gene list is enriched for all populations, but more so for EUNAM-Flint and USNAM without population structure control; with the population term only EUNAM-Flint and CNNAM (DTS) are enriched. Based on the top 918 genes for Ames, only EUNAM-Flint is enriched without population correction, and EUNAM-Flint and CNNAM and USNAM for DTS only. The p-values from SNPs associated with candidates culled from the combined analysis (Ames-centered DTA) were enriched in all populations, which suggests that the combined analysis does tag regions with enriched signal shared across populations.

**Table 6.**
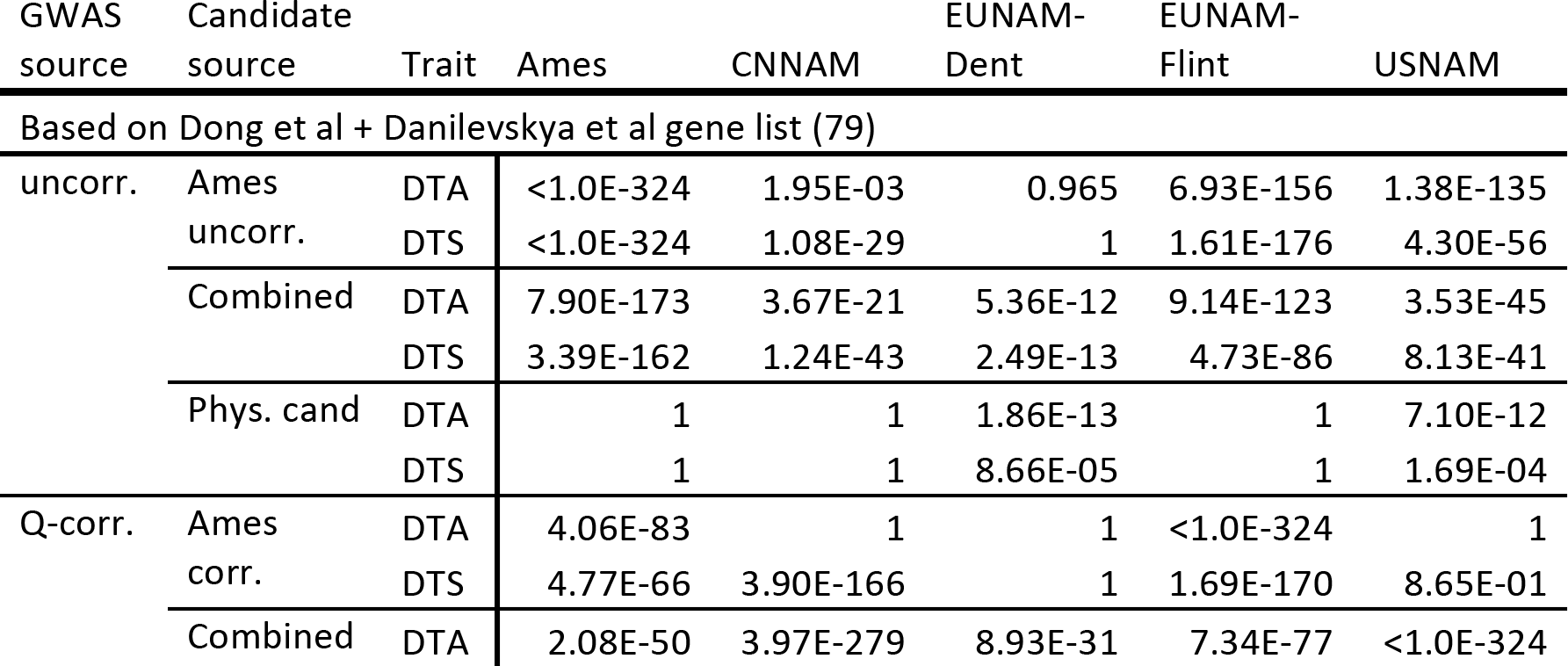
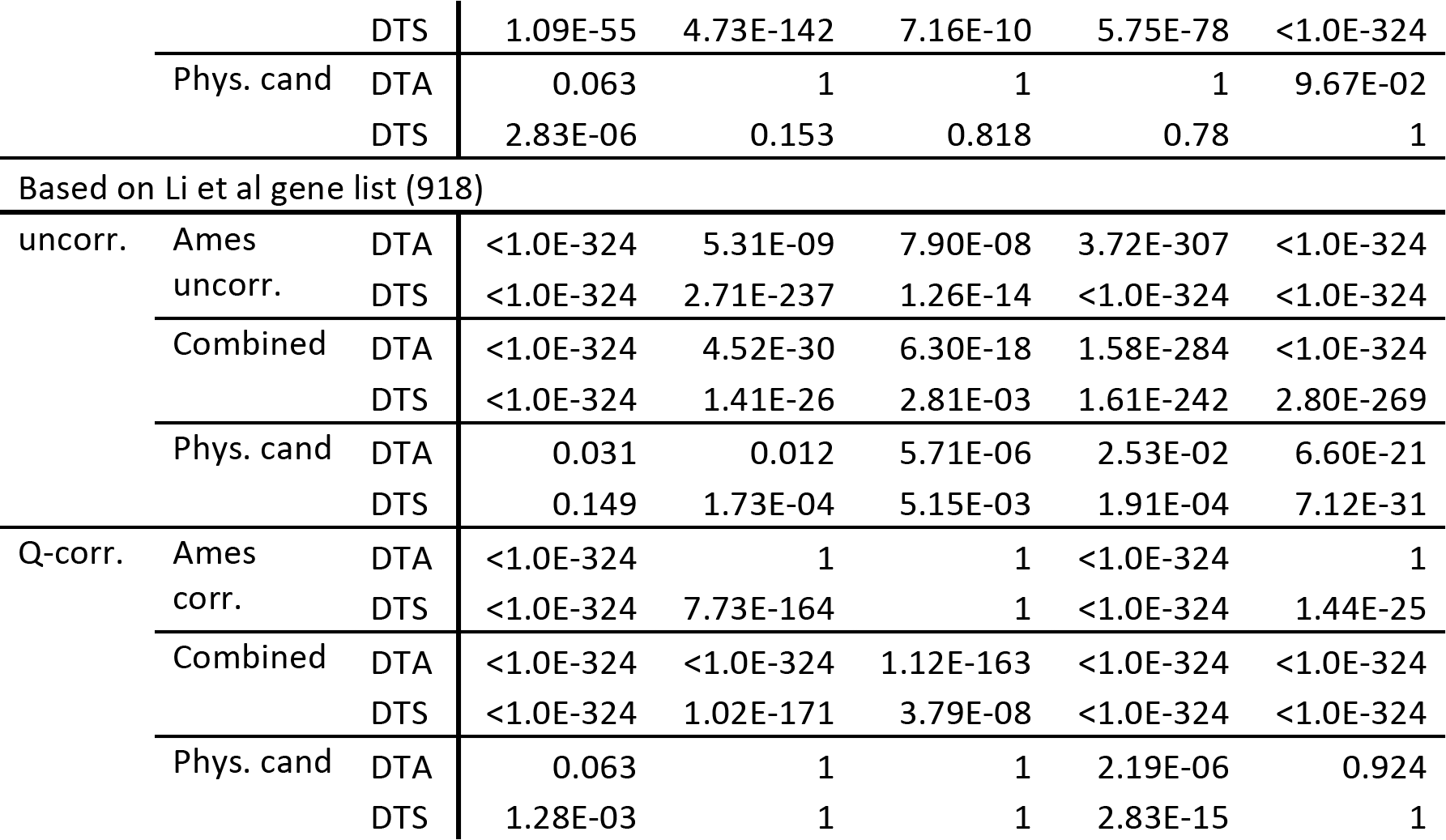
Enrichment for low p-values within 2kbp of candidate gene regions derived from the 79 and 918 flowering candidate gene lists, and matched sized lists from the Ames and combined GWAS candidate lists (Table S2). P-values from within the candidate gene region are compared with a matched set of p-values from outside the regions using a one-sided Wilcoxen signed rank test in R. Only results from the Ames candidate list matching the GWAS source file by trait and population structure correction are presented.

We used machine learning at candidate genes (Li et al, Dong et al plus Danilevskya et al, and Ames without population structure correction) to determine if non-linear combinations of BRHM and GWAS results, power considerations such as MAF and coverage, and measures of population differentiation between populations and on the temperate-tropical gradient could better explain results (Figures 12, S23). We expected this might be the case if differential segregation at candidate genes within different populations generated the observed pattern of minimal overlap within a given population. The ability to predict physiological candidate genes is only slightly enriched above baseline (an AUC of 0.5) when BRHM, GWAS, power and population differentiation are taken into account, suggesting that even non-linear combinations of these predictors cannot explain the candidate lists. Interestingly, in addition to mapping results, coverage statistics also rank high as predictors, suggesting that structural variation may be diagnostic for candidate genes. Prediction of the Ames candidates used Ames BRHM results as the top predictors, unsurprisingly, and were followed by mapping results for other populations, with EUNAM-Dent and CNNAM as the least important predictors among the top BRHM results. That BRHM results, rather than population differentiation scores or even GWAS results, are the best predictors for Ames suggests that the top hits in Ames are generally shared across populations, with EUNAM-Flint and USNAM as the most important and EUNAM-Dent and CNNAM sharing less.

**Figure 12.**
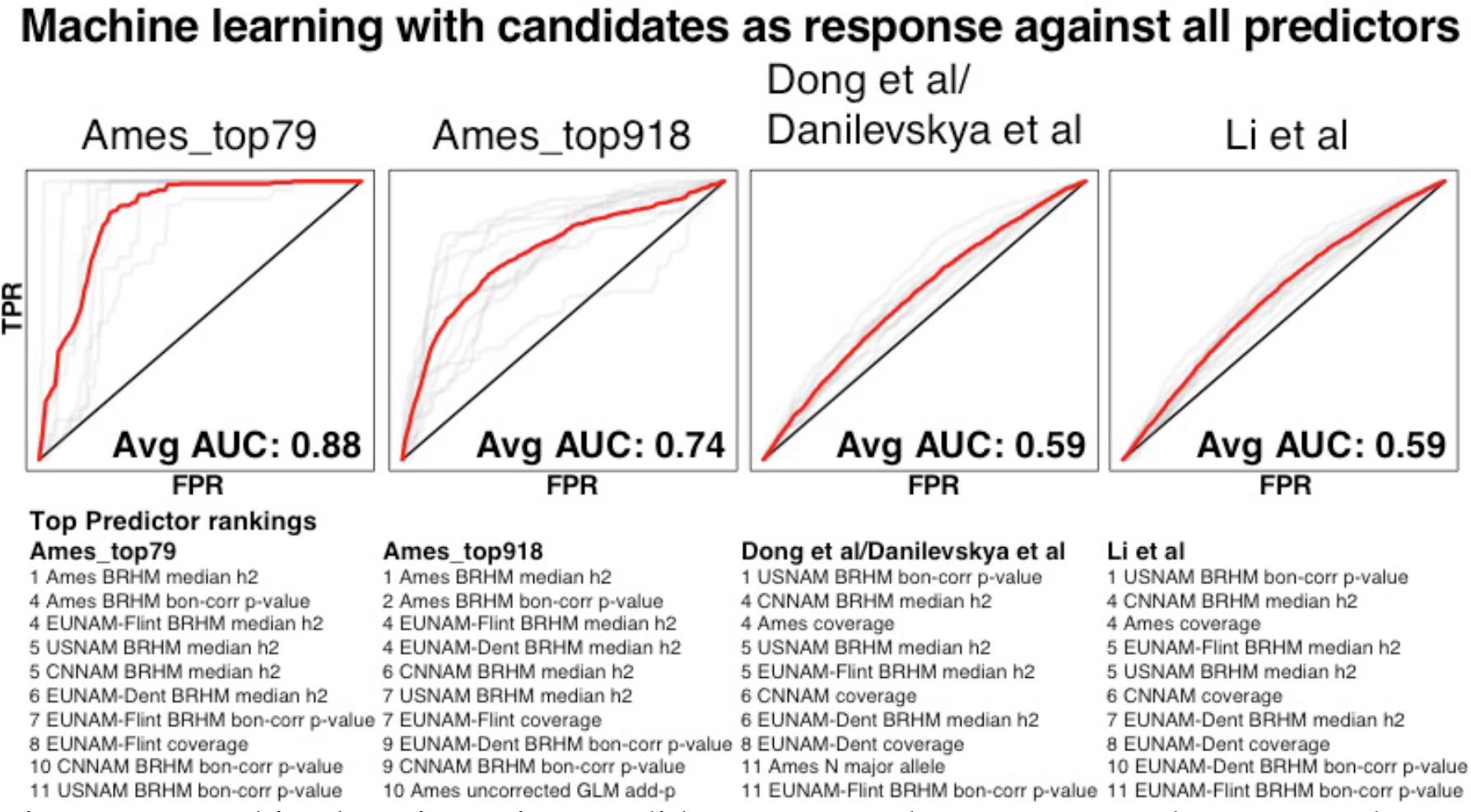
Machine learning using candidate genes as the response, and GWAS and BRHM mapping, power estimators, and F_ST_s as the predictors. The red line represents the mean of the chromosomal ROC curves, sampled at 0.01 intervals. Candidate genes were classified as 1, with a 20kbp buffer around the gene start and end position (maize AGP v3). AUCs of 0 result when no candidates are found on the chromosome tested. Not all chromosomes for the Ames top 79 candidates have top predictors, and are excluded from overall AUC calculations.

### Go term enrichment from top genes from BRHM analysis

The limited overlap in significant regions and candidate lists across all of the methods is not consistent with previous findings (*8*, *27*) using lower density SNP sets, and not supported by the generally high cross-population predictive abilities. The discrepancy could be due to truly different underlying genetic architecture that had previously been collapsed due to limited marker coverage, which should generate low cross-population predictive abilities, differential allelic series at common genes across populations as was found in the USNAM (*5*), and EUNAM (*16*) populations, or it could be due to differences in population structure, both historical and recent, shifting the signal from or differentially segregating for the same causal variant within populations. We looked at GO term enrichment for top BRHM results to see if association regions share a functional enrichment to help reconcile the disparities between the limited overlap observed in the mapping results, and the broadly shared genetic basis for flowering in most populations implicated in the cross population prediction results.

We focus on the top 1,000 genes from the BRHM analysis for each of the populations to see if differences in mapping results are also related to differences in function (Table 7). A chi-square goodness of fit test for equal representation within biological functional categories reports a highly significant deviation from equal representation across populations, with a p-value of 7.54e-17 for DTA. All of the populations show similar levels of enrichment for gene models related to housekeeping, biotic response and macromolecules, a catchall category for macromolecule biosynthesis, metabolism, and catabolism (Table S5). Reproduction, signaling, development and biotic response are the classes with the largest deviations from expected. Ames and USNAM, which have the greatest genetic overlap, have distinctly different enrichment profiles, likely reflecting differences in resolution between the populations, and absolute variance in days to flowering.

**Table 7.** GO term enrichment against expectation of even representation for each population, based on the top 1,000 genes from the most significant regions of the genome for DTA, as determined by BRHM analysis. Chi-square goodness of fit p-value for test of deviation from the null hypothesis that there are no differences between populations for annotated function is 7.54E-17

|  | Ames | CNNAM | EUNAM-Dent | EUNAM-Flint | USNAM |
| --- | --- | --- | --- | --- | --- |
| AbioticResponse | 1.026 | 0.982 | 0.765 | 1.002 | 1.224 |
| BioticResponse | 0.993 | 1.034 | 0.938 | 1.072 | 0.969 |
| Development | 1.016 | 0.894 | 0.866 | 0.935 | 1.28 |
| Housekeeping | 1.014 | 0.994 | 1.02 | 1.005 | 0.97 |
| MacroMolecules | 0.974 | 1.017 | 1.042 | 0.994 | 0.97 |
| Regulation | 0.958 | 1.05 | 1.107 | 0.978 | 0.903 |
| Reproduction | 1.159 | 0.937 | 0.83 | 1.156 | 0.952 |
| Signalling | 1.267 | 0.885 | 0.869 | 0.946 | 1.059 |

## Discussion

Complex traits, such as yield, are well known to aggregate fitness effects across all pathways and systems of a plant, but this study highlights that this is also true for days to flowering (see also Li et al (*32*)). When a plant is induced to flower depends on the alleles present at ZmCCT, ZCN8, and Vgt1, but it also depends on the pleiotropic response of hundreds of other loci responding to signals for heat, circadian signaling, light quality, moisture availability, starch accumulation, and others (*3*, *4*). Functional biological constraints ensure some overlap across populations for high effect genes in the autonomous flowering pathway; this is supported by the increase in genome-wide predictive ability for poorly predicted populations when the top significant SNPs from any population are used. Candidate gene enrichment suggested around ten genes may be universally important and segregating. However, limited overlap in mapping results and differential GO term enrichment suggest that, as maize spread across the world during improvement and modern hybrid breeding, complex population dynamics may have led to differential selection for secondary pathways implicated in temperate adaptation. These patterns are found *Arabidopsis thaliana* as well. Fournier et al find evidence for selection on different, pleiotropic, QTL in different environments across Europe in a common garden experiment (*81*). Additionally, Kover et al (*82*) find that adaptation to early flowering is partially conditional on other conditions in the selection environment. Finally, only two out of a total of 10 QTL were identified between Swedish and Italian RIL populations; however, unlike our findings in maize, candidate genes are more highly enriched in QTL results (*83*).

That some populations, namely EUNAM-Flint and CNNAM, do not predict Ames as expected based on population differentiation suggests more complicated population dynamics than simple relatedness (Figure 7). What better explains the patterning in predictive ability is the spread of the population founders on North American temperate/tropical gradient (Figure 4). The two populations with a localized geographic origin on this gradient, CNNAM and EUNAM-Dent, are the two populations with more limited cross-population predictive ability, and low predictive ability for GWAS results in machine learning. In contrast, EUNAM-Flint, which has higher than expected cross-population predictive ability when predicting USNAM, has parents spanning the tropical/temperate North American gradient, and F_ST_s between Northern Flints and tropical American germplasm from Hapmap 3.21 are top predictors in machine learning against uncorrected GWAS results. Lower than expected predictive ability relative to population differentiation could be interpreted in two ways; a narrow germplasm base or, if the population has high variance for flowering time, that these populations contain novel temperate adaptation not captured on either American temperate / tropical axis.

The history of germplasm introduction can shed light on the discrepancies between predictive ability and population differentiation especially for the EUNAM-Dent and CNNAM. The earliest germplasm in Europe was Caribbean in origin, and subjected to selection for early flowering upon entry into Spain, but it was only after the introduction of the Northern Flint landraces in the mid-1600s from the northeast of the modern US that maize agriculture spread to European climates north of the Pyranees (*44*, *47*, *48*, *84*). Genetic evidence from European maize suggests that most of the early flowering adaptation was acquired from the North American Northern Flint germplasm introduced in the 1600s (*44*, *47*), but also that there are unique rare alleles in the Southern European (Spanish and Portuguese) germplasm (*48*). Finally, in the past century, the development of the European heterotic groups introduced primarily American Iodent, Stiff Stalk, Lancaster, and Minnesota germplasm into Europe, captured primarily in the EUNAM-Dent panel (*47*, *85*, *86*). Although Chinese maize now contains both early and late flowering varieties, the most likely first point of entry for maize to China was by Spanish and Portuguese traders through the Port of Macau, and these are reported to have been humid, tropical adapted varieties (*45*, *46*). A regular trading route between Acapulco and Manila, which started in CE 1565, could have introduced western lowland Mexican maize to Asia (*87*). This panel of Chinese lines is dominated by lines whose origins likely predate modern hybrids of the 20^th^ century, suggesting an indigenous temperate adaptation to China based.

The recurrent parent of the EUNAM-Dent population is representative of the agronomically important Iodent germplasm (derived from an Iodent/BSSS cross), and the additional lines in the EUNAM-Dent panel derive from the US Stiff Stalk, US Lancaster, and Hohenheim Dent populations (*88*, *89*). The Hohenheim Dents were bred from US temperate germplasm, and especially the early flowering Minnesota lines (*86*), which result from crosses between the US Southern Dents and Northern Flints, followed by selection for extreme temperate adaptation, with early flowering contributed by the Northern Flints (*85*). EUNAM-Dent is well predicted by Ames, but not USNAM; Ames is enriched for temperate US germplasm, including Iodent and Minnesota lines (*11*), while USNAM is not (*60*). It is not surprising that Ames then predicts the EUNAM-Dent panel, as it is a superset, but it is perhaps surprising that the Ames GWAS results predict the EUNAM-Dent germplasm so poorly in machine learning, given that the pedigree suggests that temperate adaptation in the EUNAM-Dent panel is Northern Flint in origin. The lack of overlap in significant regions and poor cross-population prediction suggests that the EUNAM-Dent germplasm base is narrow with respect to flowering such that there is less variance for flowering time or temperate adaption. This is possible, given the good prediction of Dent germplasm by Ames, and moderately supported by the relatively low, relative to Ames and USNAM, narrow-sense heritability for the EUNAM-Dent panel, 0.7 (DTS) and 0.61 (DTA). Additionally, Giraud et al found (*16*) fewer QTL in the EUNAM-Dent relative to the EUNAM-Flint panel, and these QTL explain less variance. Alternately, poor overlap and machine learning prediction could be a result of recent selection for temperate adaptation in the development of the Minnesota germplasm and subsequent selection in Europe, which is not sufficiently represented in the other datasets.

Almost all of the CNNAM parents derive from Chinese sources across heterotic groups, and the recurrent parent is a derivative of the temperate Chinese landrace TangPiSingTou (*90*). While most of the CNNAM parents are temperate adapted (*32*), they are significant at the major photoperiod locus ZmCCT confirming that the population contains tropical alleles. Additionally, broad sense heritabilities for CNNAM were high (0.91 and 0.90 for DTA and DTS respectively) so the genetic basis for days to flowering is not narrow. High heritability and lack of overlap in significant regions and poor cross-population prediction across all populations suggests that the CNNAM population does not suffer from a narrow germplasm base but rather contains novel alleles for temperate adaptation, suggesting an independent origin relative to the other four populations.

The minimal enrichment for physiological candidate genes, even in a machine learning framework that can incorporate information from all of the predictors in a non-linear framework, suggest that days to flowering is primarily mapping temperate adaptation in maize. If temperate adaptation is a suite of traits that incorporates more loci than those associated with the autonomous flowering time pathway, and independent adaptations for these traits from China and the US/Europe are relevant in NAM germplasm, genes in significant regions should be enriched for different non-autonomous flowering loci. The CNNAM and EUNAM-Dent panels show different patterns of enrichment in the top 1,000 genes from the other panels, tentatively supporting this hypothesis. In contrast, Ames and the EUNAM-Flint population are the most enriched of all the populations for reproductive GO terms, perhaps helping explain how these populations have good cross-population prediction, despite high population differentiation.

However, questions remain as to why Ames, USNAM, and EUNAM-Flint, which all share parents across the temperate tropical gradient, have high cross-population predictive ability, and can predict each others' GWAS results reasonably well (EUNAM-Flint) or very well (Ames and USNAM) do not share more significant loci in the mapping results. The good cross-population predictions and machine learning results suggest that these three populations share or partially share a common genetic architecture, and that the lack of overlap is an artifact of our inability to isolate signals within the data. We had hoped to achieve more resolution than is typically possible in NAM populations in meta-analysis using whole-genome markers, but differences in population structure, and how LD shifts the signal in mapping results from true causal loci, certainly contribute to lack of overlap in these structurally diverse populations. Environment, and genotype-by-environment interactions in the testing environments may also explain some of this discrepancy. The construction of the NAM designs reduce the spread of days to flowering across the population to those that can be crossed, reducing the total variance in ways which may be subjected to similarly unique pressures, even across multi-year/location studies. Likewise, the EUNAM trials were all located in Europe, much colder and further north than most of the USNAM or Ames environments; the EUNAM populations predict B73 to flower a full two weeks later than in Ames, due to reduced GDD accumulation, and systemic genotype by environment interactions may affect associated regions. Finally, it is possible that some of the differences between EUNAM-Flint and the USNAM result from partially different genetic architecture derived from early Spanish selection for temperate adaption in early-introduced Caribbean germplasm.

These results raise the question of the suitability of simple threshold values for detecting similarity in complex, population structure confounded traits like flowering time (*91*). Significance or heritability thresholds are modulated by population diversity, population allele frequency, LD with neighboring SNPs, population structure of the founders, and the unique set of environments the populations were evaluated in. This almost guarantees that there will not be a simple threshold to look for meaningful overlap. Machine learning provides an opportunity to look for site overlap in much more rigorous way, by allowing threshold values to vary we can better model non-linear relationships between power, true causal variants and population structure to understand genetic architecture and target causal variation more precisely for breeding. Finally, this study highlights in CNNAM a unique set of germplasm not extensively utilized outside of China, which may provide a source of novel alleles for breeding programs.

## Acknowledgements

The authors thank Robert Bukowski from BRC Bioinformatics Facility, Institute of Biotechnology at Cornell University, for early access to the KNNi imputed HapMap 3 dataset, and Jeffrey Ross-Ibarra and Matt Hufford for early access to the American landraces. This work was supported by National Science Foundation Grants IOS- 0922493and IOS-1238014 and the US Department of Agriculture–Agricultural Research Service.

## Supplemental figures

**Figure S1.**
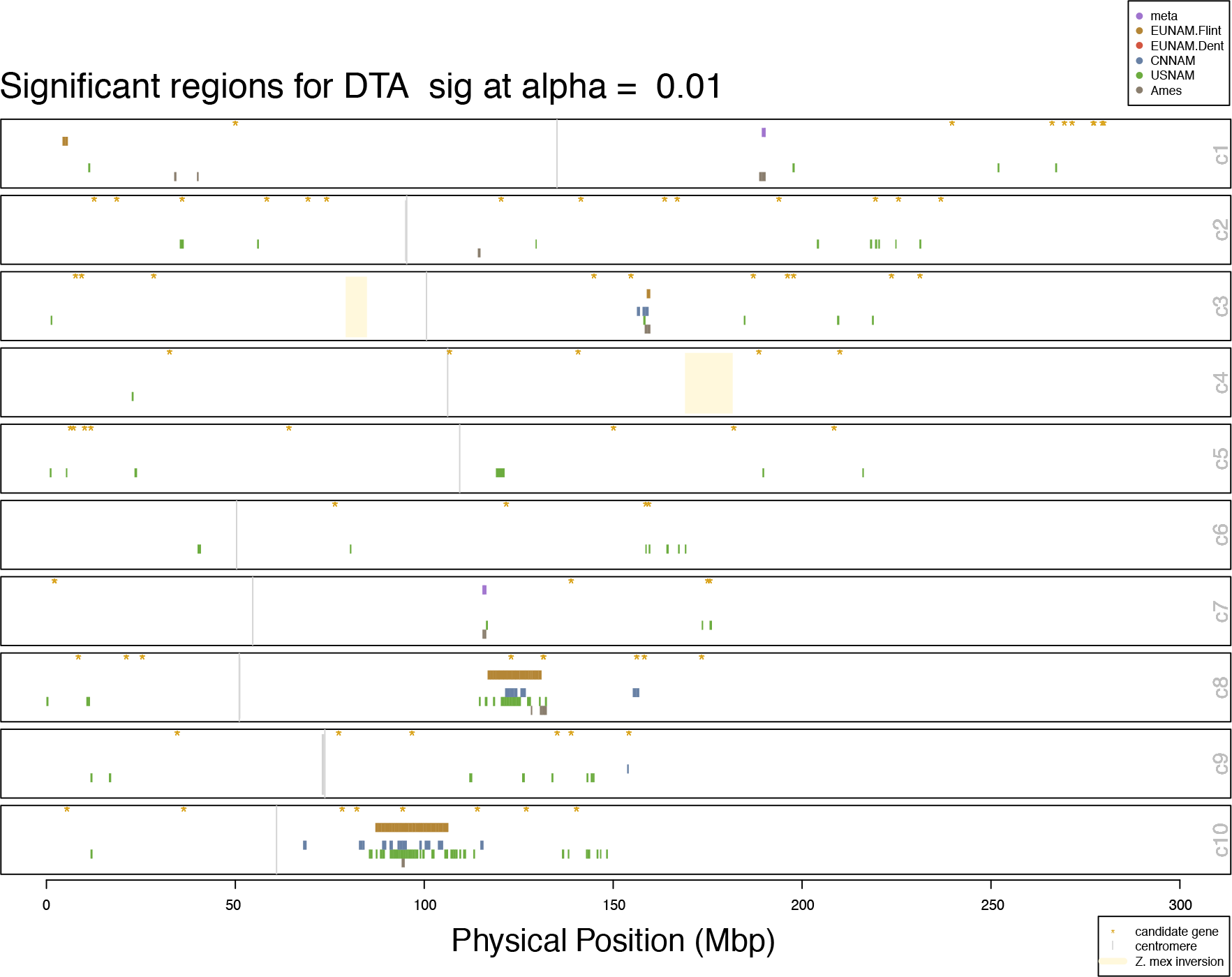
Significant regions and meta-analysis results for RHM method, DTA

**Figure S2.**
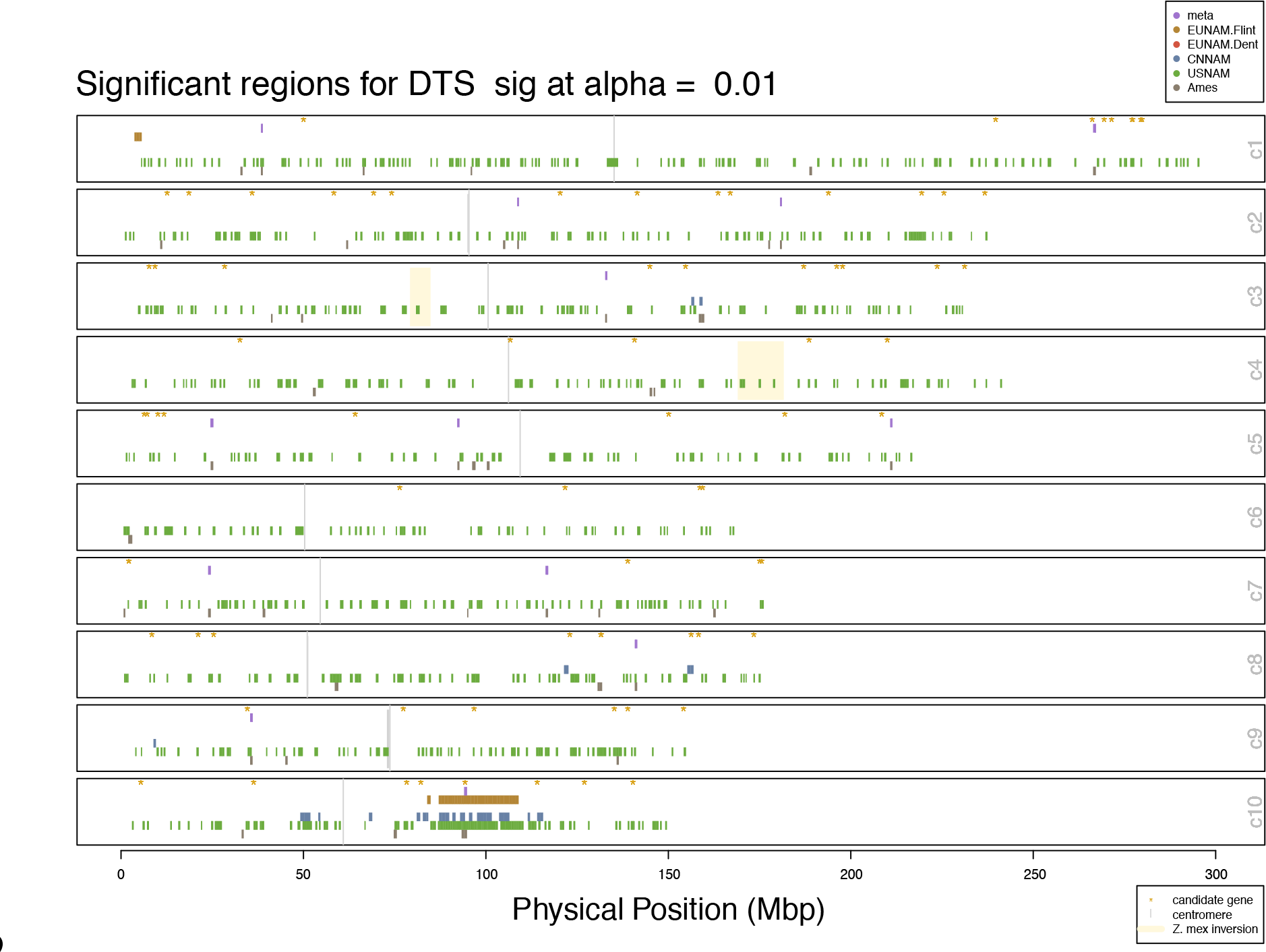
Significant regions and meta-analysis results for RHM method, DTS

**Figure S3.**
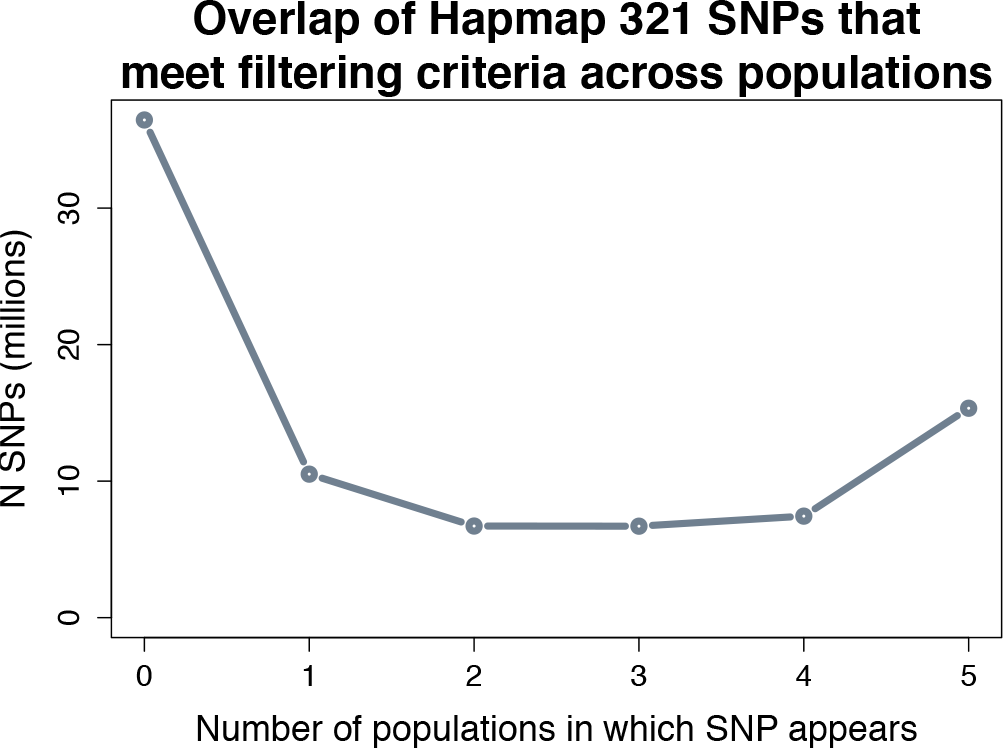
SNP overlap between populations in filtered projected Hapmap3.21 genotypes

**Figure S4.**
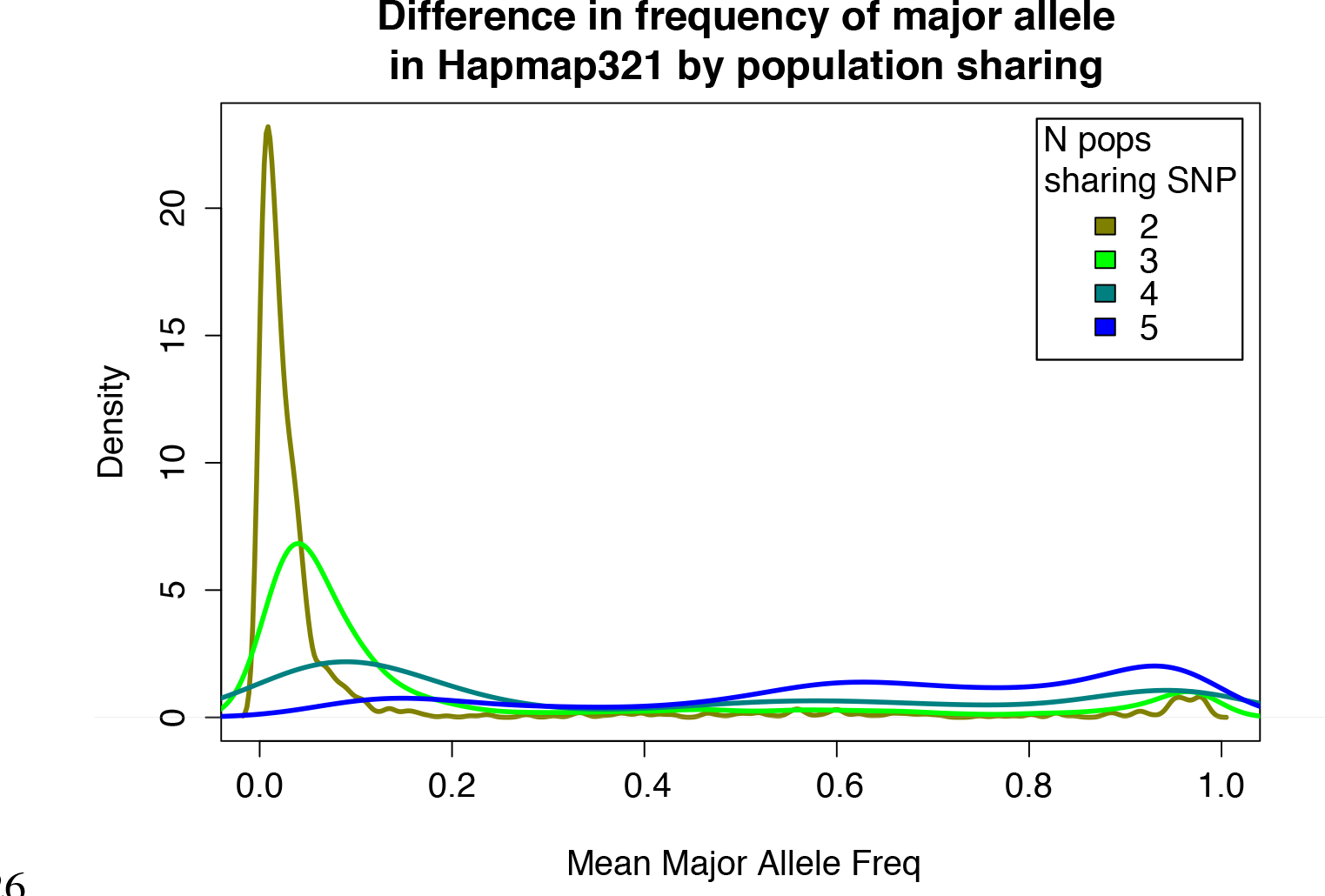
Density of the difference between the minimum and maximum Hapmap 3.21 major allele frequencies in shared filtered projected Hapmap3.21 genotypes

**Figure S5.**
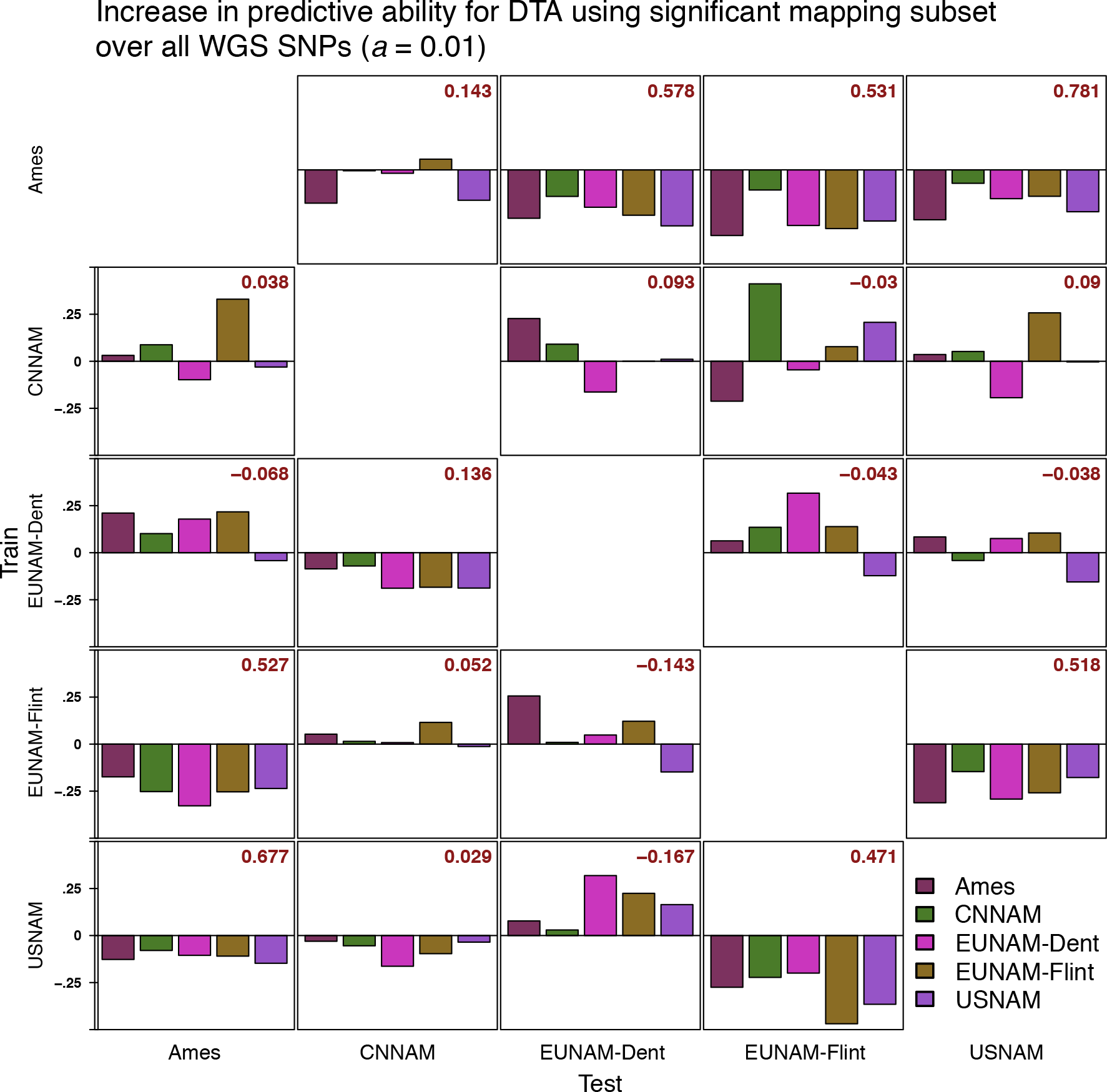
Increase in prediction accuracypredictive ability for DTA using subsets of SNPs from significant regions in BRHM results. Whole genome accuracies reported in red.

**Figure S6.**
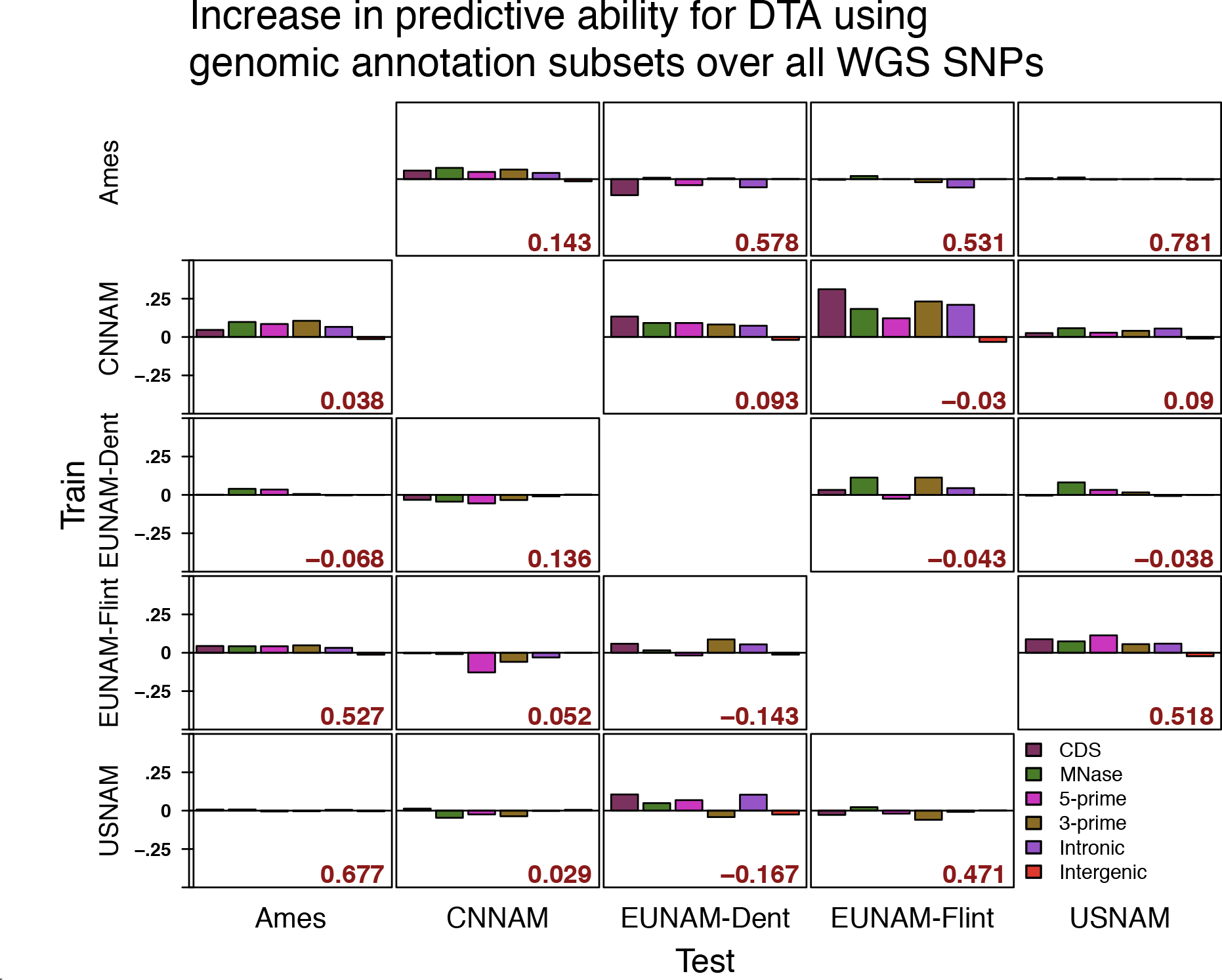
Increase in predictive ability for DTA using subsets of SNPs based on genomic annotations.

**Figure S7.**
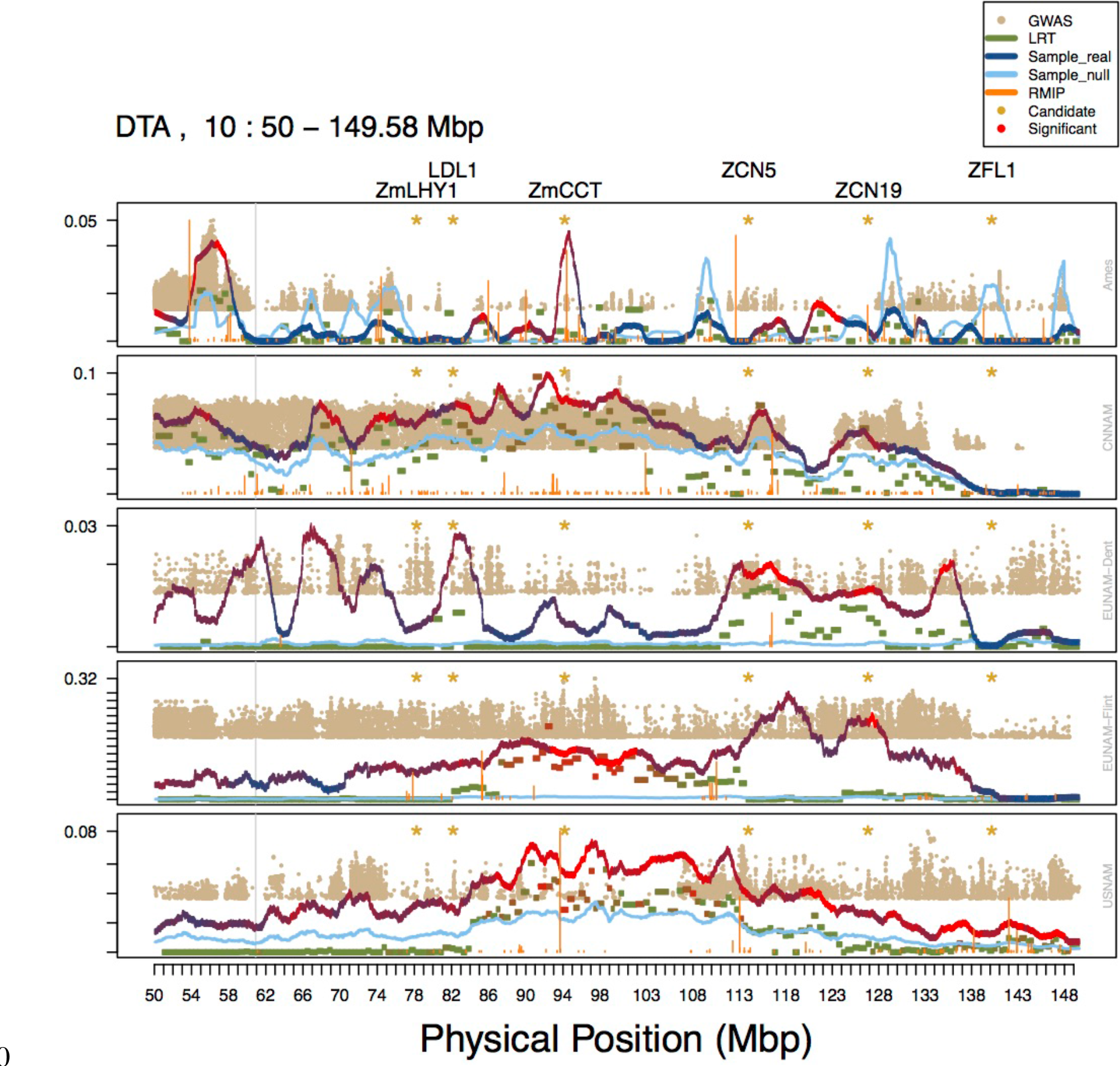
Combined results for all methods (GWAS (no population structure control), resampling GWAS (RMIP), RHM, and BRHM) for 50 Mbp region on Chromosome 10. RHM and BRHM variance estimates are unmodified. GWAS –log10 p-values and the proportion of models for which a SNP was chosen in resampling GWAS rescaled to the maximum value of the variances in the window, and only the top 5,000 SNPs across the genome plotted in a window. Both the median estimated heritability for the null phenotypes (light blue) and the real estimated heritability (blue to red) is shown for BRHM. RMIP bars in orange represent the proportion of models in which that SNP was chosen. Candidate genes (based on Dong et al./ Danilevskya et al) are noted with a yellow asterisk and listed across the top.

**Figure S8.**
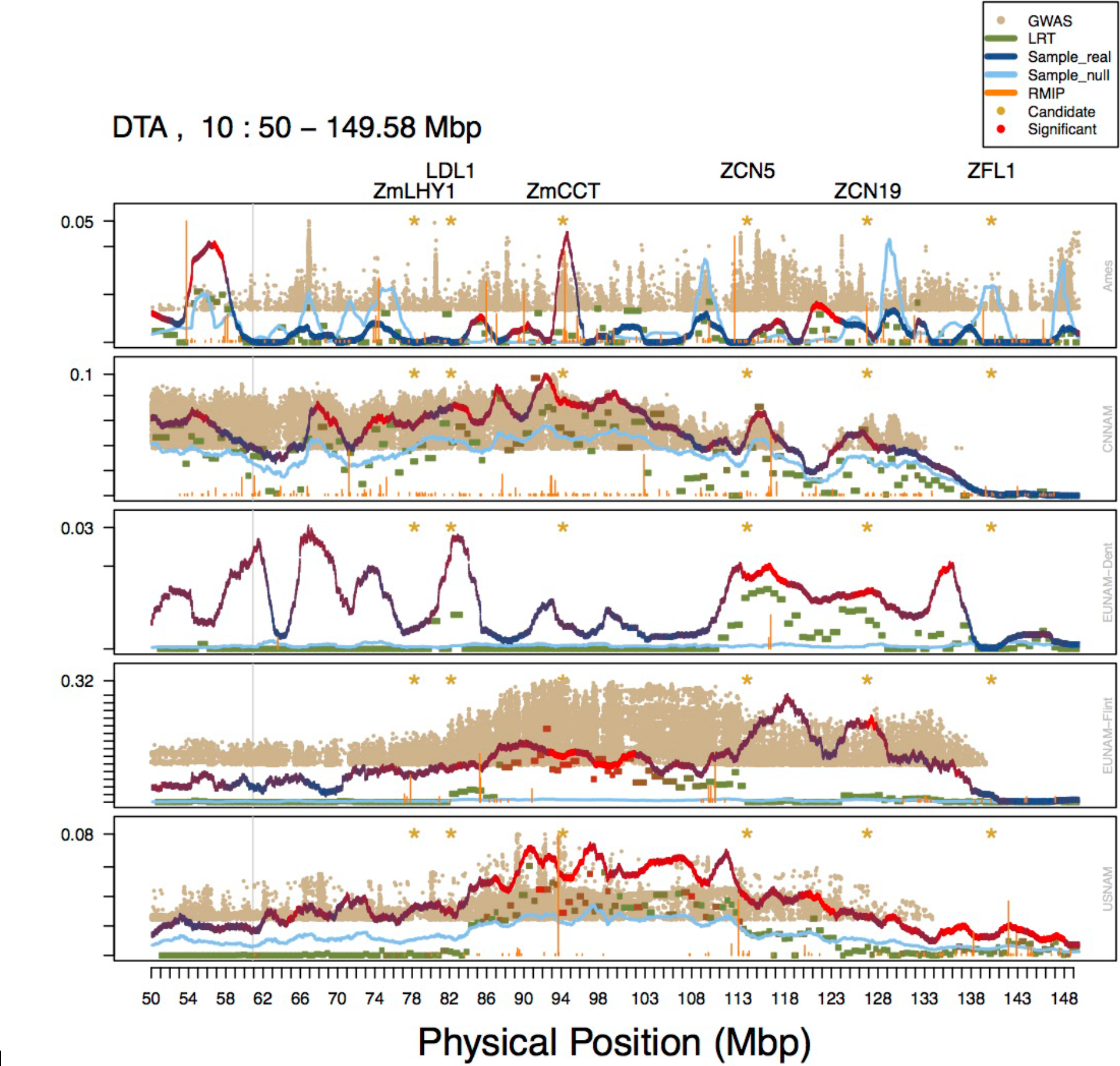
Combined results for all methods (GWAS (Family term or 5PCs for Ames), resampling GWAS (RMIP), RHM, and BRHM) for 50 Mbp region on Chromosome 10. RHM and BRHM variance estimates are unmodified. GWAS –log10 p-values and the proportion of models for which a SNP was chosen in resampling GWAS rescaled to the maximum value of the variances in the window, and only the top 5,000 SNPs across the genome plotted in a window. Both the median estimated heritability for the null phenotypes (light blue) and the real estimated heritability (blue to red) is shown for BRHM. RMIP bars in orange represent the proportion of models in which that SNP was chosen. Candidate genes (based on Dong et al./ Danilevskya et al) are noted with a yellow asterisk and listed across the top.

**Figure S9.**
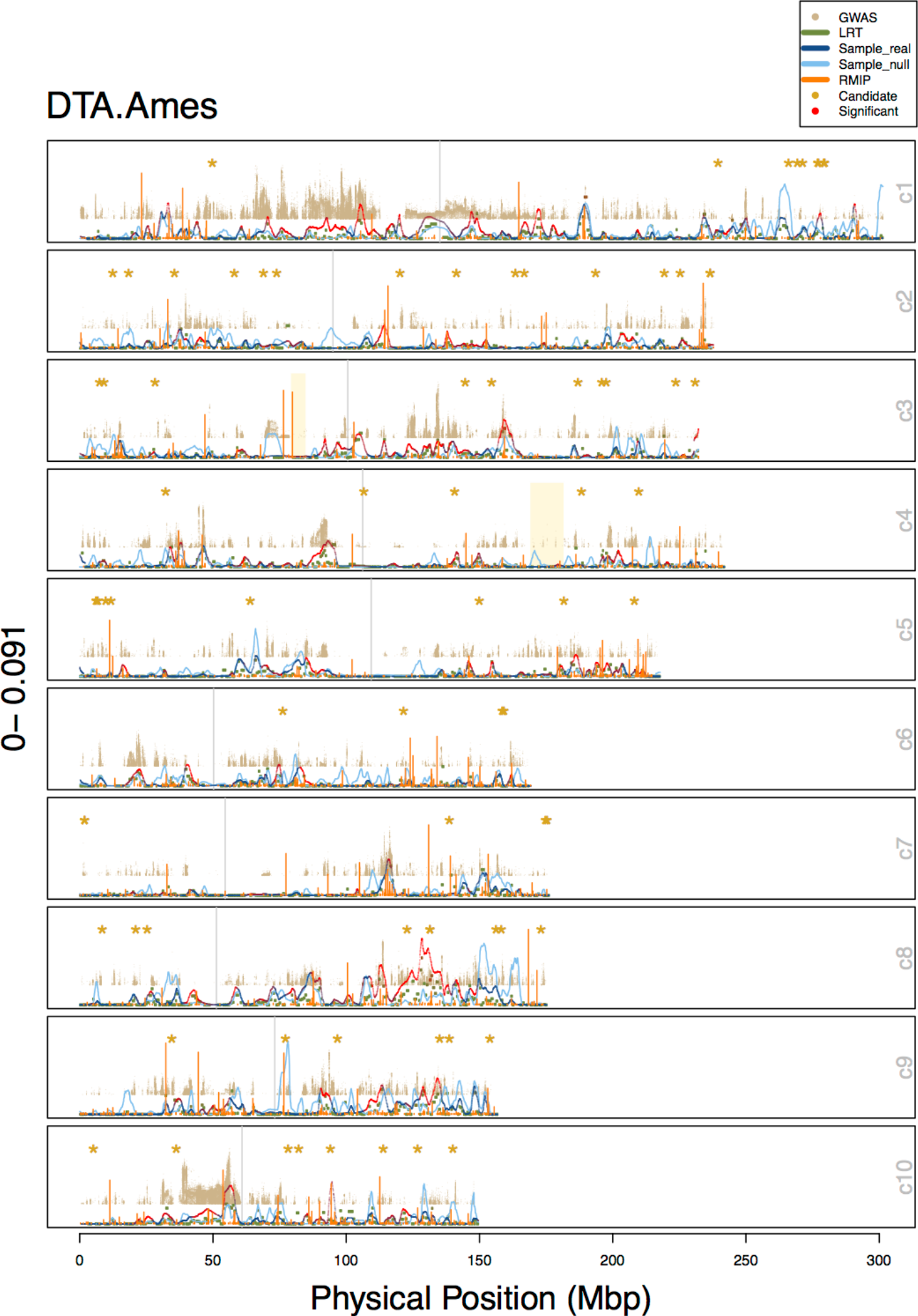
Combined results for all methods (GWAS (uncorrected), resampling GWAS (RMIP), RHM, and BRHM) for Ames, DTA. RHM and BRHM variance estimates are unmodified. GWAS –log10 p-values and the proportion of models for which a SNP was chosen in resampling GWAS rescaled to the maximum value of the variances in the window, and only the top 5,000 SNPs across the genome plotted in a window. Both the median estimated heritability for the null phenotypes (light blue) and the real estimated heritability (blue to red) is shown for BRHM. RMIP bars in orange represent the proportion of models in which that SNP was chosen. Candidate genes (based on Dong et al./ Danilevskya et al) are noted with a yellow asterisk and listed across the top.

**Figure S10.**
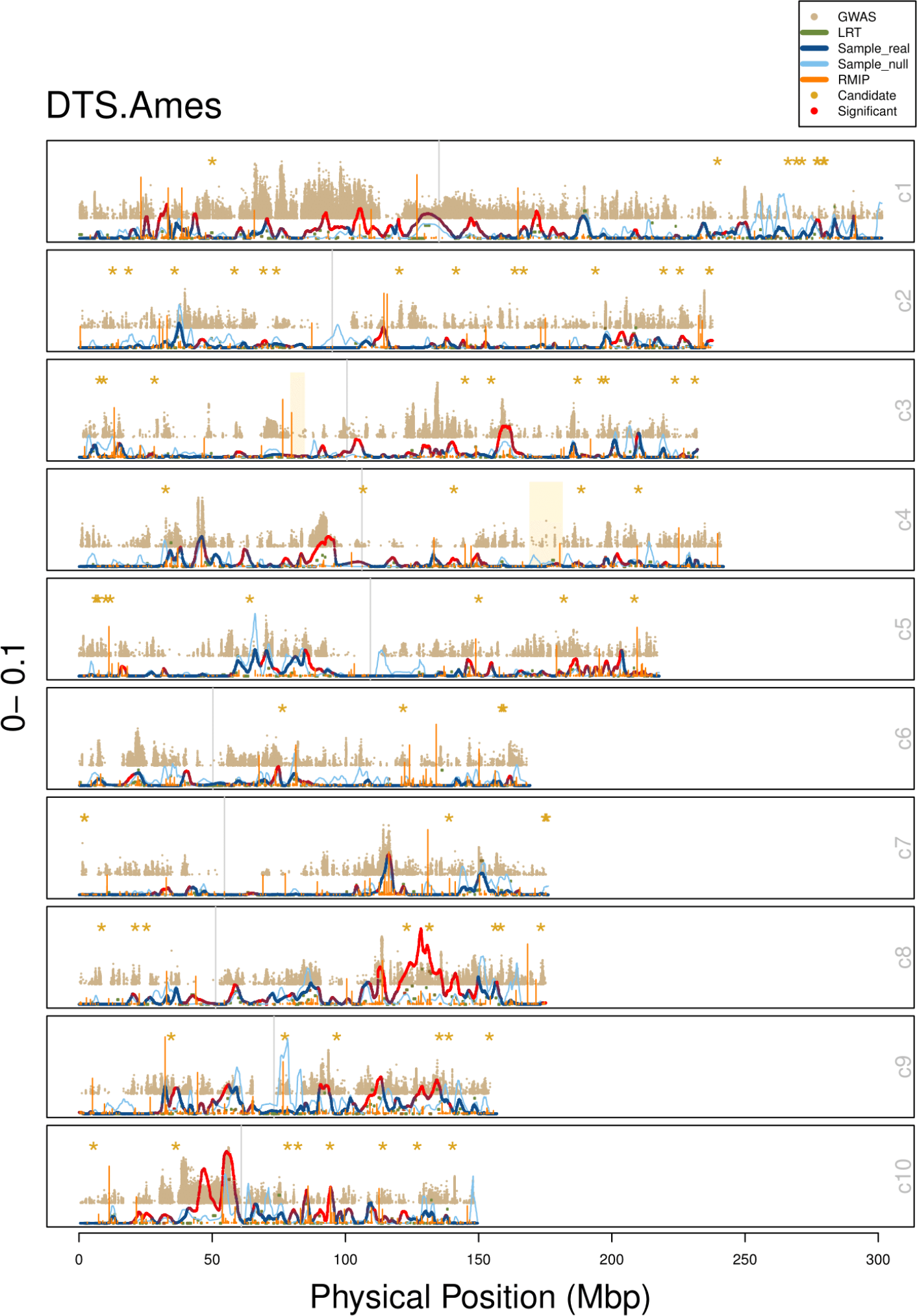
Combined results for all methods (GWAS (uncorrected), resampling GWAS (RMIP), RHM, and BRHM) for Ames, DTS. RHM and BRHM variance estimates are unmodified. GWAS –log10 p-values and the proportion of models for which a SNP was chosen in resampling GWAS rescaled to the maximum value of the variances in the window, and only the top 5,000 SNPs across the genome plotted in a window. Both the median estimated heritability for the null phenotypes (light blue) and the real estimated heritability (blue to red) is shown for BRHM. RMIP bars in orange represent the proportion of models in which that SNP was chosen. Candidate genes (based on Dong et al./ Danilevskya et al) are noted with a yellow asterisk and listed across the top.

**Figure S11.**
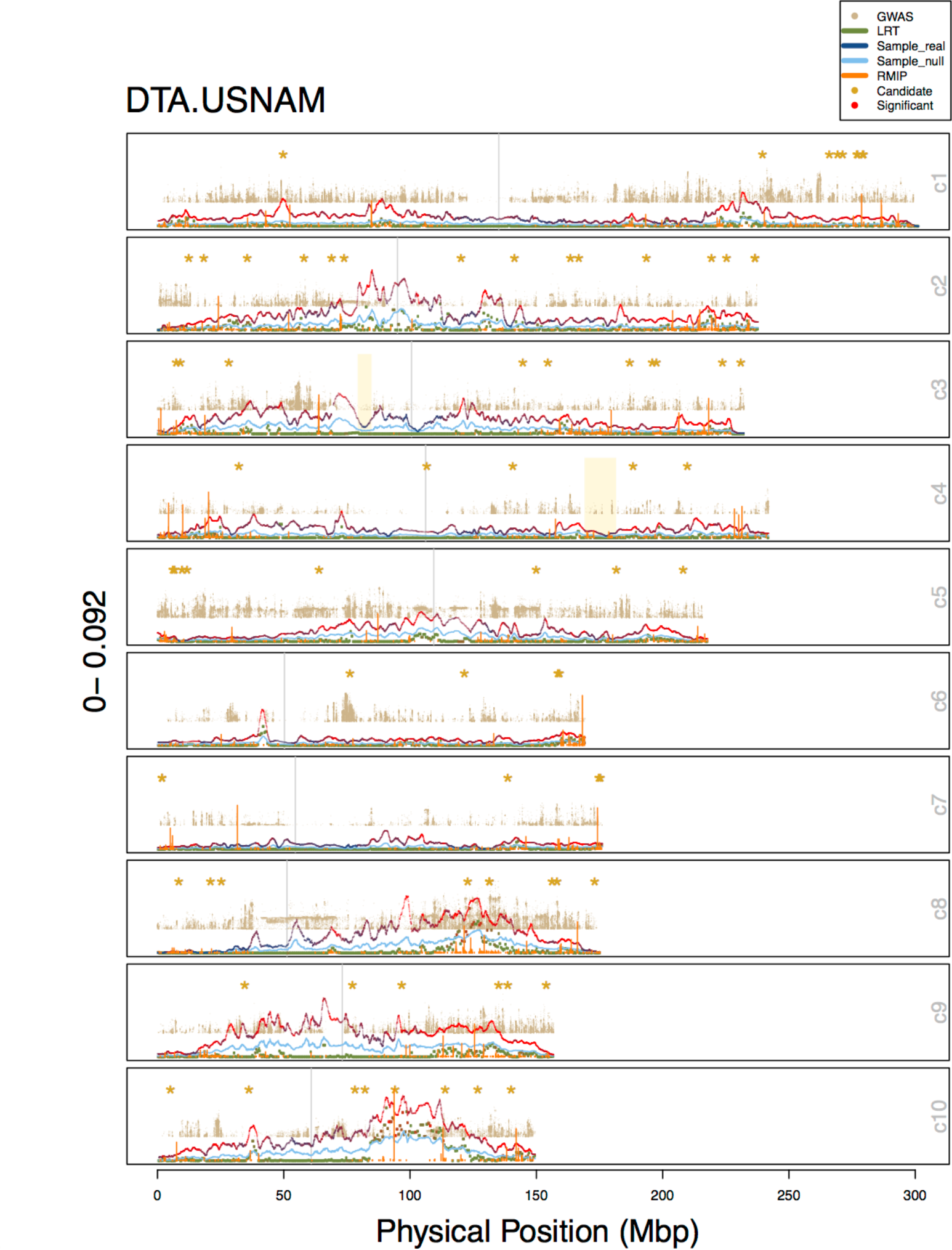
Combined results for all methods (GWAS (uncorrected), resampling GWAS (RMIP), RHM, and BRHM) for USNAM, DTA. RHM and BRHM variance estimates are unmodified. GWAS–log10 p-values and the proportion of models for which a SNP was chosen in resampling GWAS rescaled to the maximum value of the variances in the window, and only the top 5,000 SNPs across the genome plotted in a window. Both the median estimated heritability for the null phenotypes (light blue) and the real estimated heritability (blue to red) is shown for BRHM. RMIP bars in orange represent the proportion of models in which that SNP was chosen. Candidate genes (based on Dong et al./ Danilevskya et al) are noted with a yellow asterisk and listed across the top.

**Figure S12.**
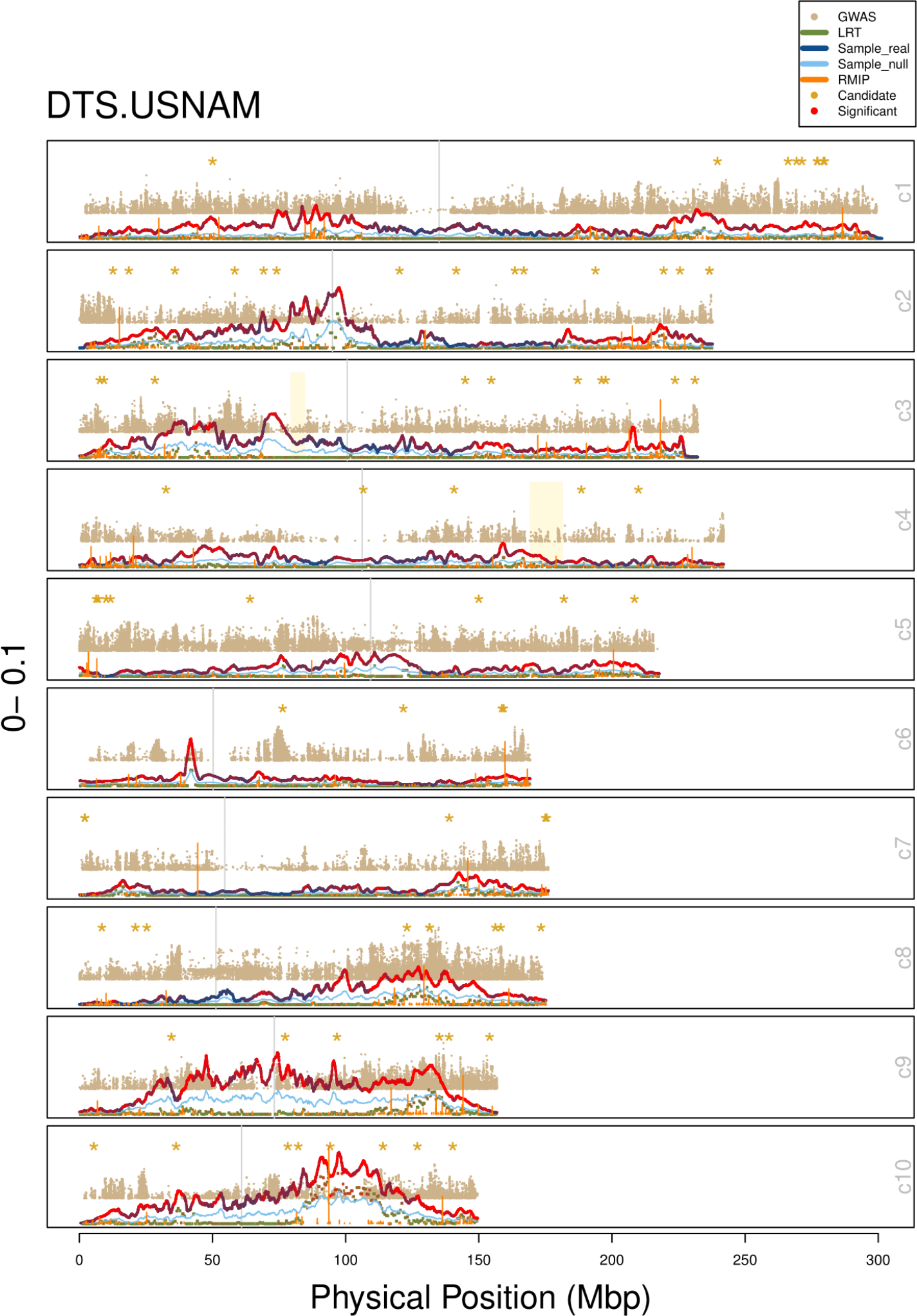
Combined results for all methods (GWAS (uncorrected), resampling GWAS (RMIP), RHM, and BRHM) for USNAM, DTS. RHM and BRHM variance estimates are unmodified. GWAS–log10 p-values and the proportion of models for which a SNP was chosen in resampling GWAS rescaled to the maximum value of the variances in the window, and only the top 5,000 SNPs across the genome plotted in a window. Both the median estimated heritability for the null phenotypes (light blue) and the real estimated heritability (blue to red) is shown for BRHM. RMIP bars in orange represent the proportion of models in which that SNP was chosen. Candidate genes (based on Dong et al./ Danilevskya et al) are noted with a yellow asterisk and listed across the top.

**Figure S13.**
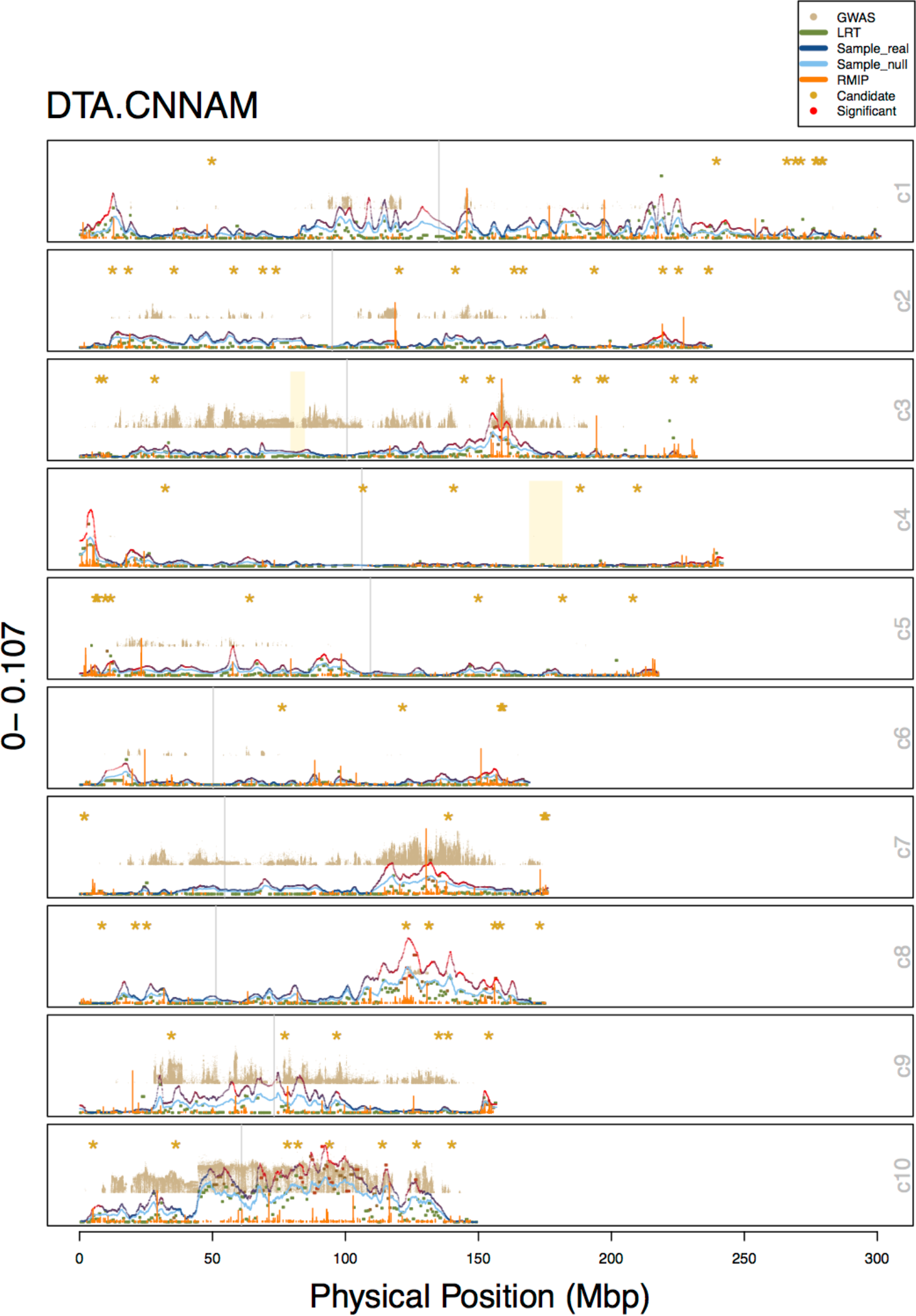
Combined results for all methods (GWAS (uncorrected), resampling GWAS (RMIP), RHM, and BRHM) for CNNAM, DTA. RHM and BRHM variance estimates are unmodified. GWAS–log10 p-values and the proportion of models for which a SNP was chosen in resampling GWAS rescaled to the maximum value of the variances in the window, and only the top 5,000 SNPs across the genome plotted in a window. Both the median estimated heritability for the null phenotypes (light blue) and the real estimated heritability (blue to red) is shown for BRHM. RMIP bars in orange represent the proportion of models in which that SNP was chosen. Candidate genes (based on Dong et al./ Danilevskya et al) are noted with a yellow asterisk and listed across the top.

**Figure S14.**
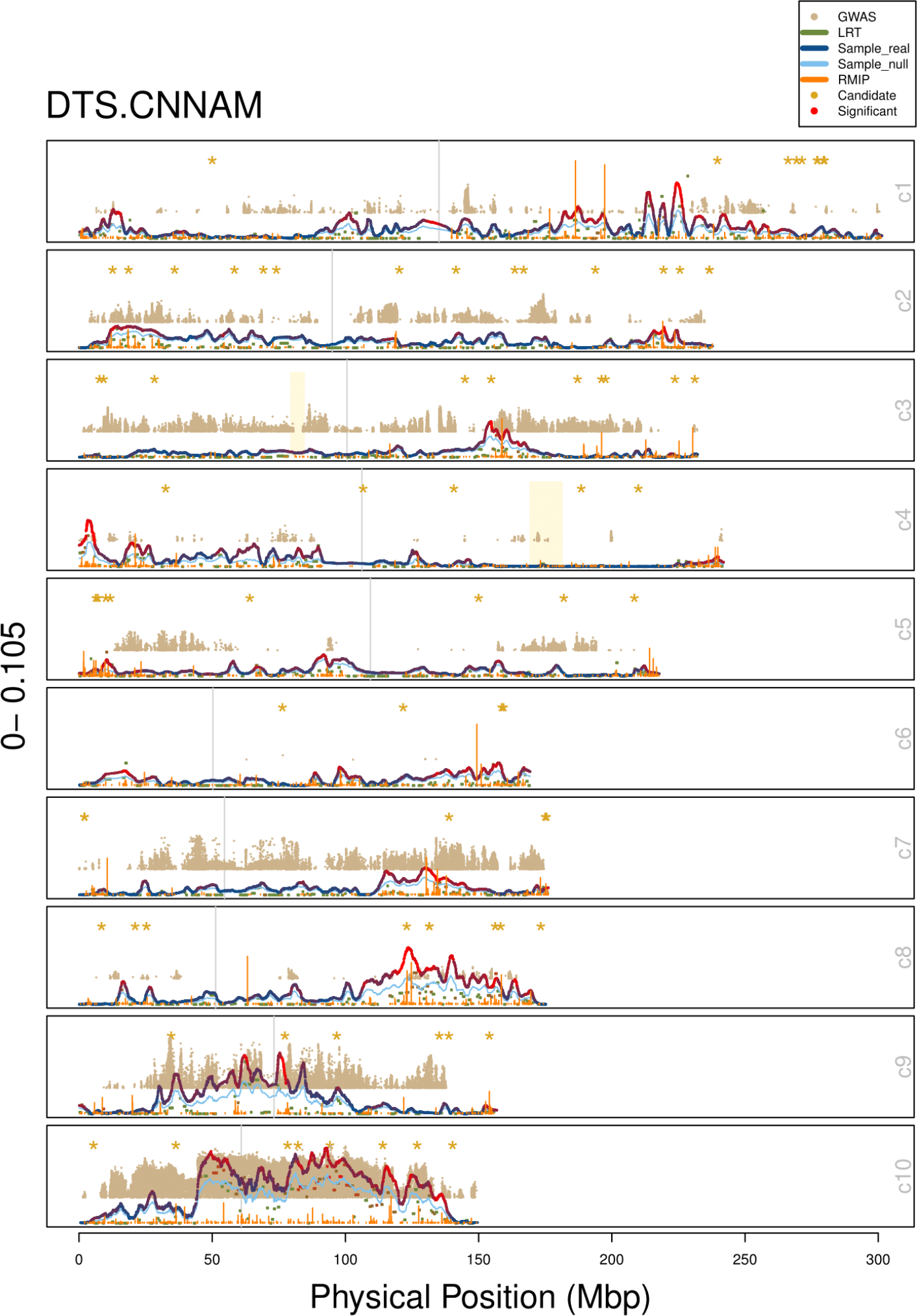
Combined results for all methods (GWAS (uncorrected), resampling GWAS (RMIP), RHM, and BRHM) for CNNAM, DTS. RHM and BRHM variance estimates are unmodified. GWAS–log10 p-values and the proportion of models for which a SNP was chosen in resampling GWAS rescaled to the maximum value of the variances in the window, and only the top 5,000 SNPs across the genome plotted in a window. Both the median estimated heritability for the null phenotypes (light blue) and the real estimated heritability (blue to red) is shown for BRHM. RMIP bars in orange represent the proportion of models in which that SNP was chosen. Candidate genes (based on Dong et al./ Danilevskya et al) are noted with a yellow asterisk and listed across the top.

**Figure S15.**
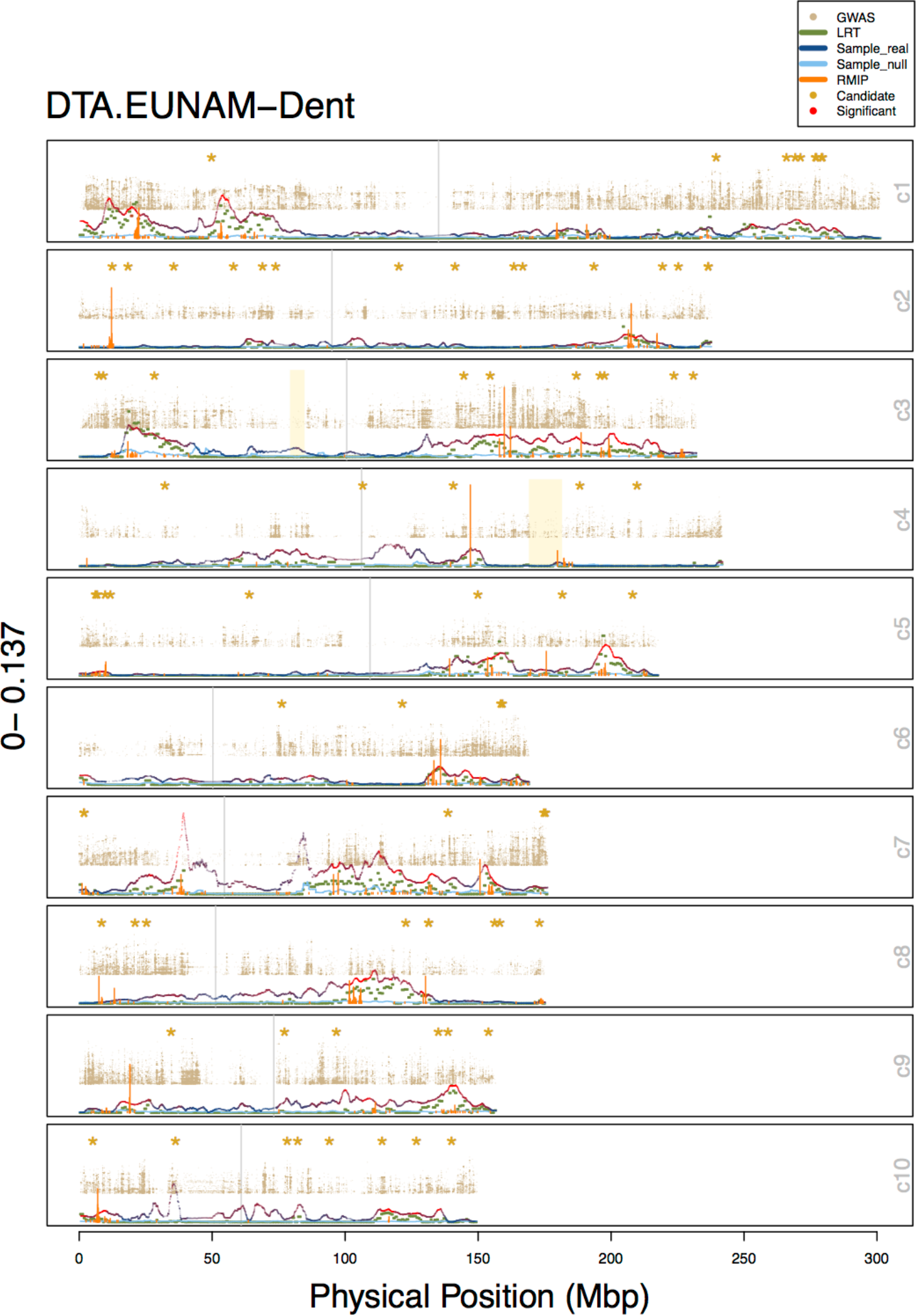
Combined results for all methods (GWAS (uncorrected), resampling GWAS (RMIP), RHM, and BRHM) for EUNAM-Dent, DTA. RHM and BRHM variance estimates are unmodified. GWAS–log10 p-values and the proportion of models for which a SNP was chosen in resampling GWAS rescaled to the maximum value of the variances in the window, and only the top 5,000 SNPs across the genome plotted in a window. Both the median estimated heritability for the null phenotypes (light blue) and the real estimated heritability (blue to red) is shown for BRHM. RMIP bars in orange represent the proportion of models in which that SNP was chosen. Candidate genes (based on Dong et al./ Danilevskya et al) are noted with a yellow asterisk and listed across the top.

**Figure S16.**
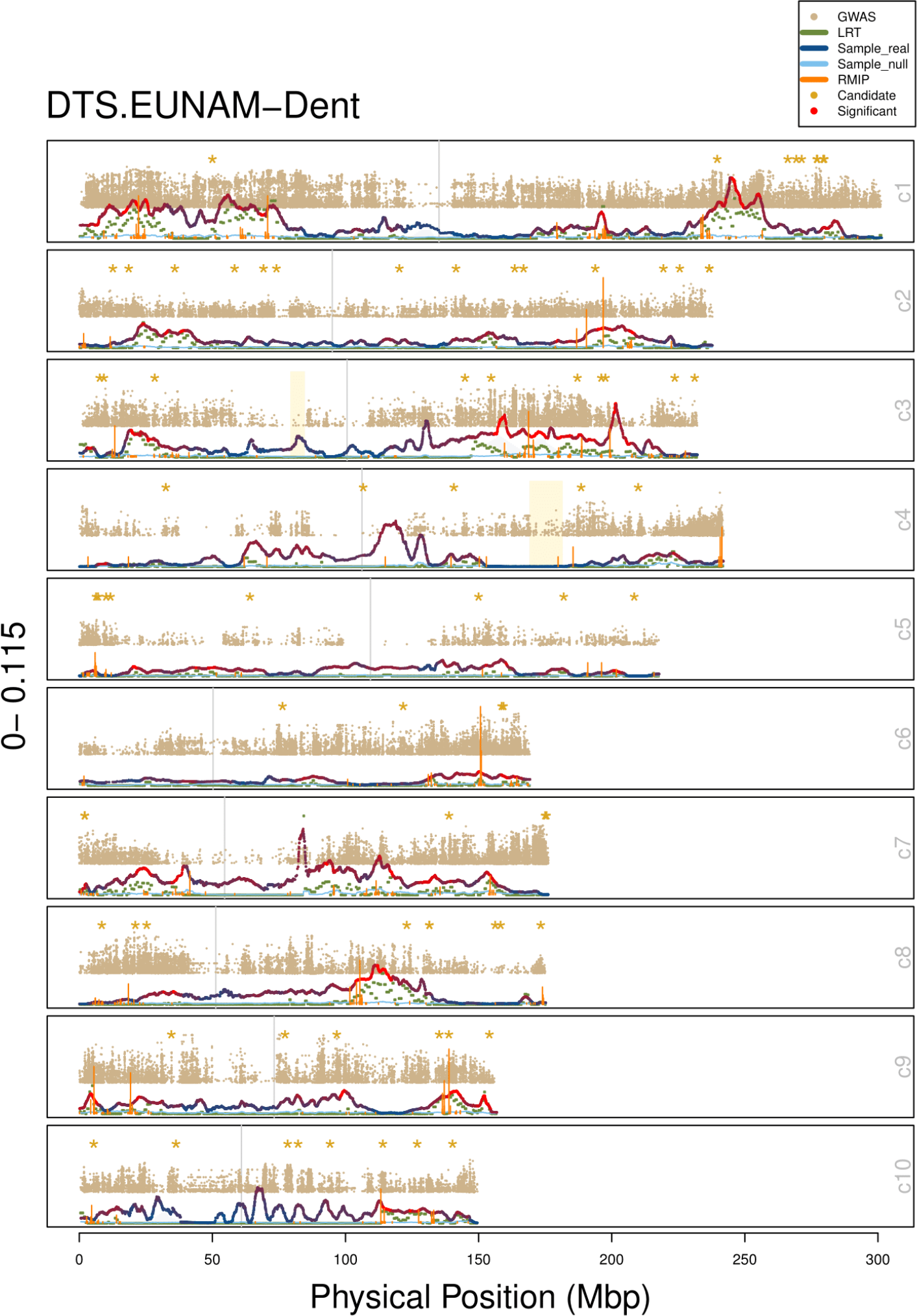
Combined results for all methods (GWAS (uncorrected), resampling GWAS (RMIP), RHM, and BRHM) for EUNAM-Dent, DTS. RHM and BRHM variance estimates are unmodified. GWAS–log10 p-values and the proportion of models for which a SNP was chosen in resampling GWAS rescaled to the maximum value of the variances in the window, and only the top 5,000 SNPs across the genome plotted in a window. Both the median estimated heritability for the null phenotypes (light blue) and the real estimated heritability (blue to red) is shown for BRHM. RMIP bars in orange represent the proportion of models in which that SNP was chosen. Candidate genes (based on Dong et al./ Danilevskya et al) are noted with a yellow asterisk and listed across the top.

**Figure S17.**
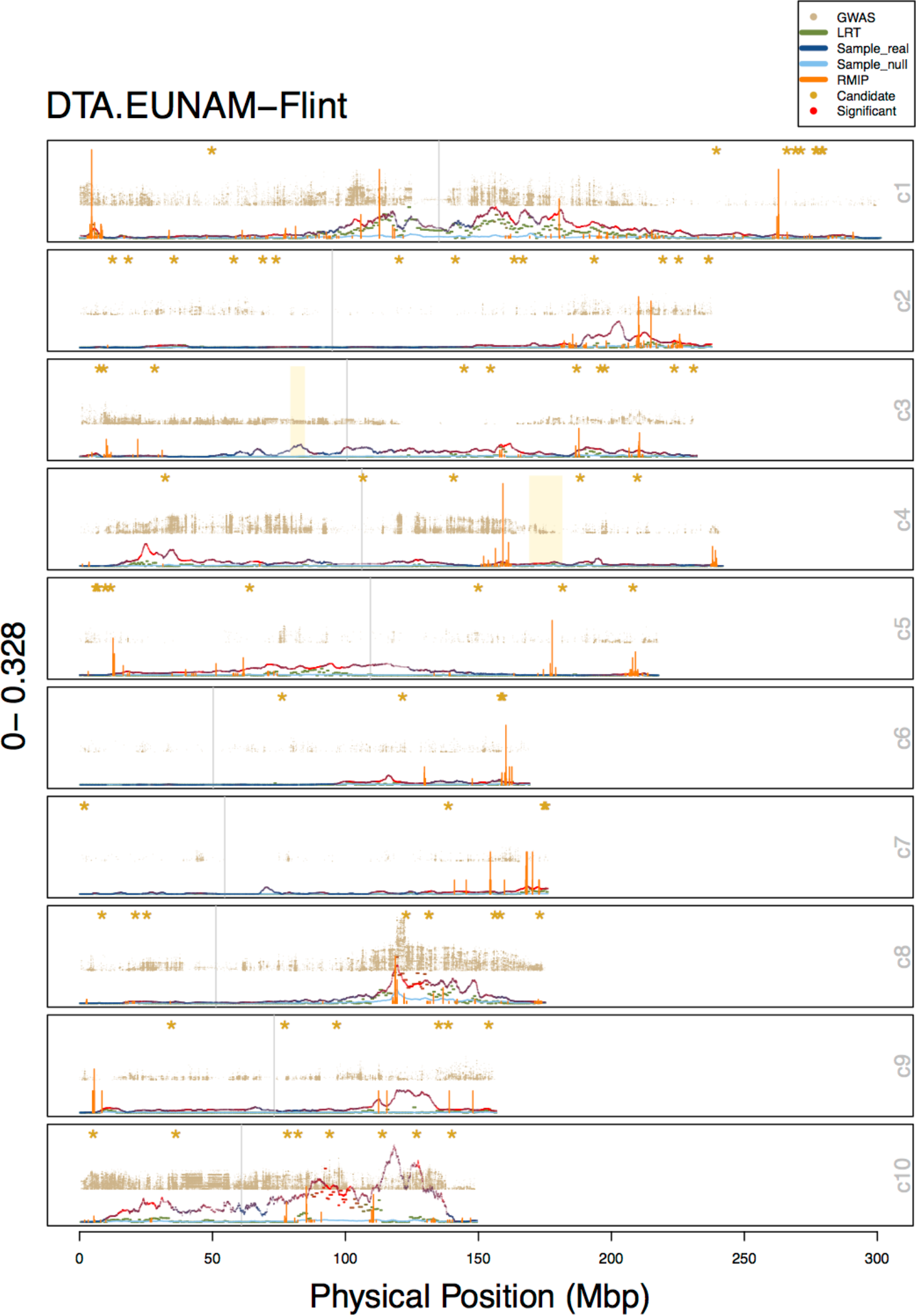
Combined results for all methods (GWAS (uncorrected), resampling GWAS RMIP), RHM, and BRHM) for EUNAM-Flint, DTA. RHM and BRHM variance estimates are unmodified. GWAS–log10 p-values and the proportion of models for which a SNP was chosen in resampling GWAS rescaled to the maximum value of the variances in the window, and only the top 5,000 SNPs across the genome plotted in a window. Both the median estimated heritability for the null phenotypes (light blue) and the real estimated heritability (blue to red) is shown for BRHM. RMIP bars in orange represent the proportion of models in which that SNP was chosen. Candidate genes based on Dong et al./ Danilevskya et al) are noted with a yellow asterisk and listed across the top.

**Figure S18.**
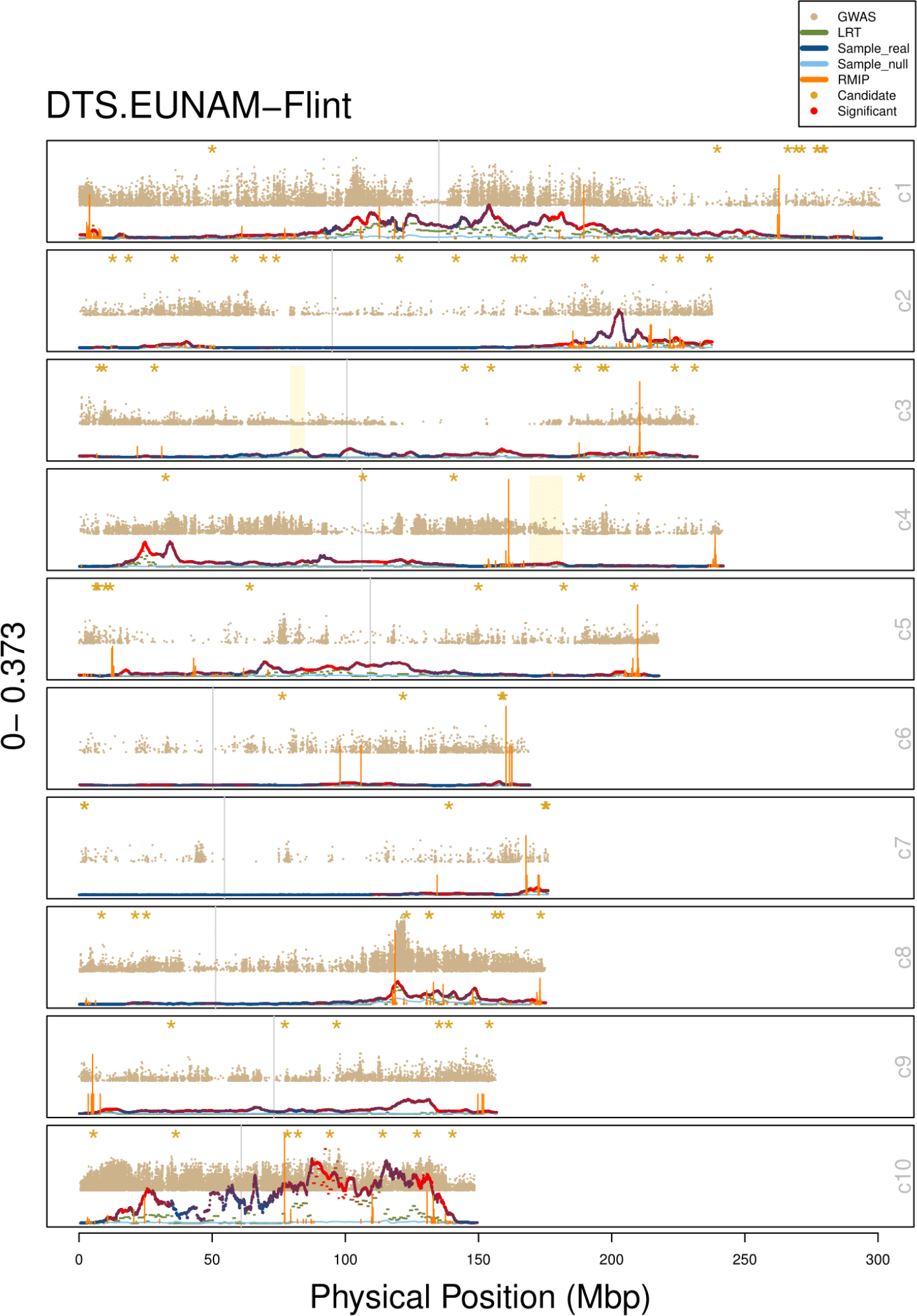
Combined results for all methods (GWAS (uncorrected), resampling GWAS (RMIP), RHM, and BRHM) for EUNAM-Flint, DTS. RHM and BRHM variance estimates are unmodified. GWAS–log10 p-values and the proportion of models for which a SNP was chosen in resampling GWAS rescaled to the maximum value of the variances in the window, and only the top 5,000 SNPs across the genome plotted in a window. Both the median estimated heritability for the null phenotypes (light blue) and the real estimated heritability (blue to red) is shown for BRHM. RMIP bars in orange represent the proportion of models in which that SNP was chosen. Candidate genes (based on Dong et al./ Danilevskya et al) are noted with a yellow asterisk and listed across the top.

**Figure S19.**
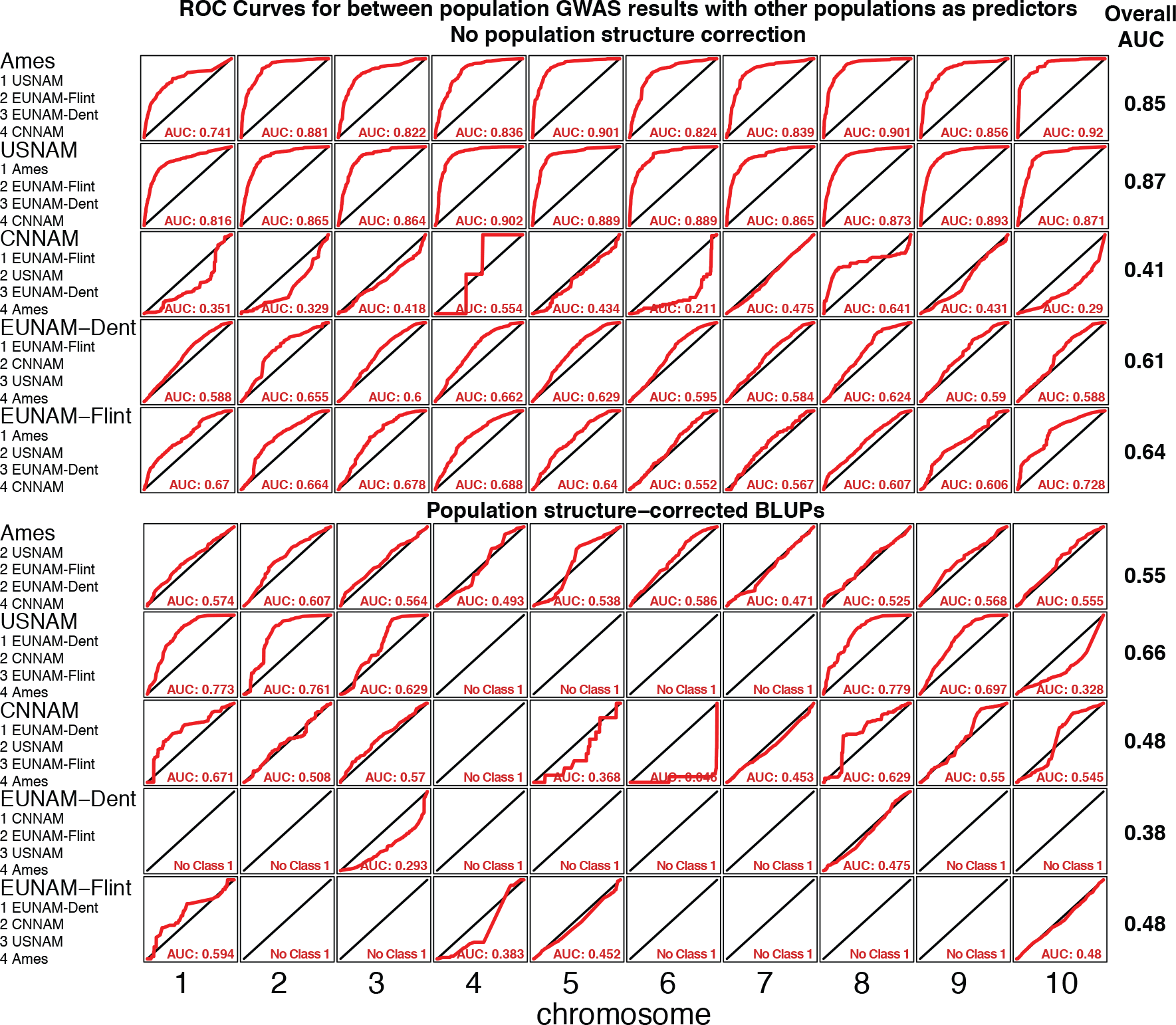
Random forest classifier for GWAS results between populations to evaluate overlap in GWAS results between populations. Predictors are equivalent GWAS (with or without population structure correction) for the other populations. Trained on nine chromosomes and tested on the 10^th^. Not all chromosomes for the population structure corrected results have top predictors, and are excluded from overall AUC calculations.

**Figure S20.**
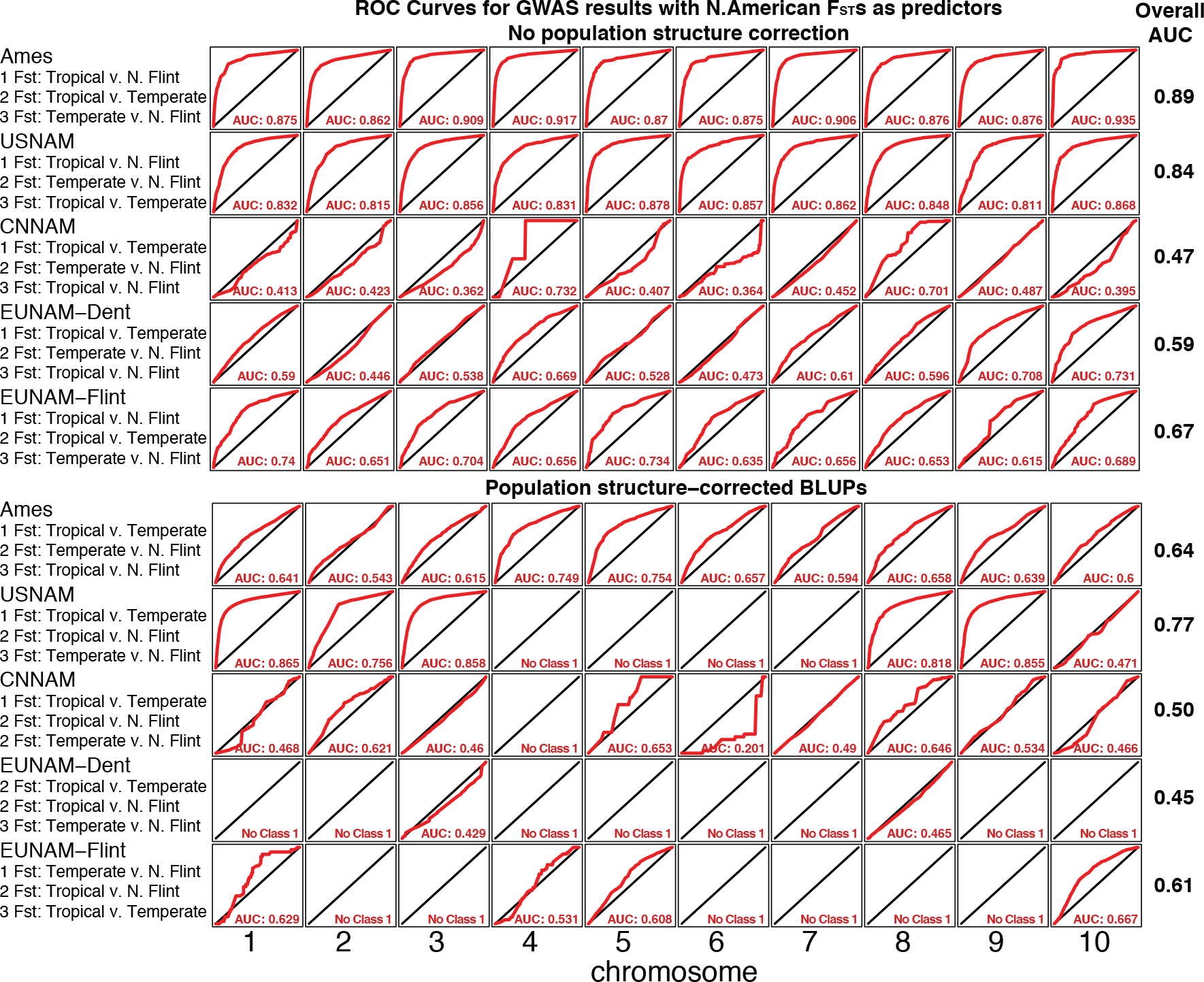
Random forest classifier with GWAS additive p-values as the response, and N. American F_ST_s as the predictors. Trained on nine chromosomes and tested on the 10^th^. Not all chromosomes for the population structure corrected results have top predictors, and are excluded from overall AUC calculations.

**Figure S21.**
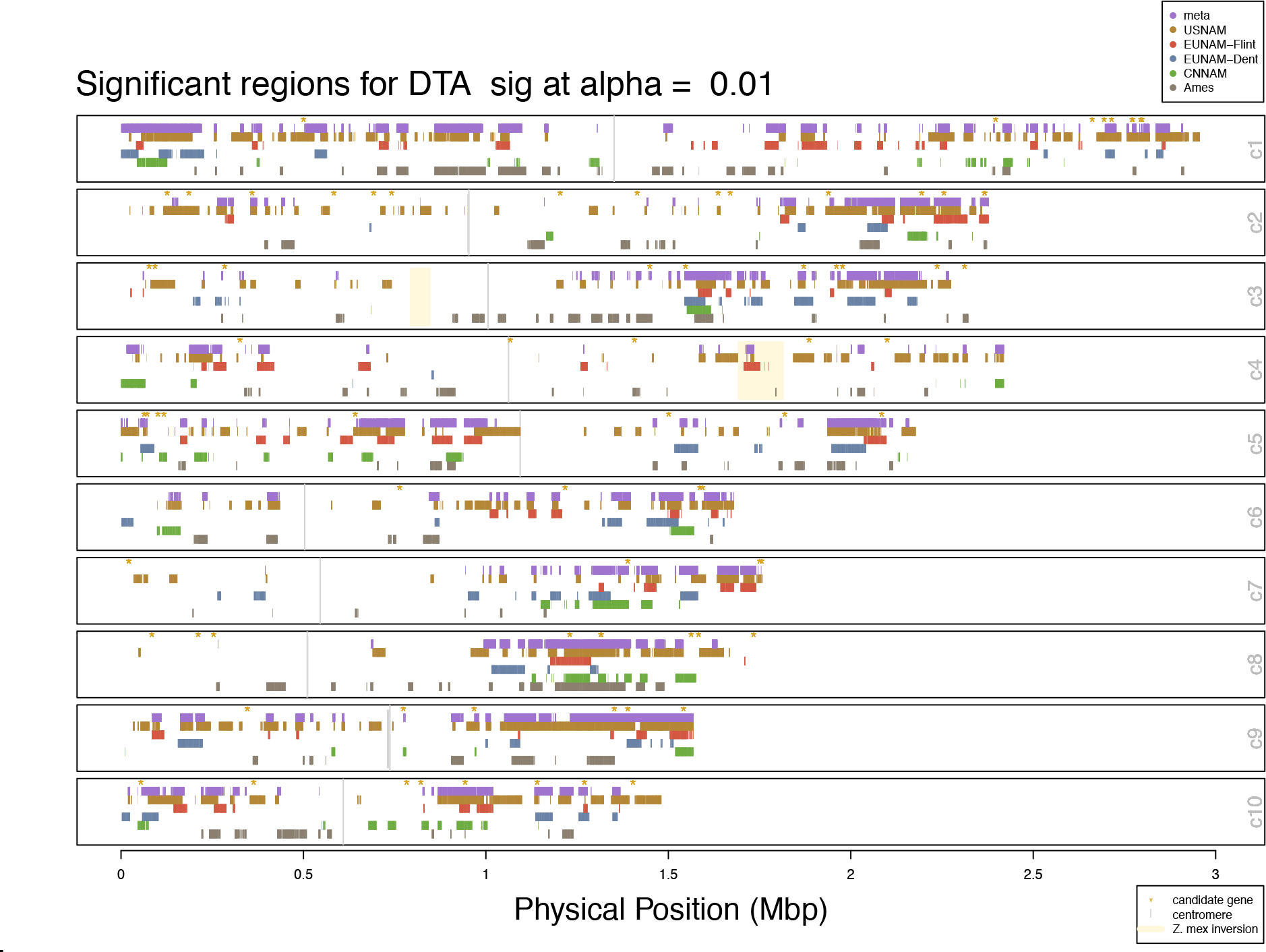
Significant regions and metaanalysis for BRHM results DTA

**Figure S22.**
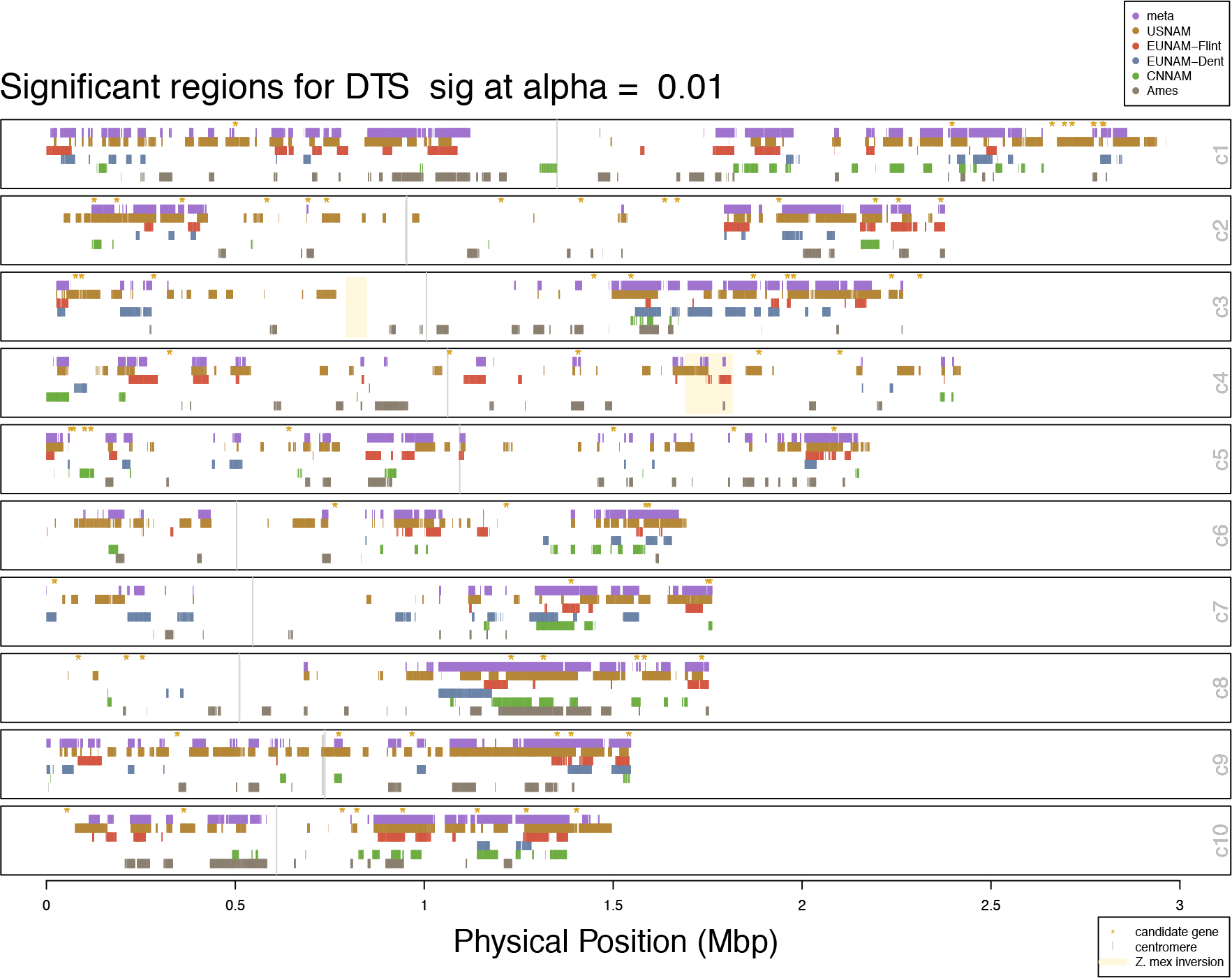
Significant regions and metaanalysis for BRHM results DTS

**Figure S23.**
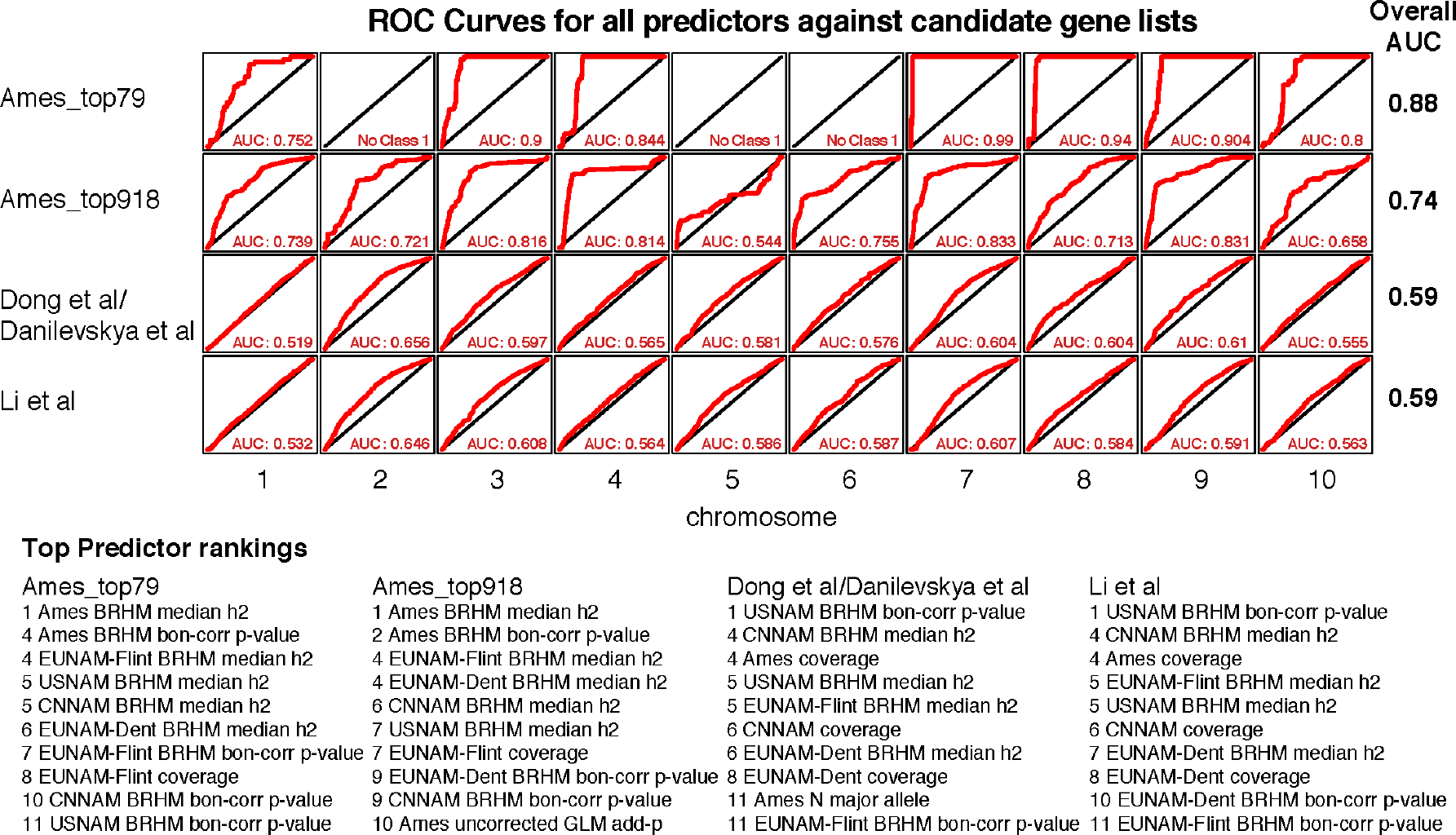
Machine learning using candidate genes as the response, and GWAS and BRHM mapping, power estimators, and F_ST_s as the predictors. Candidate genes were classified as 1, with a 20kbp buffer around the gene start and end position (maize AGP v3). AUCs of 0 result when no candidates are found on the chromosome tested. Not all chromosomes for the Ames top 79 candidates have top predictors, and are excluded from overall AUC calculations.

## Supplemental tables

**Table S1.**
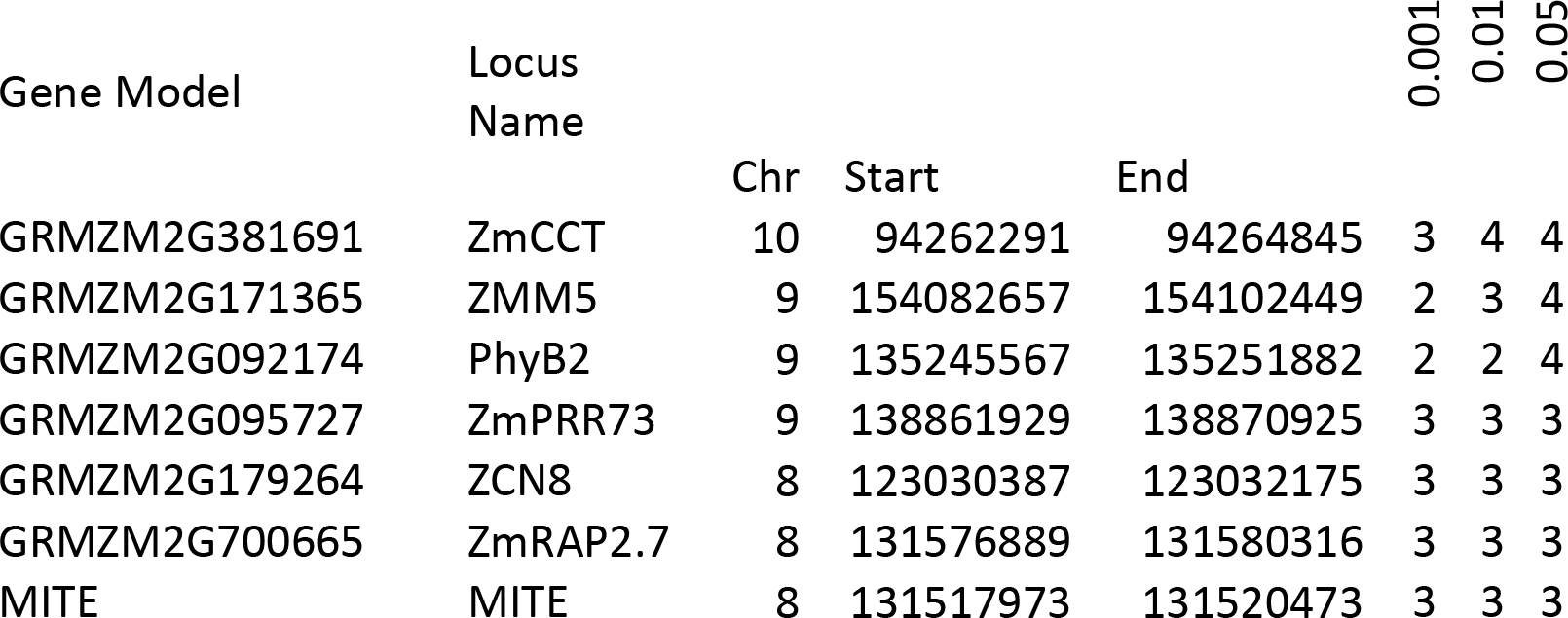
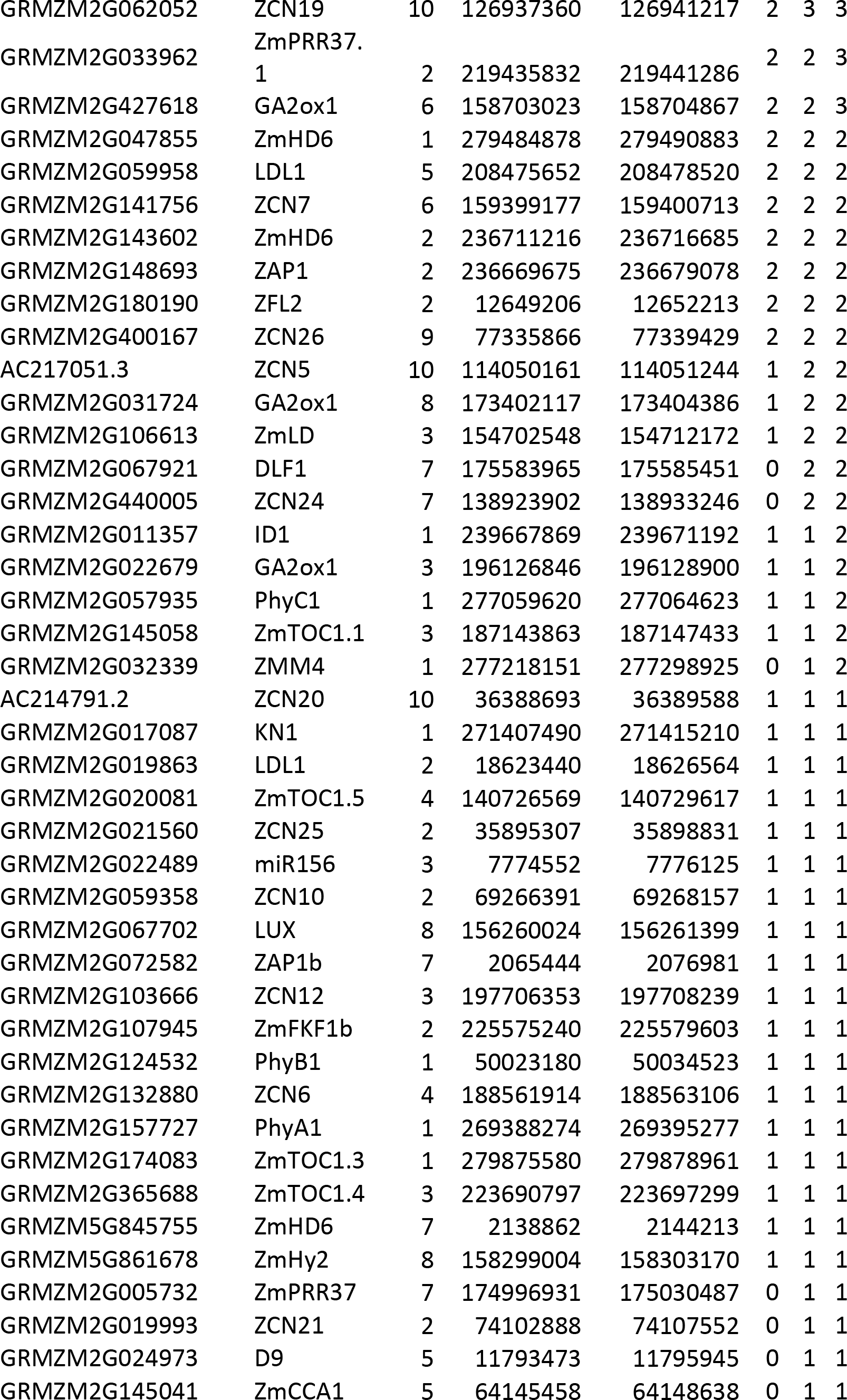
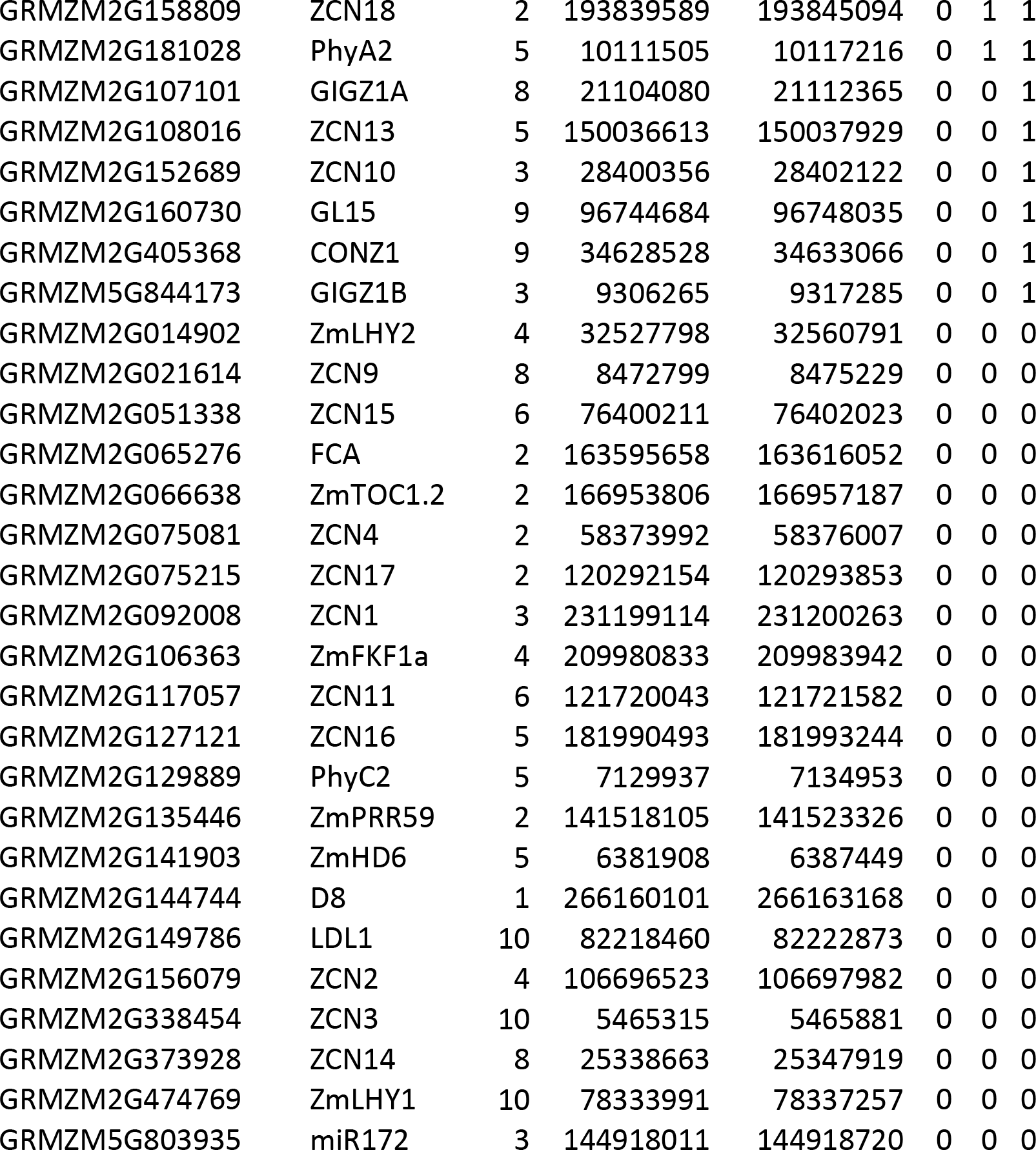
Number of populations significant for autonomous flowering time candidates (after Dong et al) based on Bonferroni corrected significance for BRHM method

**Table S2.**
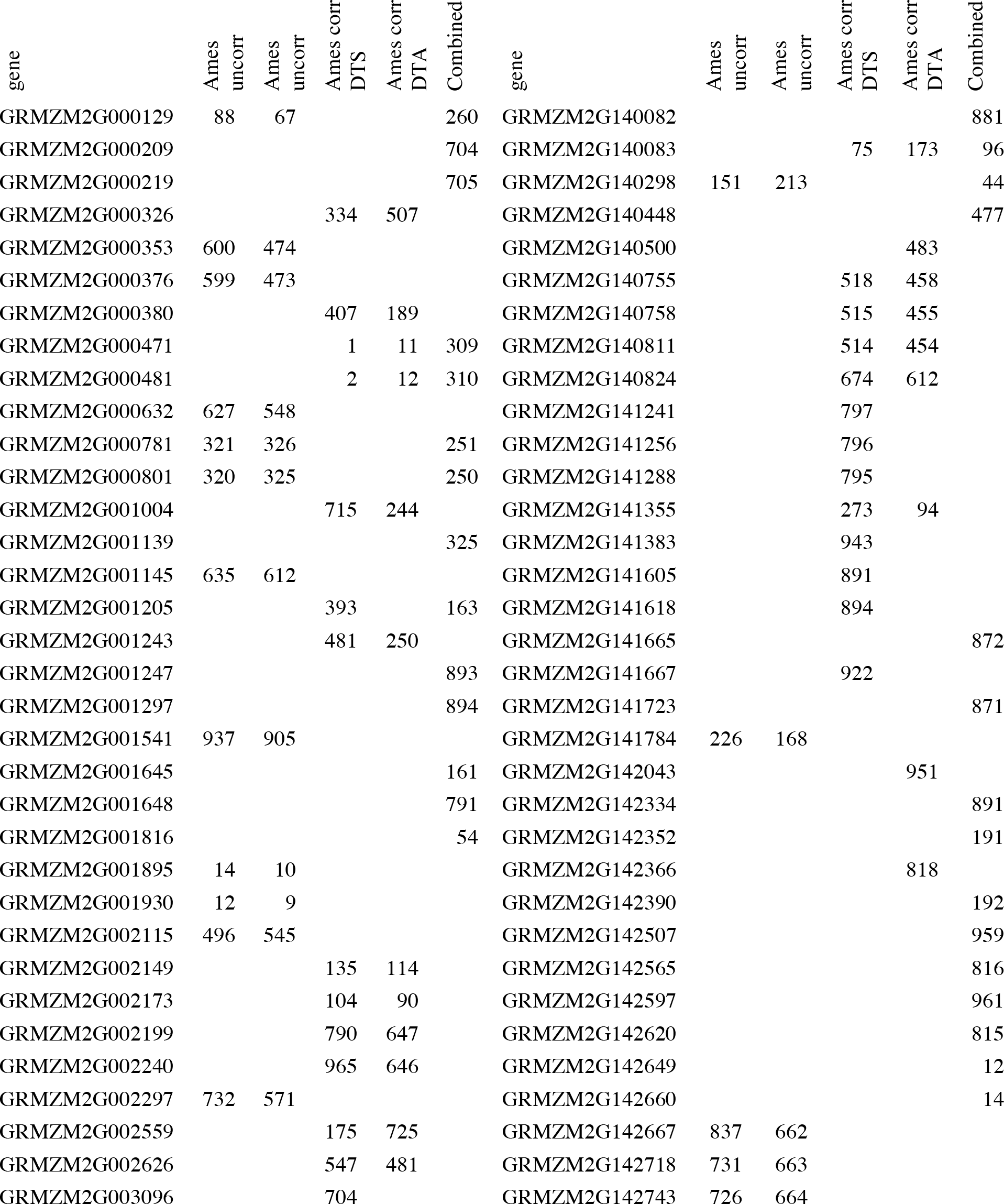
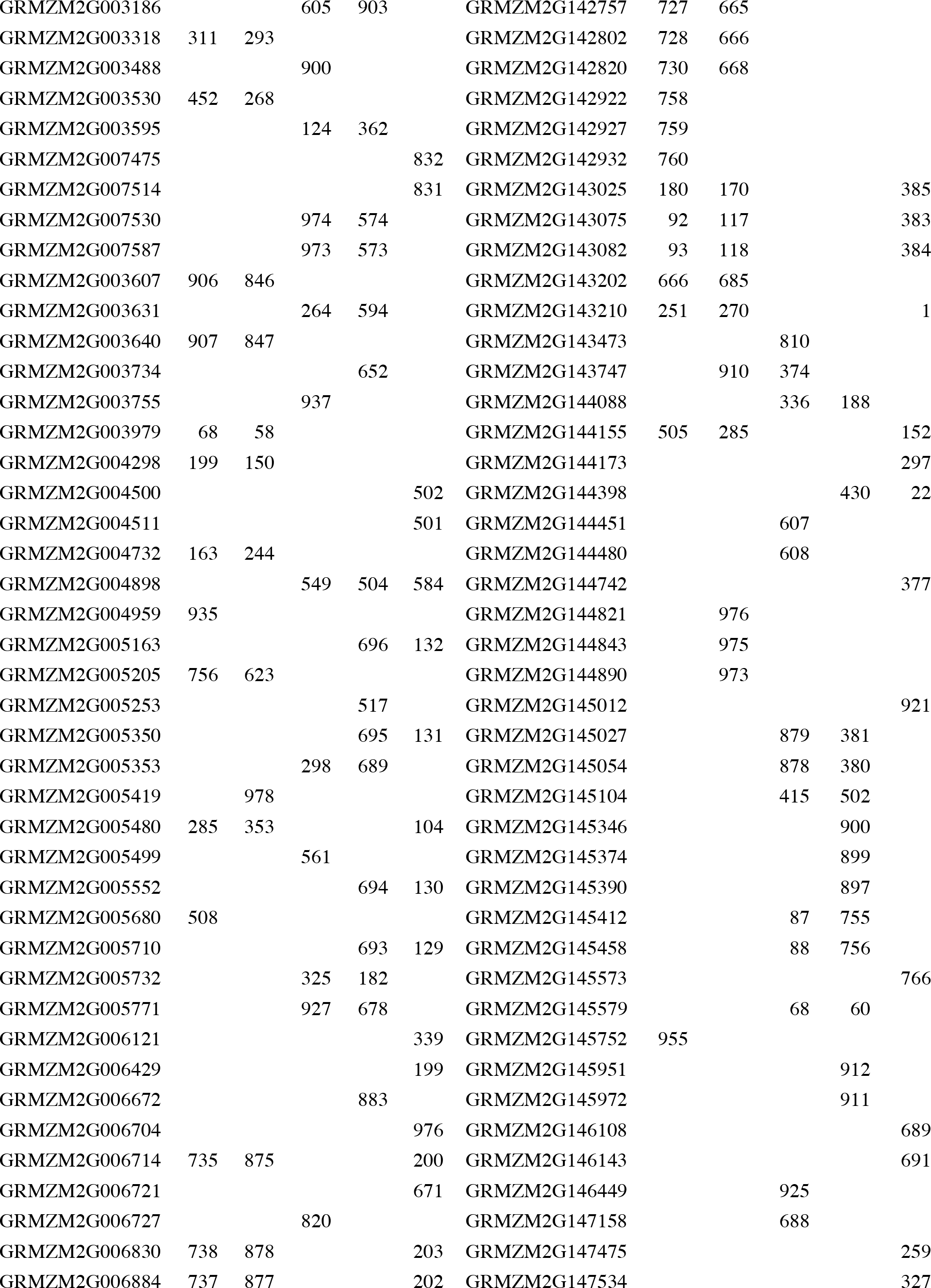
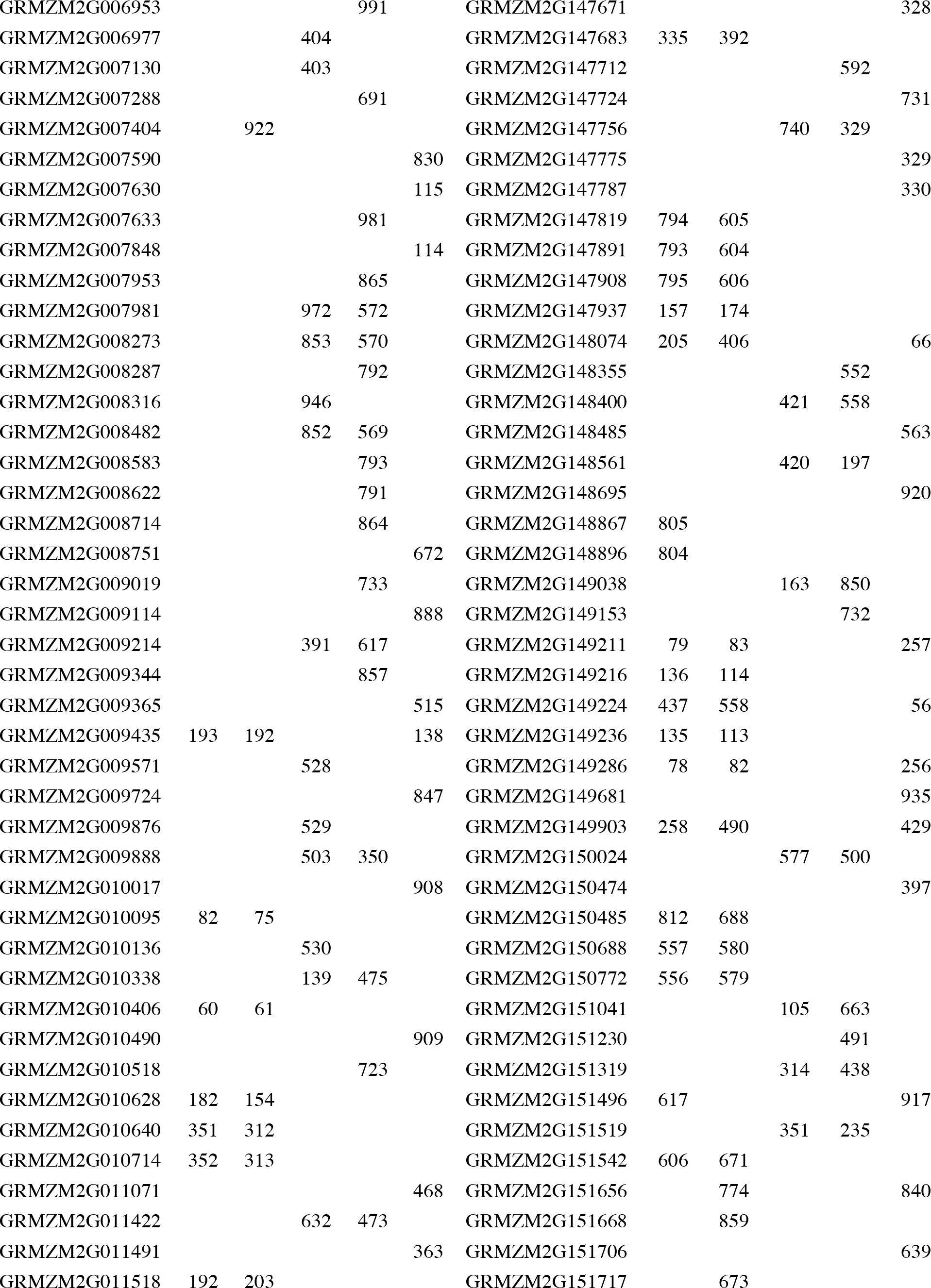
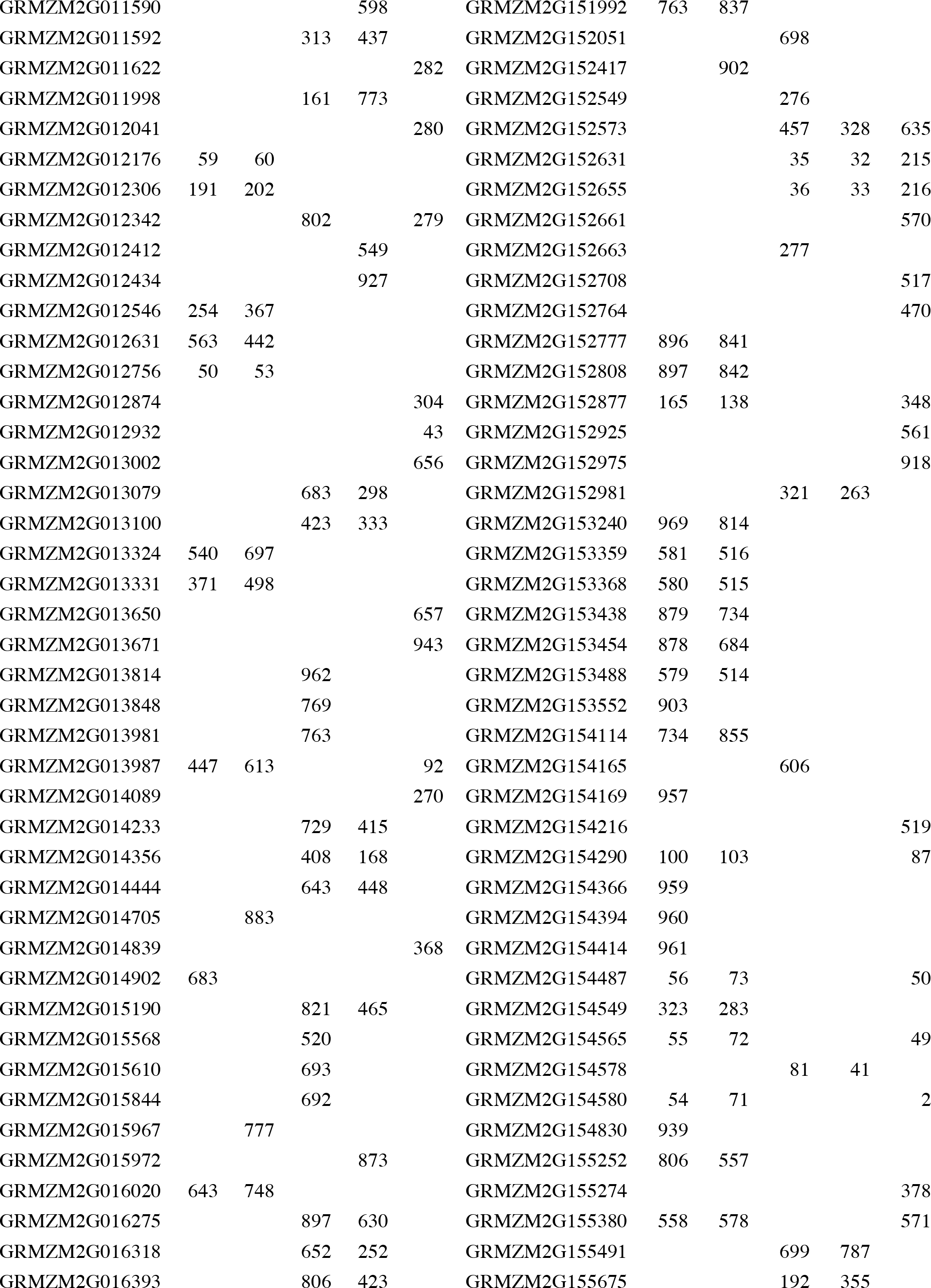
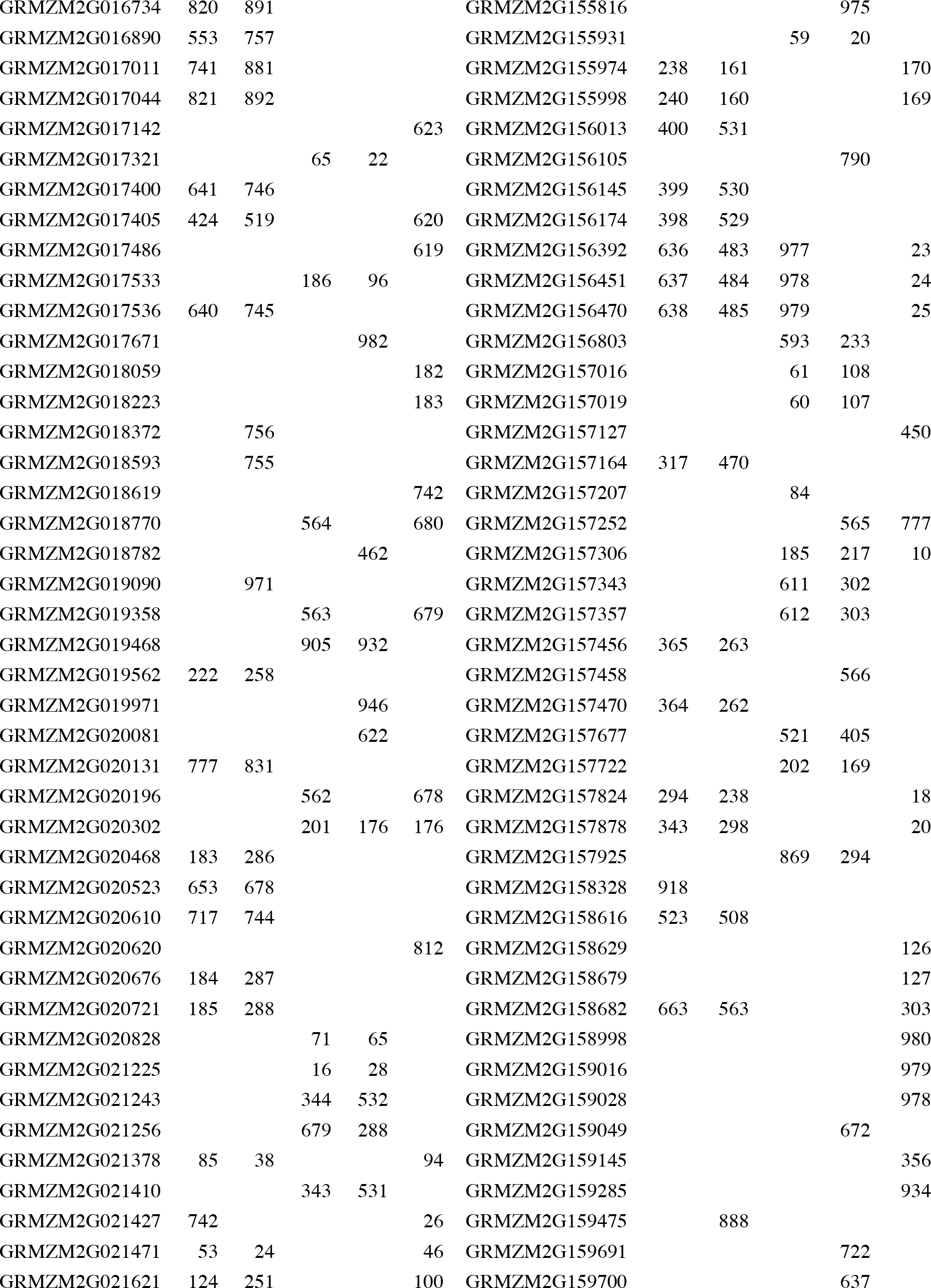
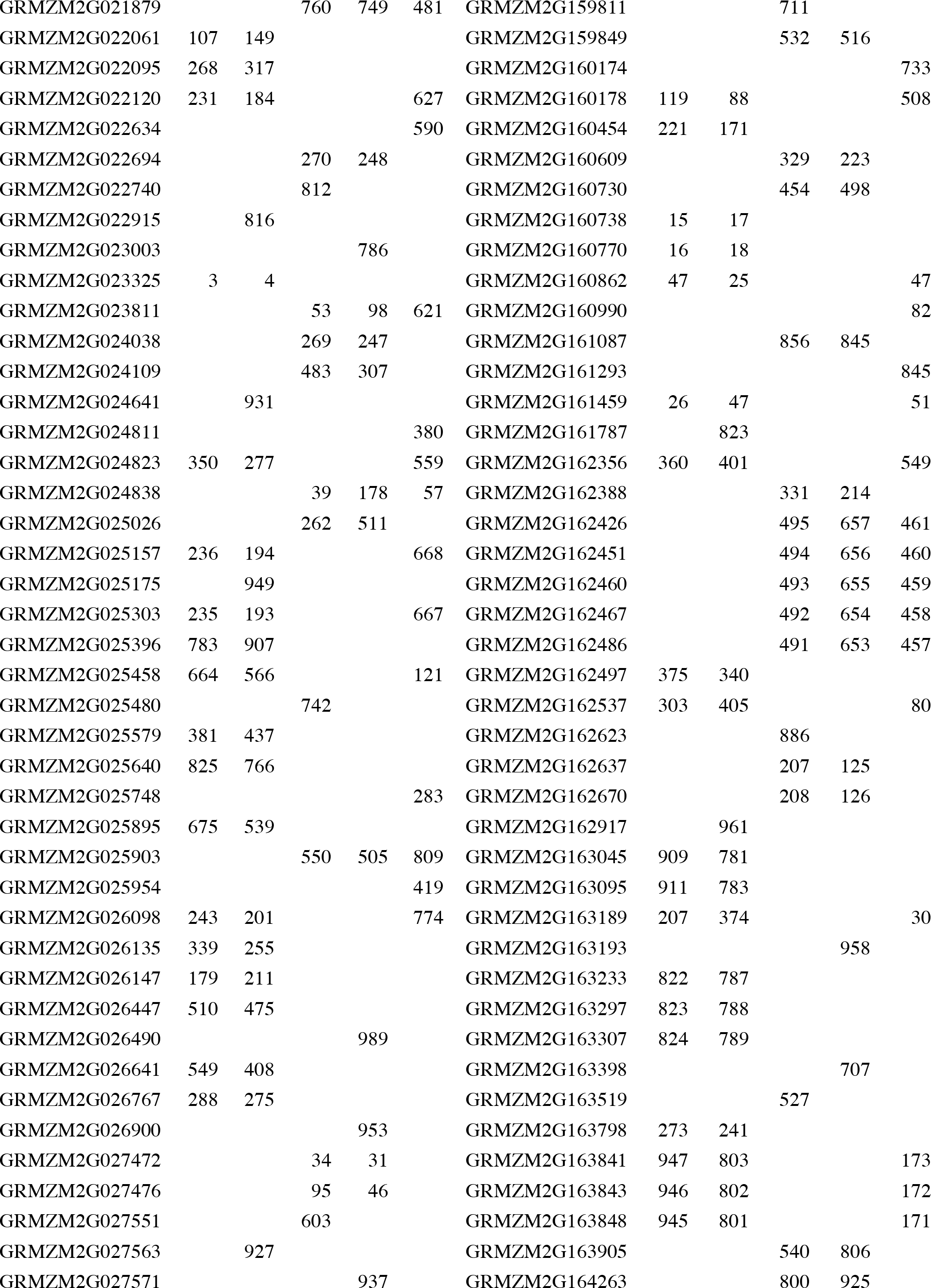
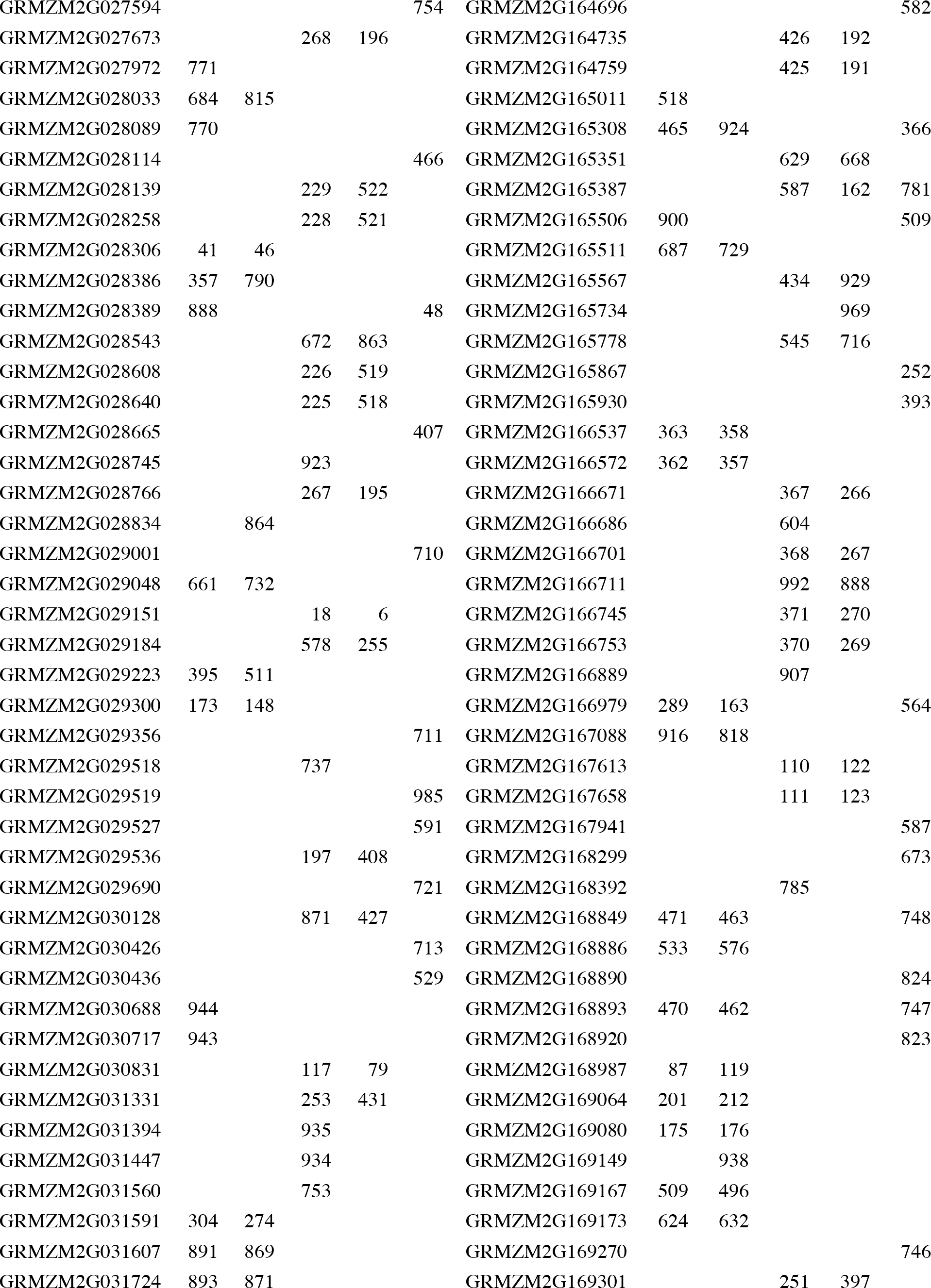
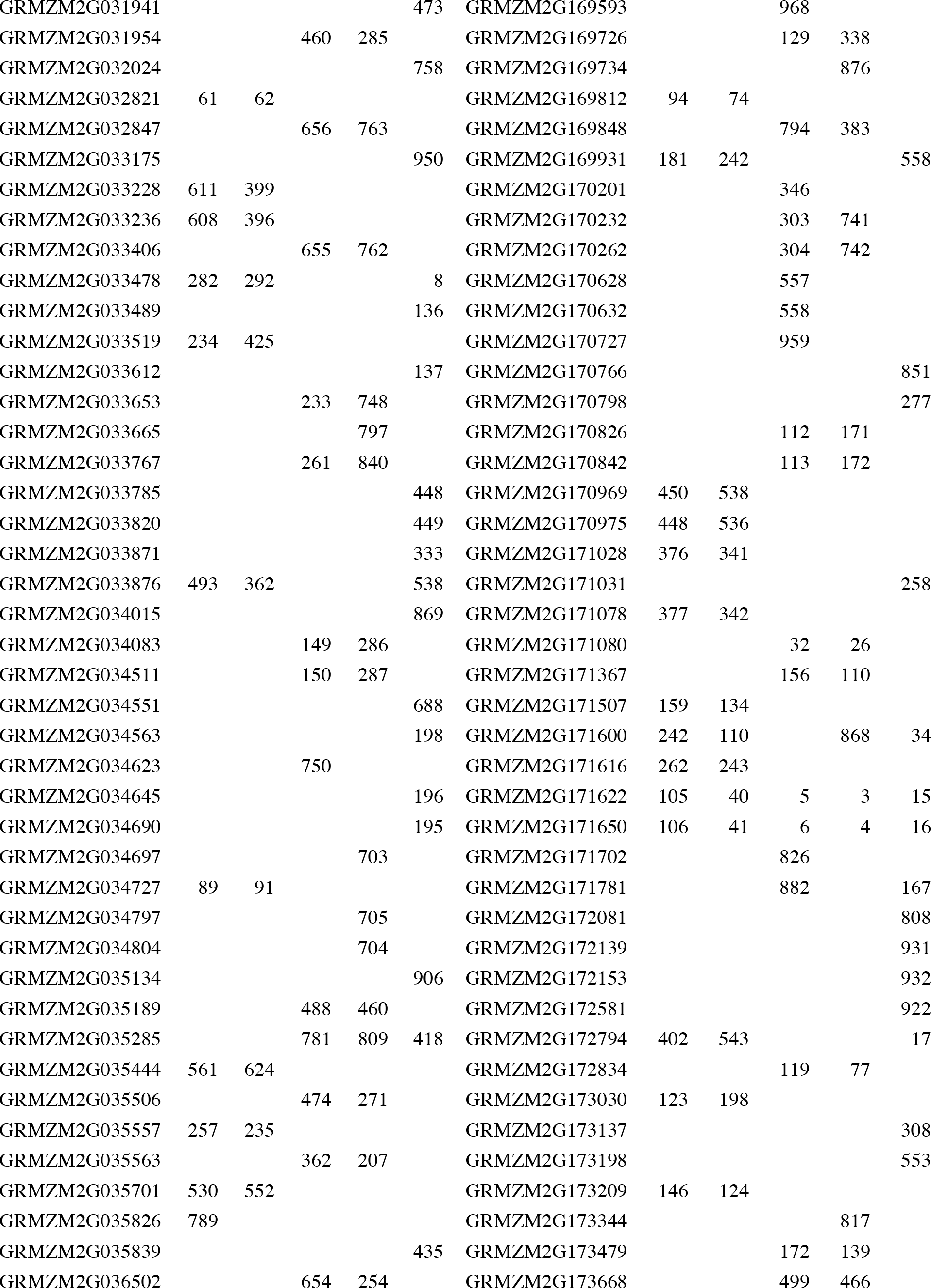
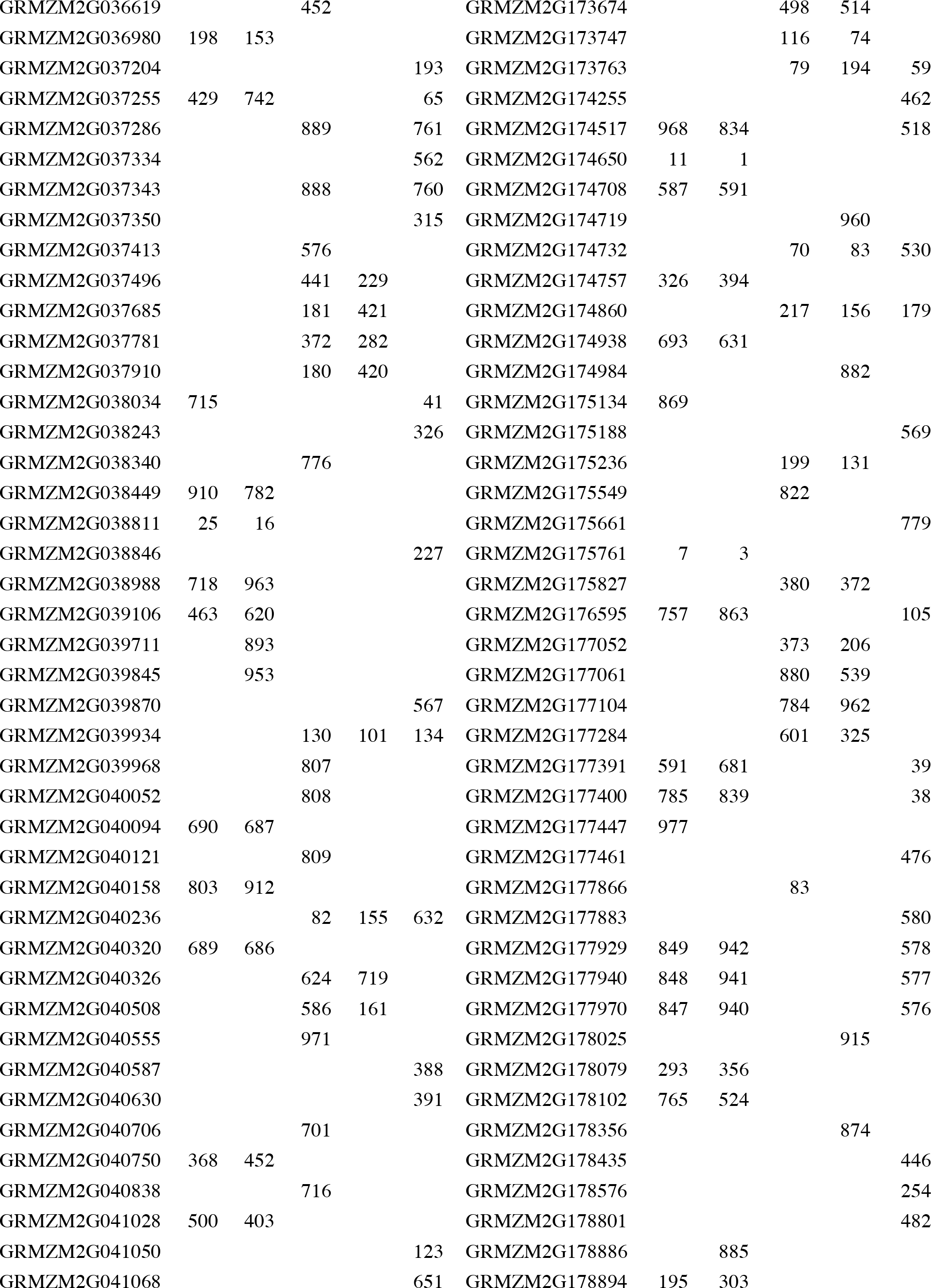
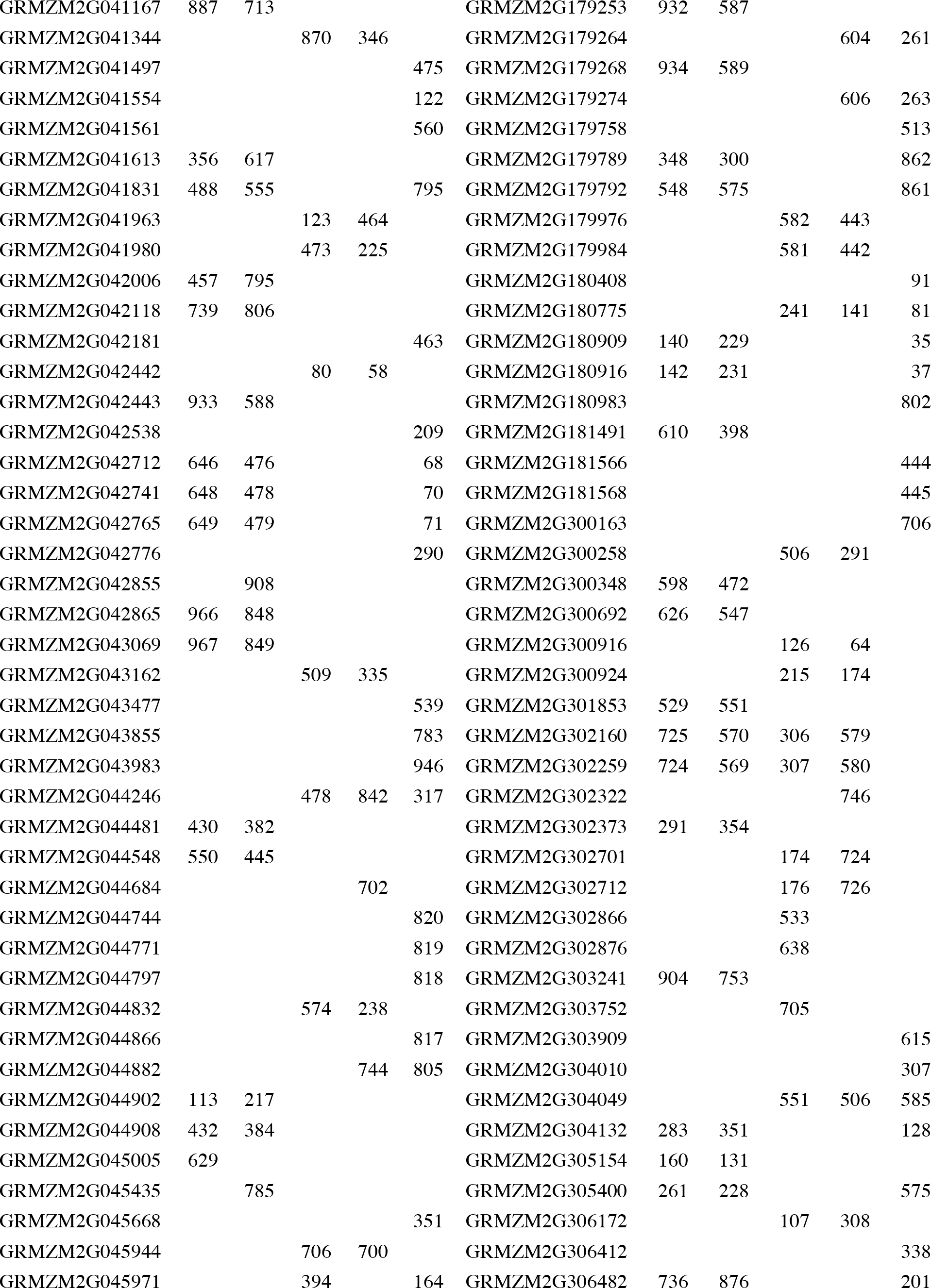
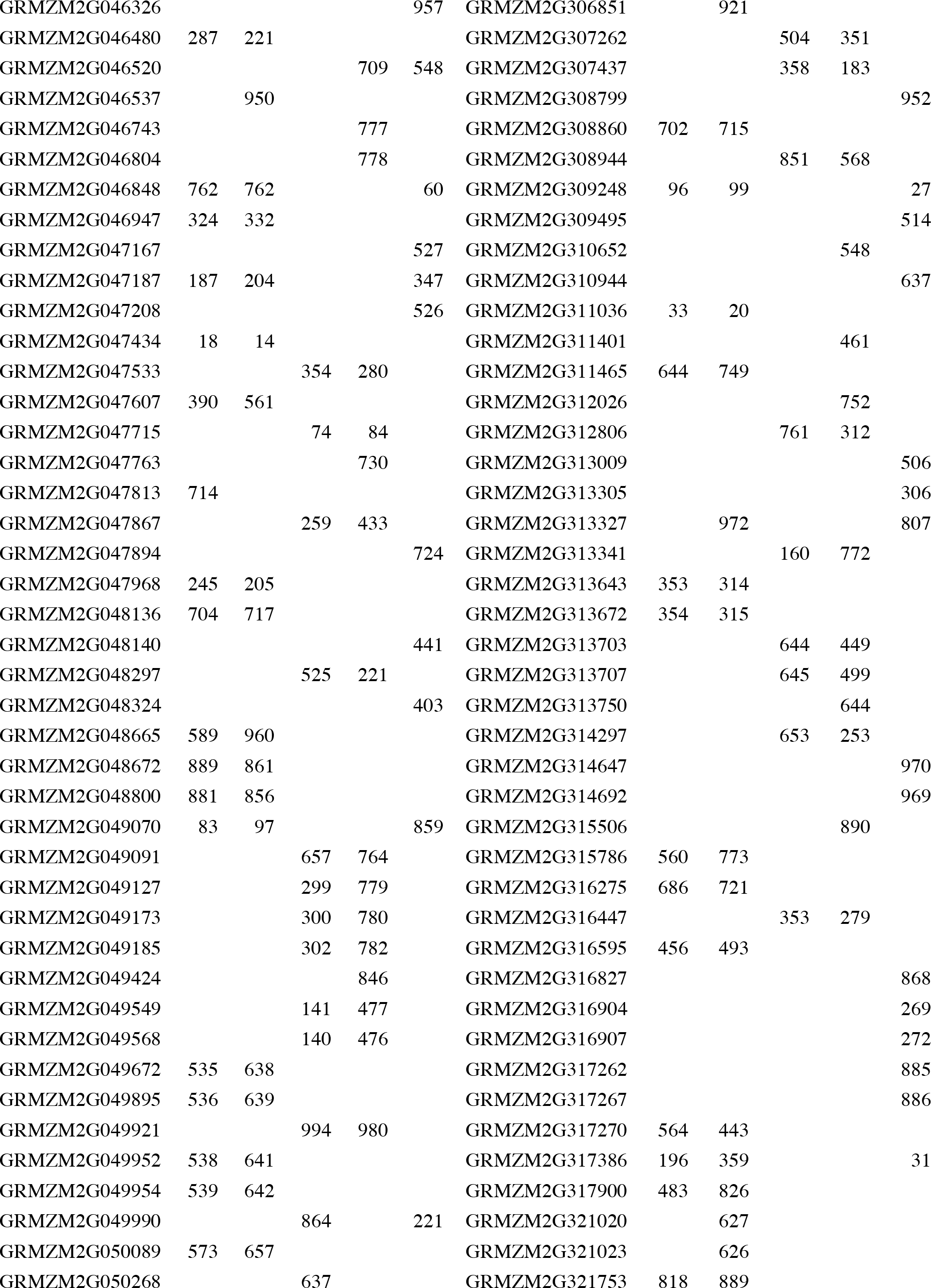
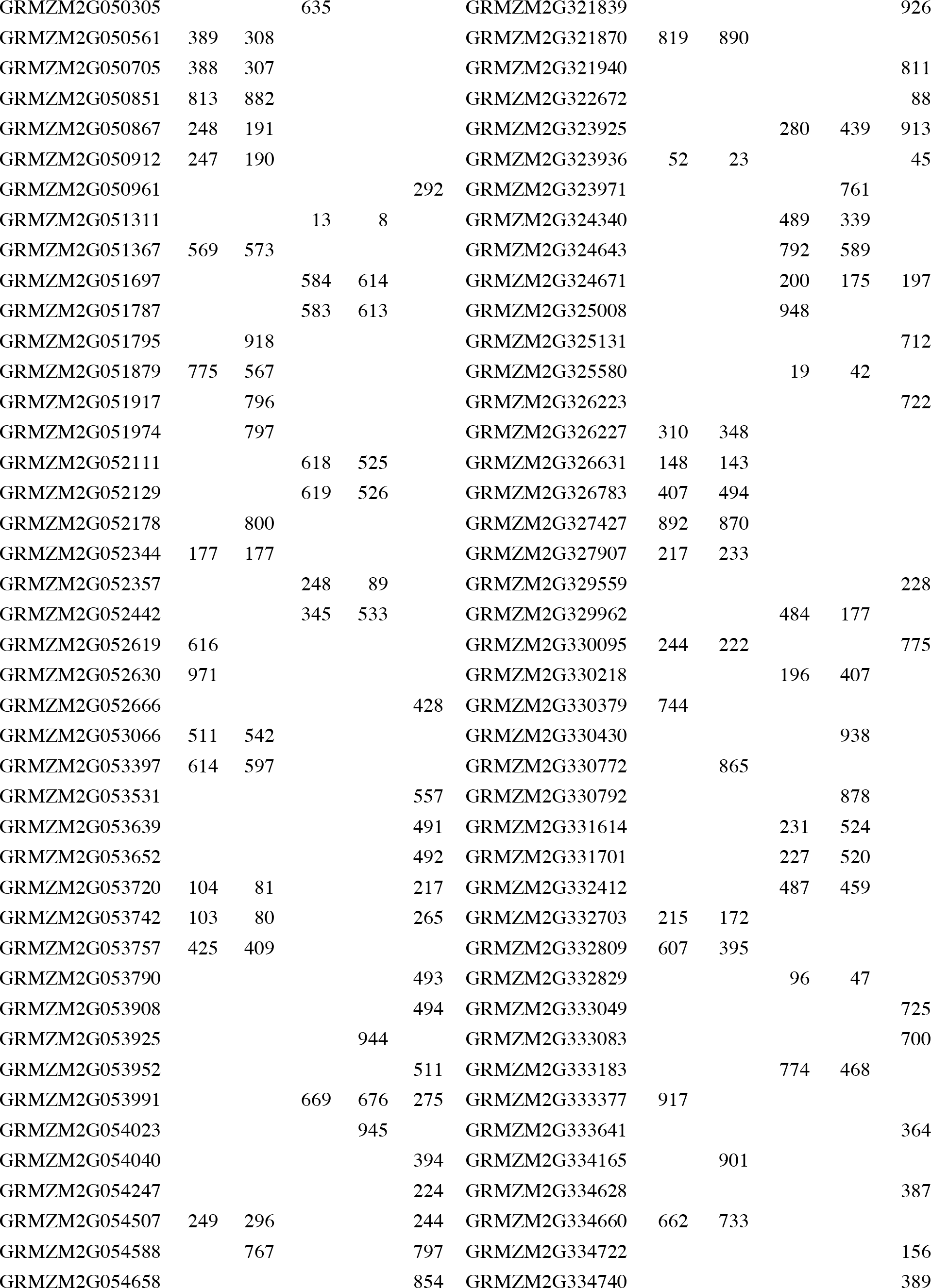
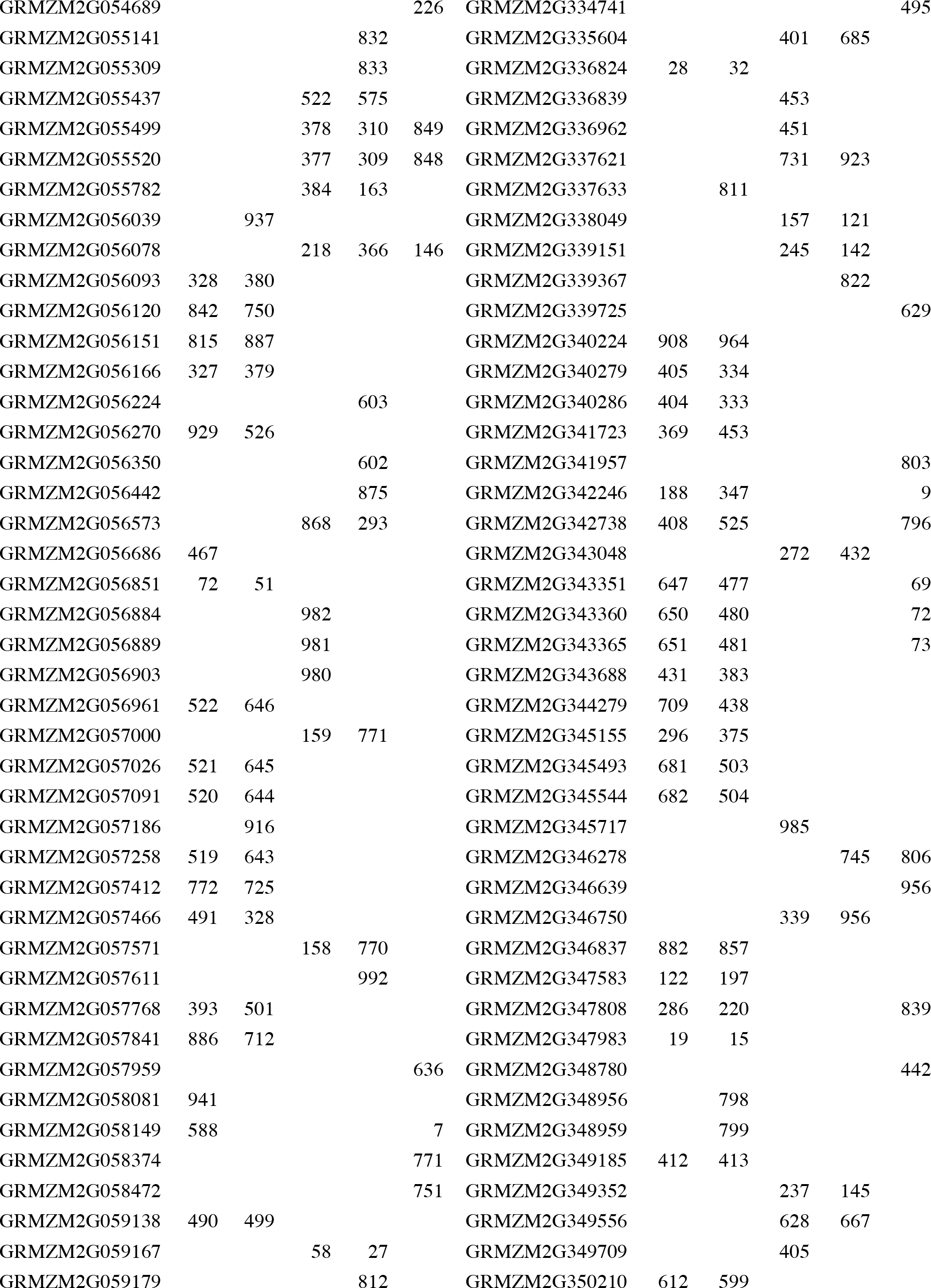
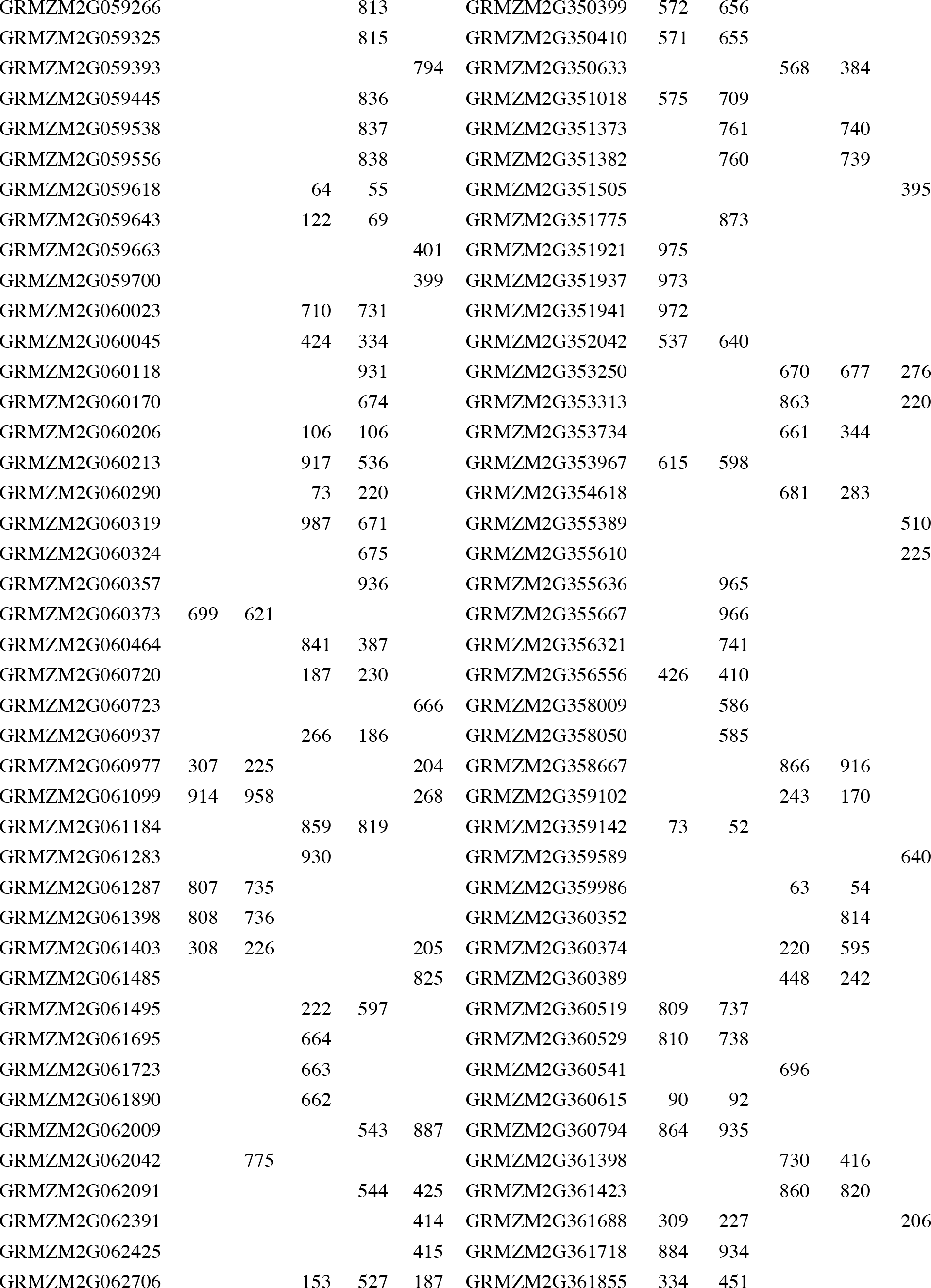
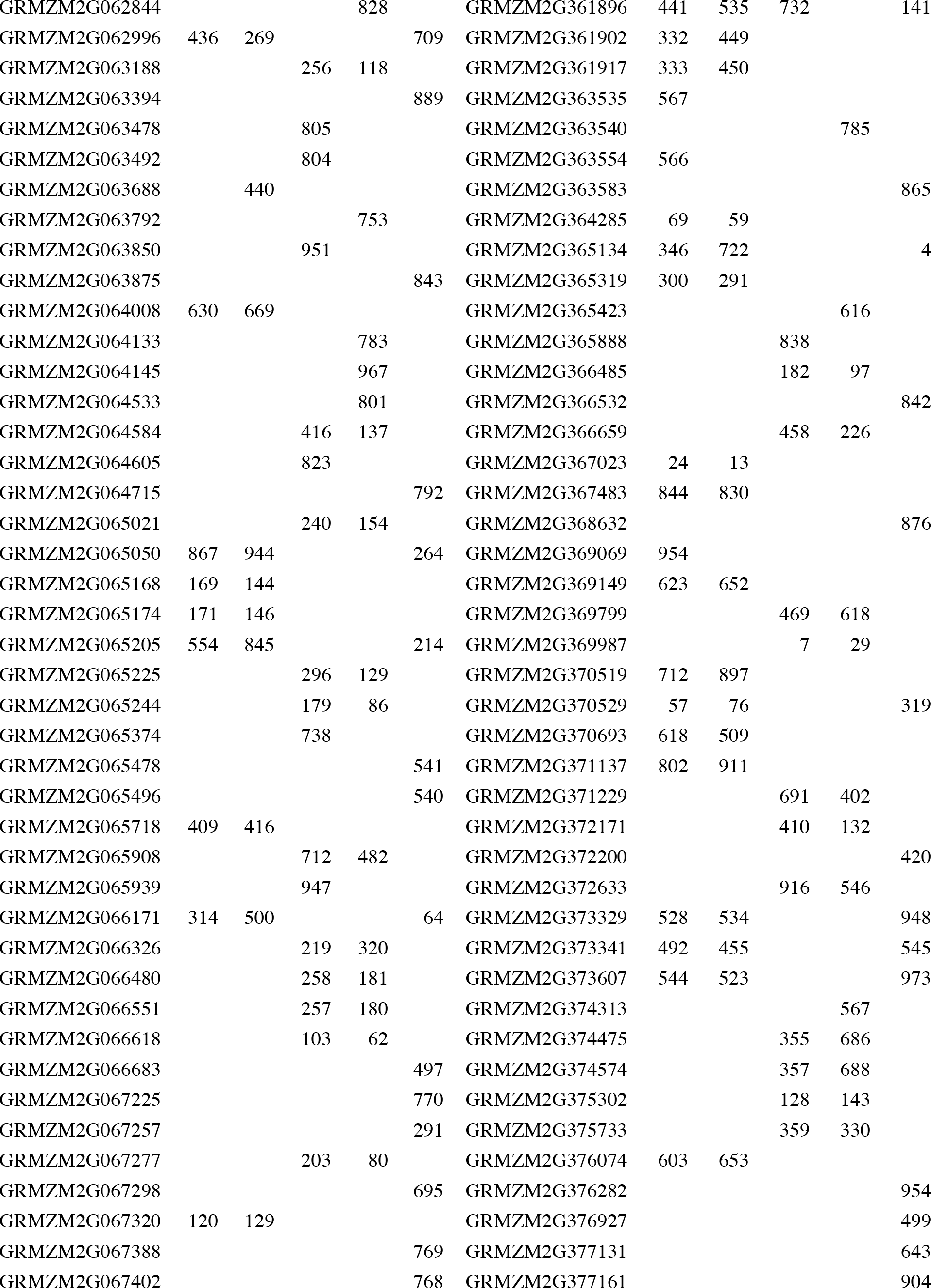
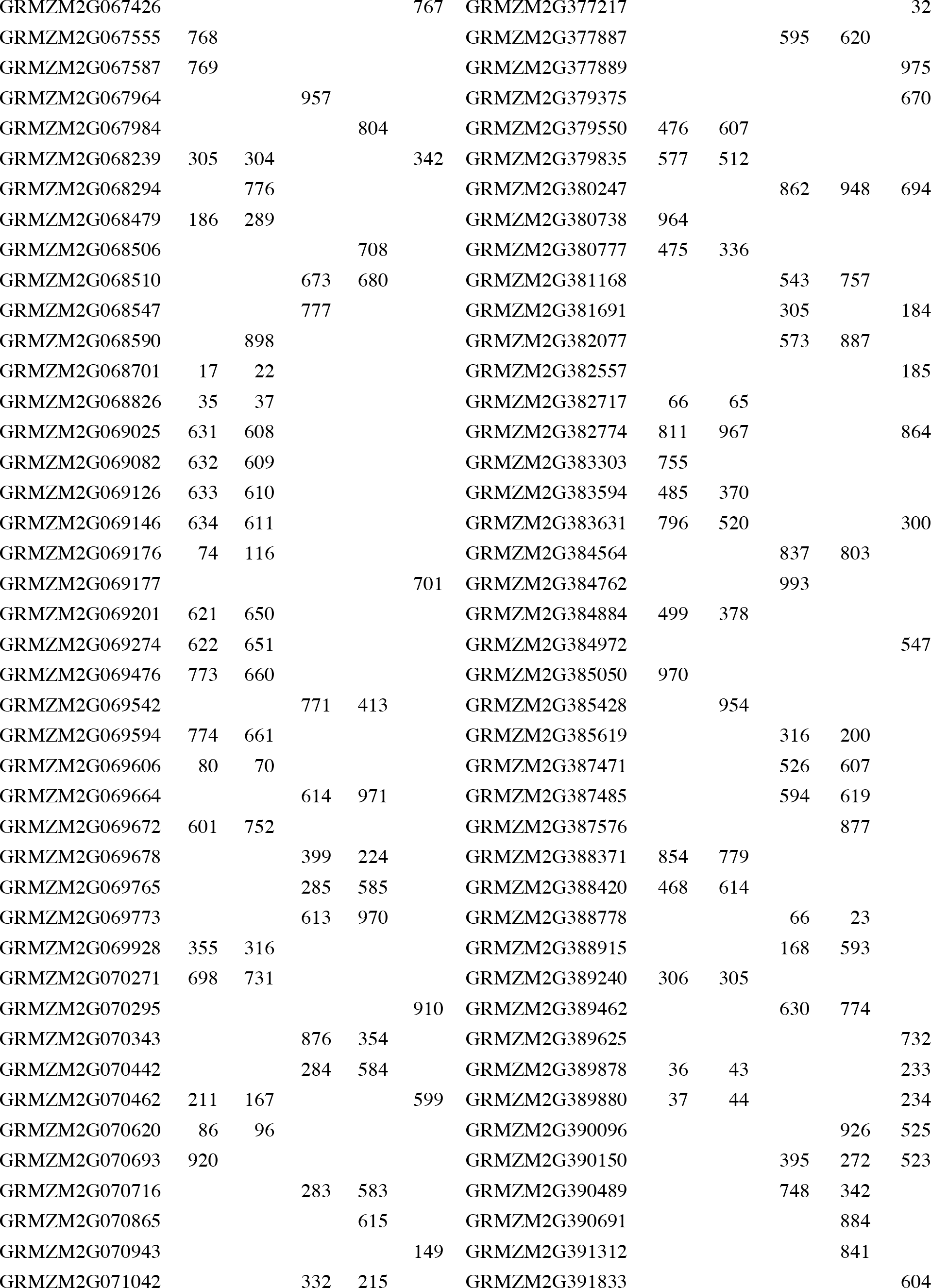
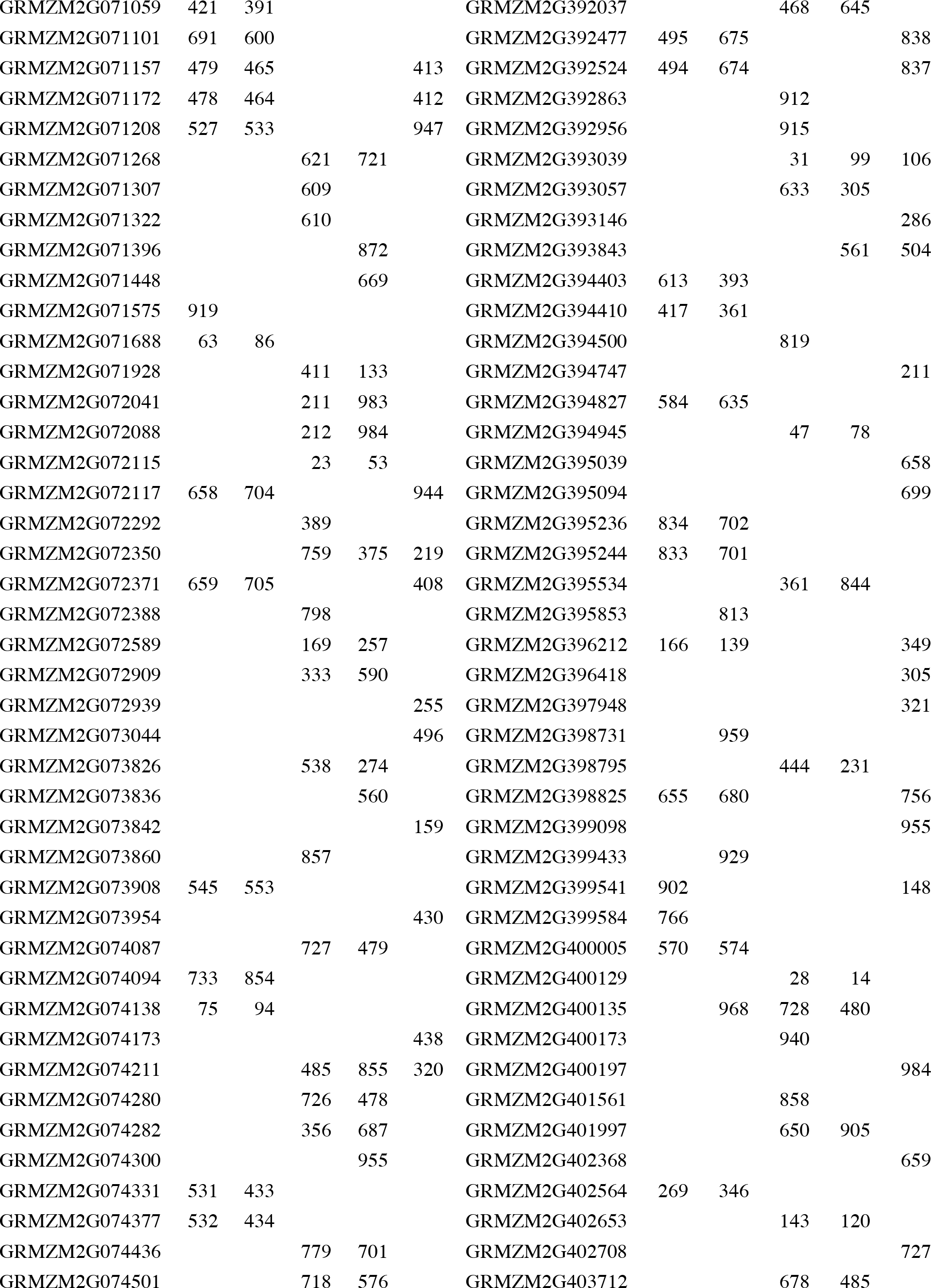
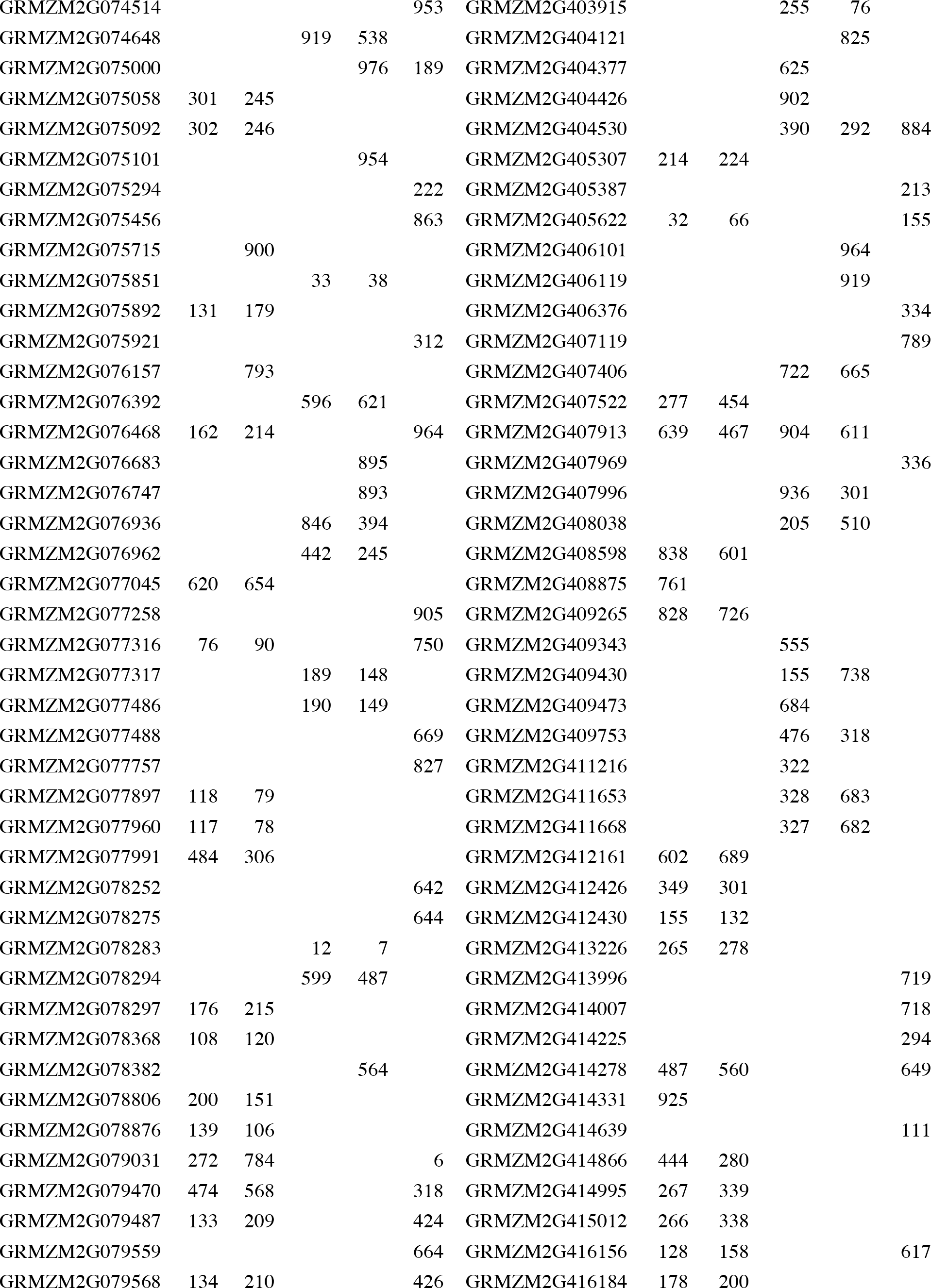
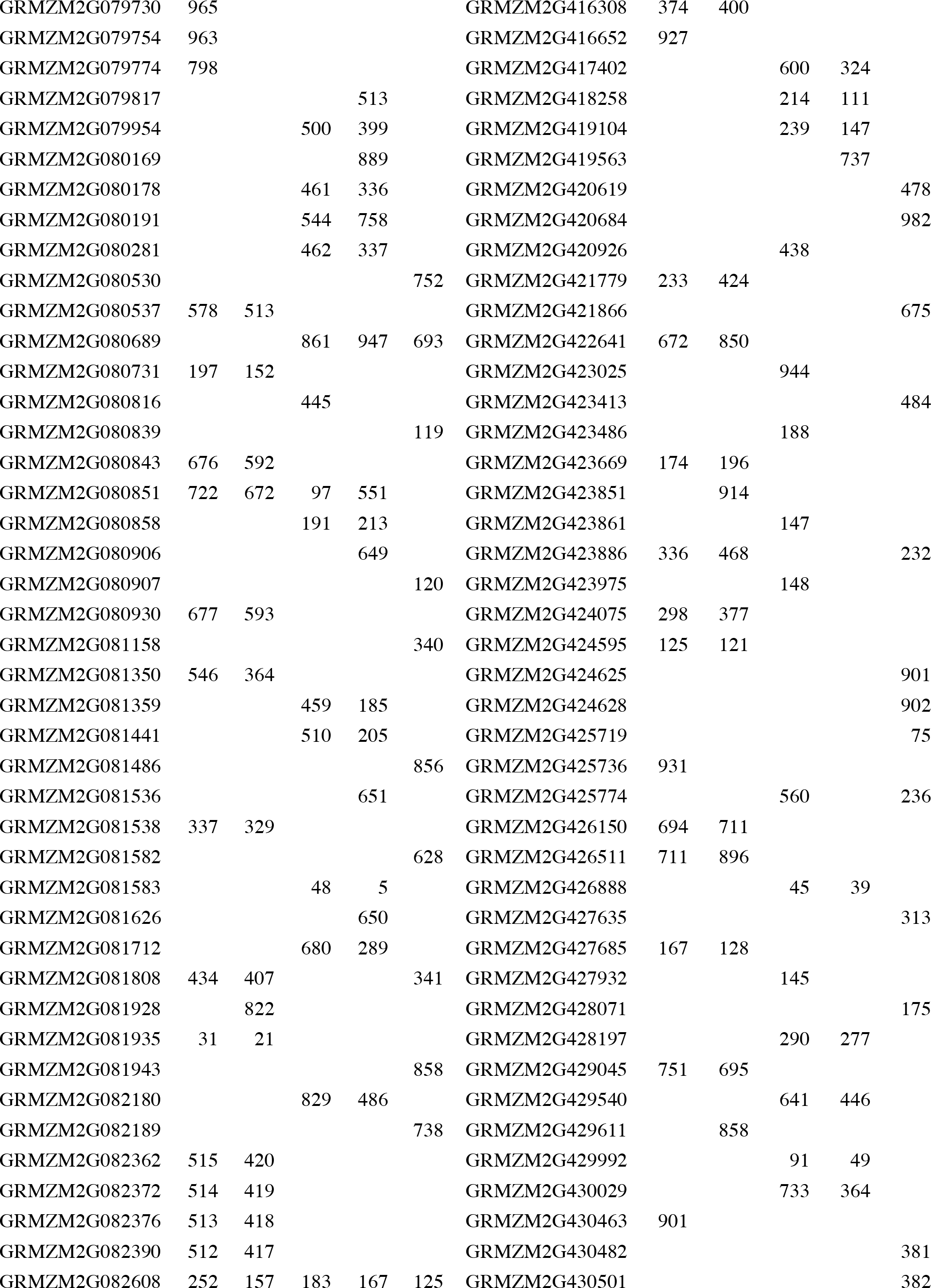
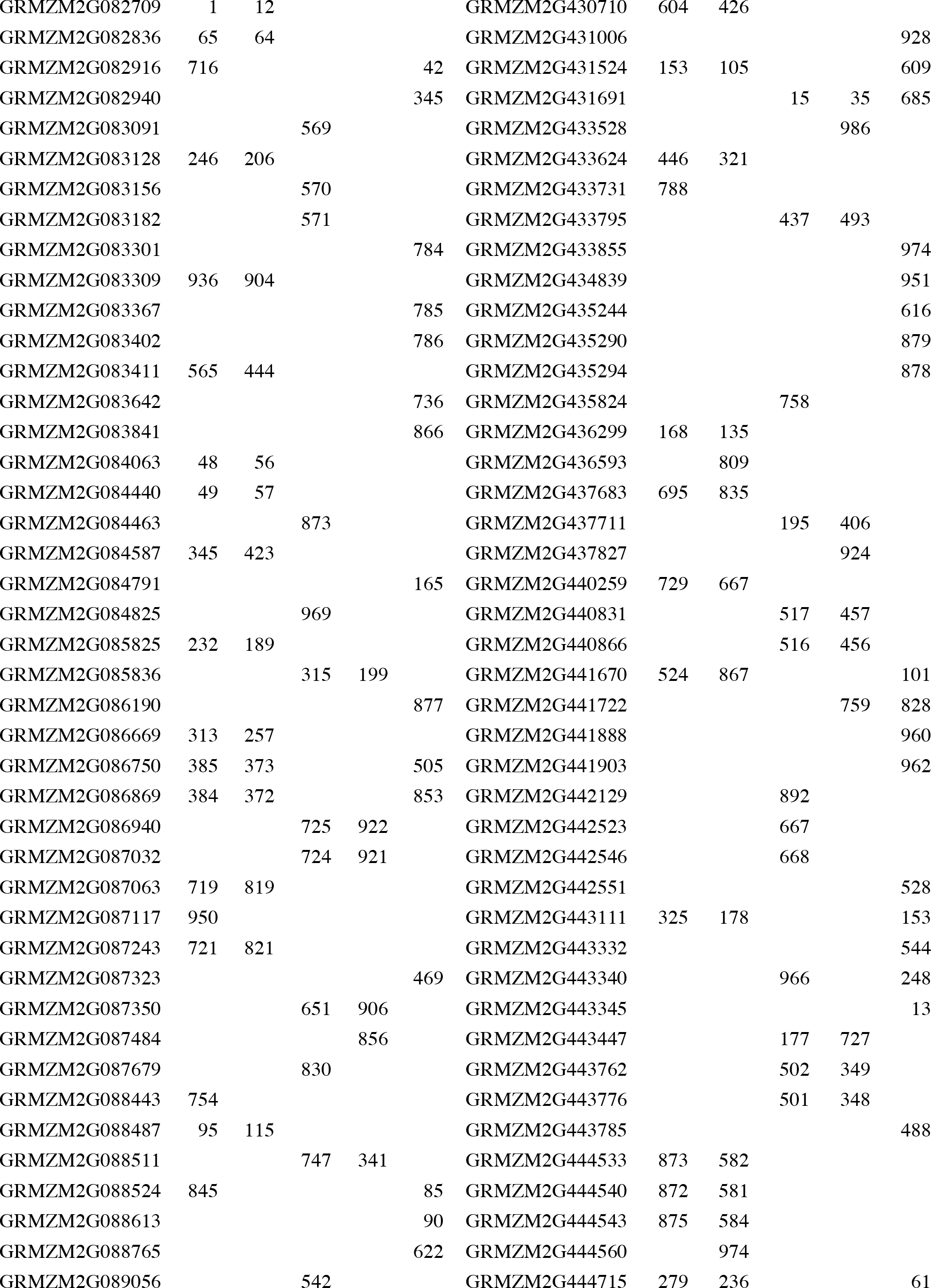
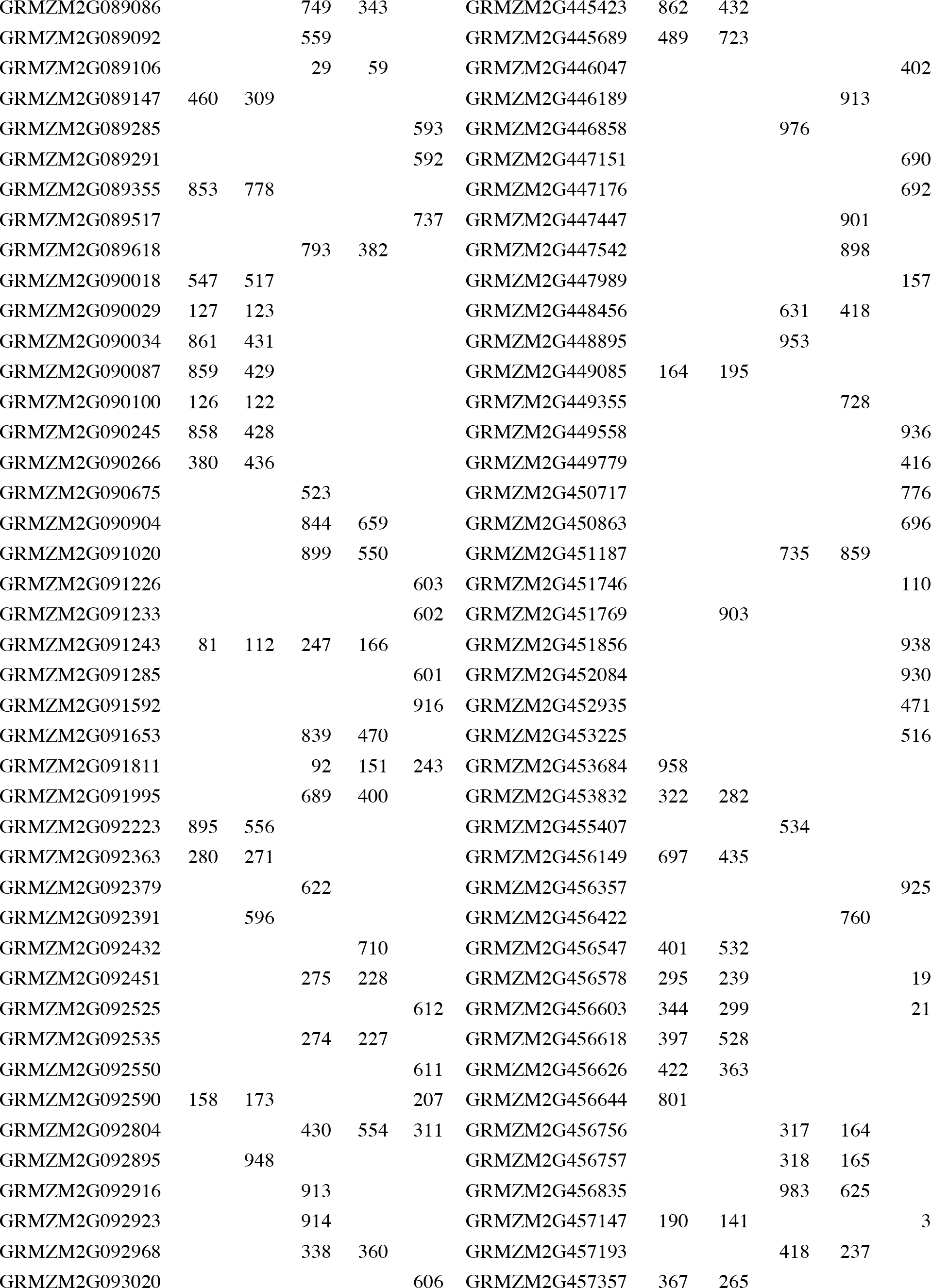
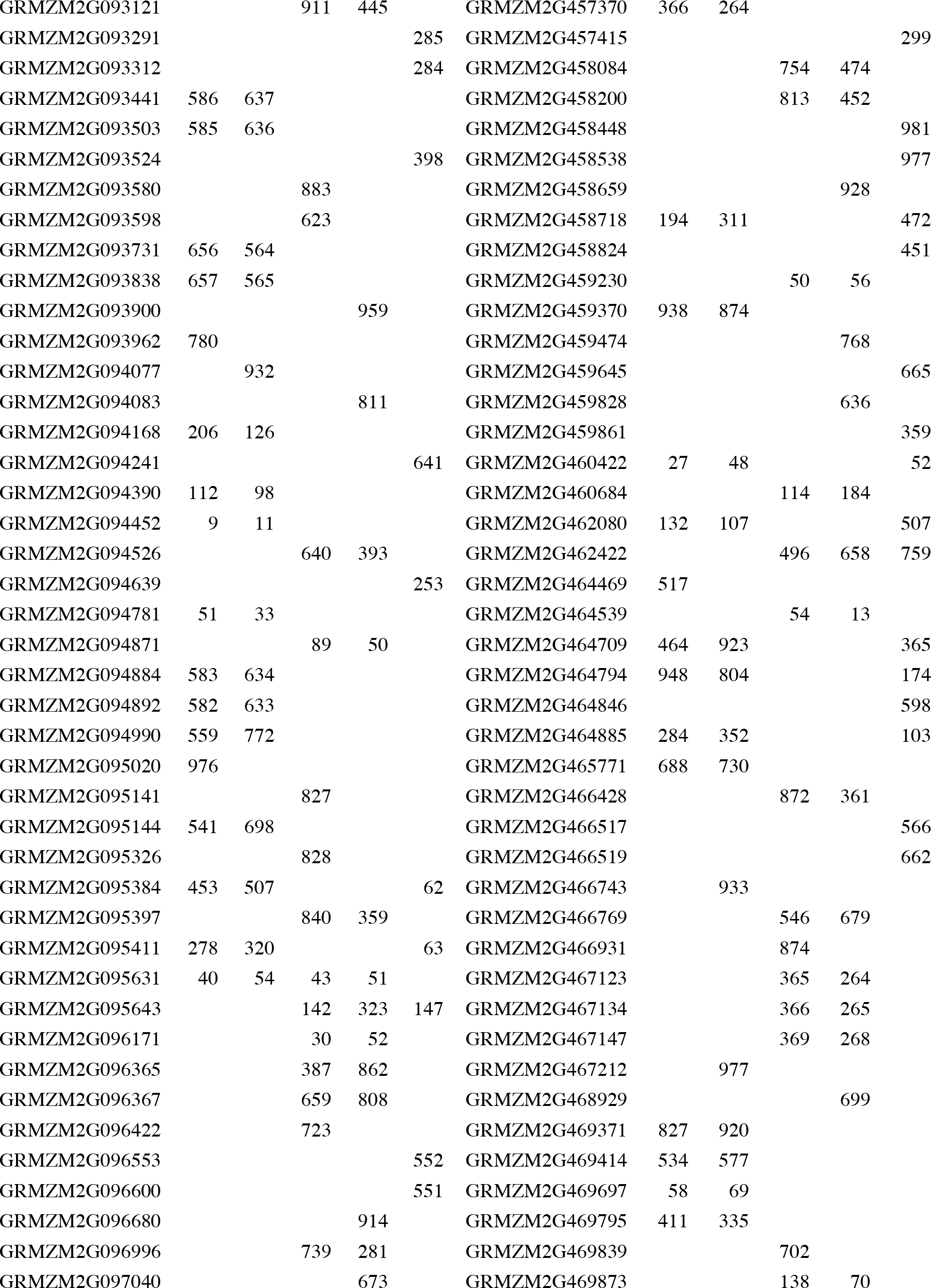
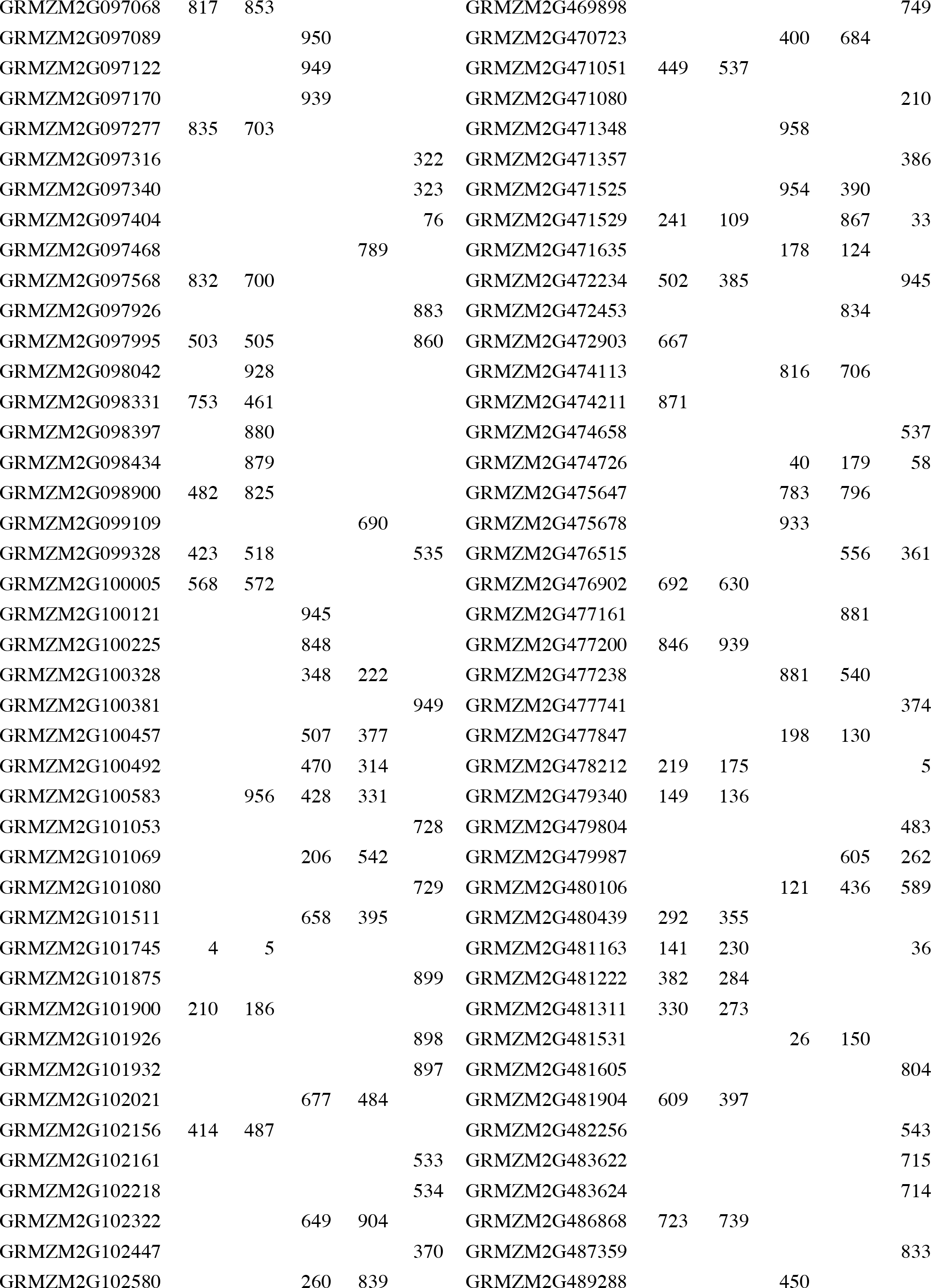
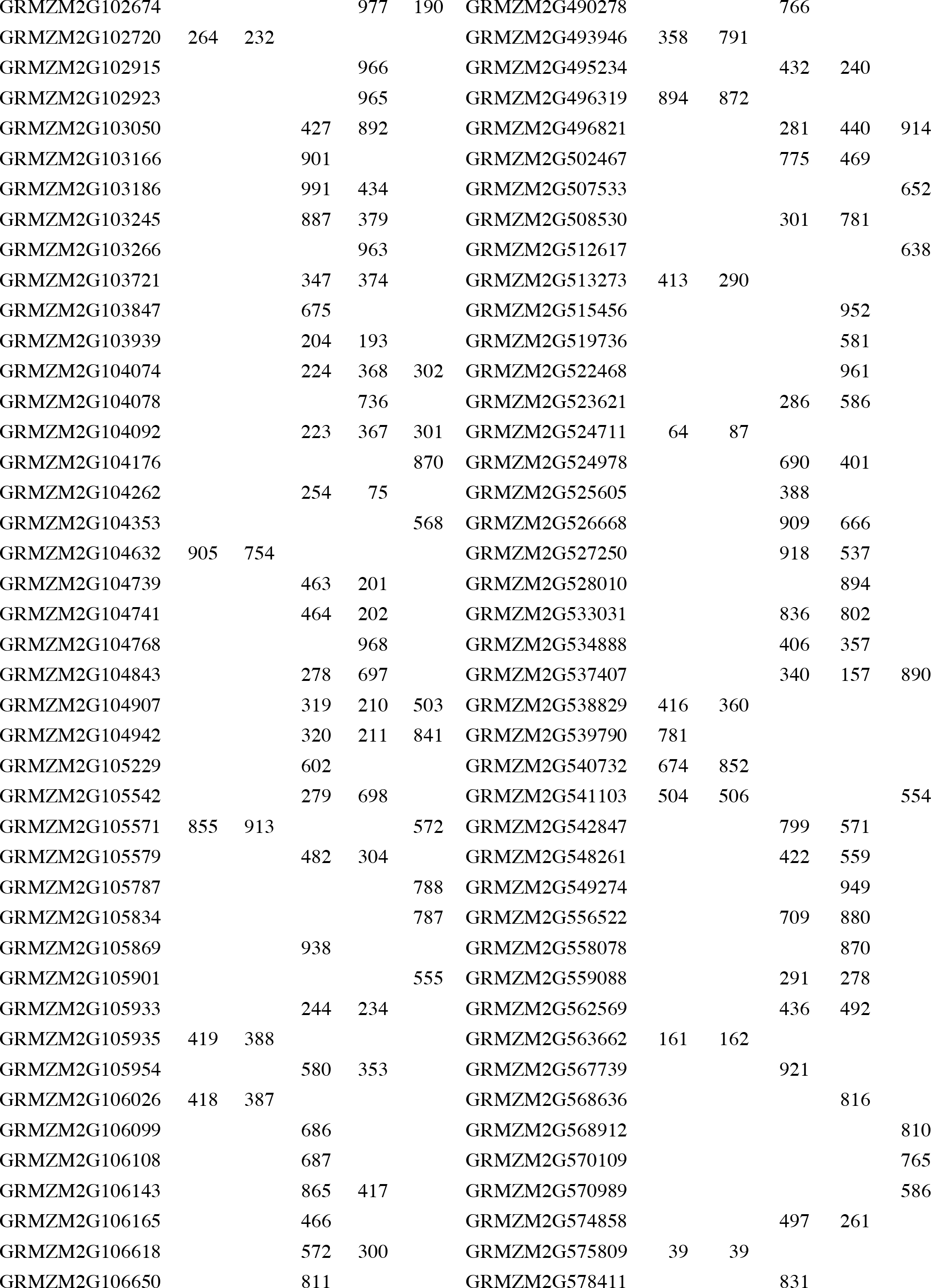
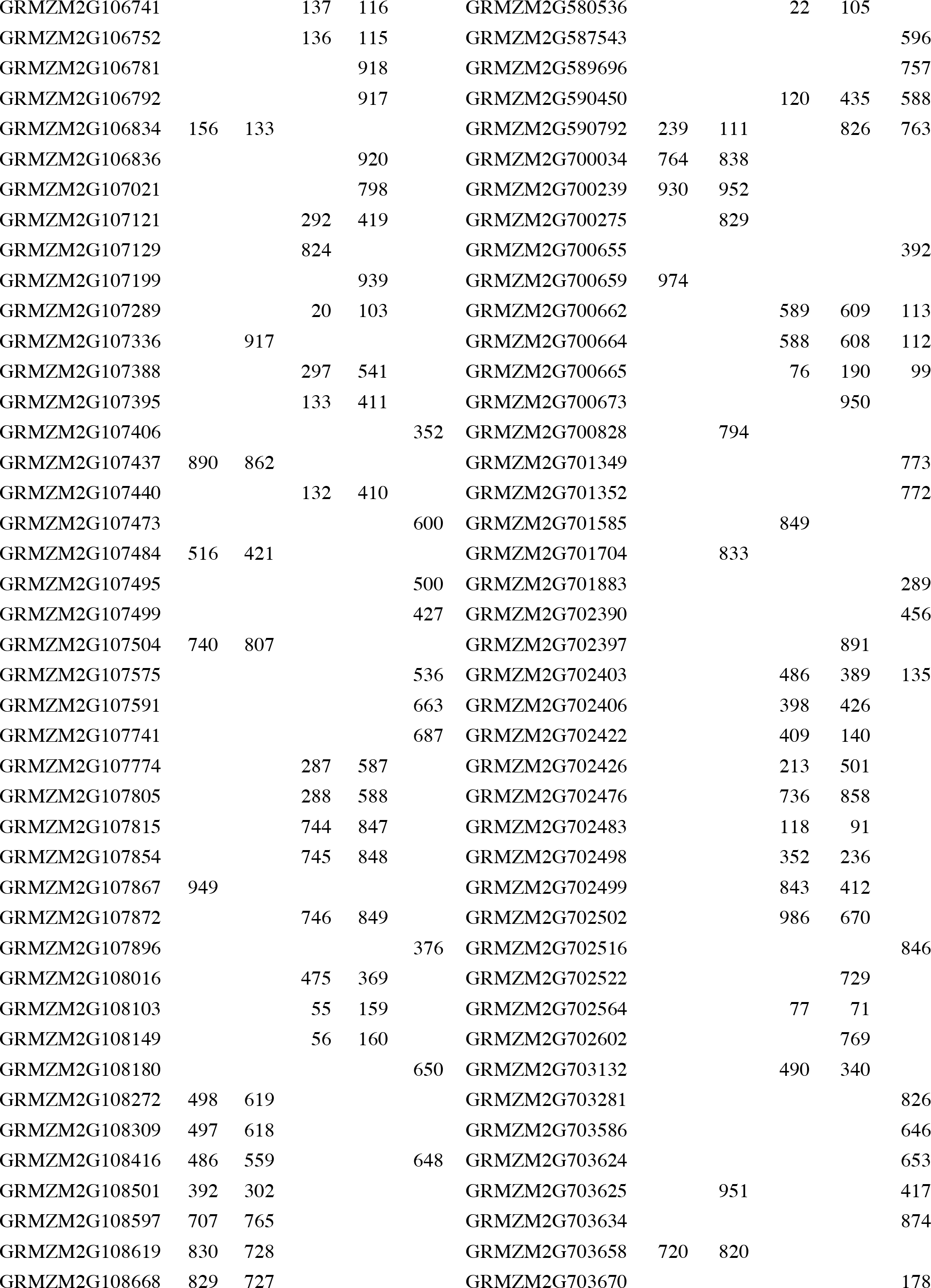
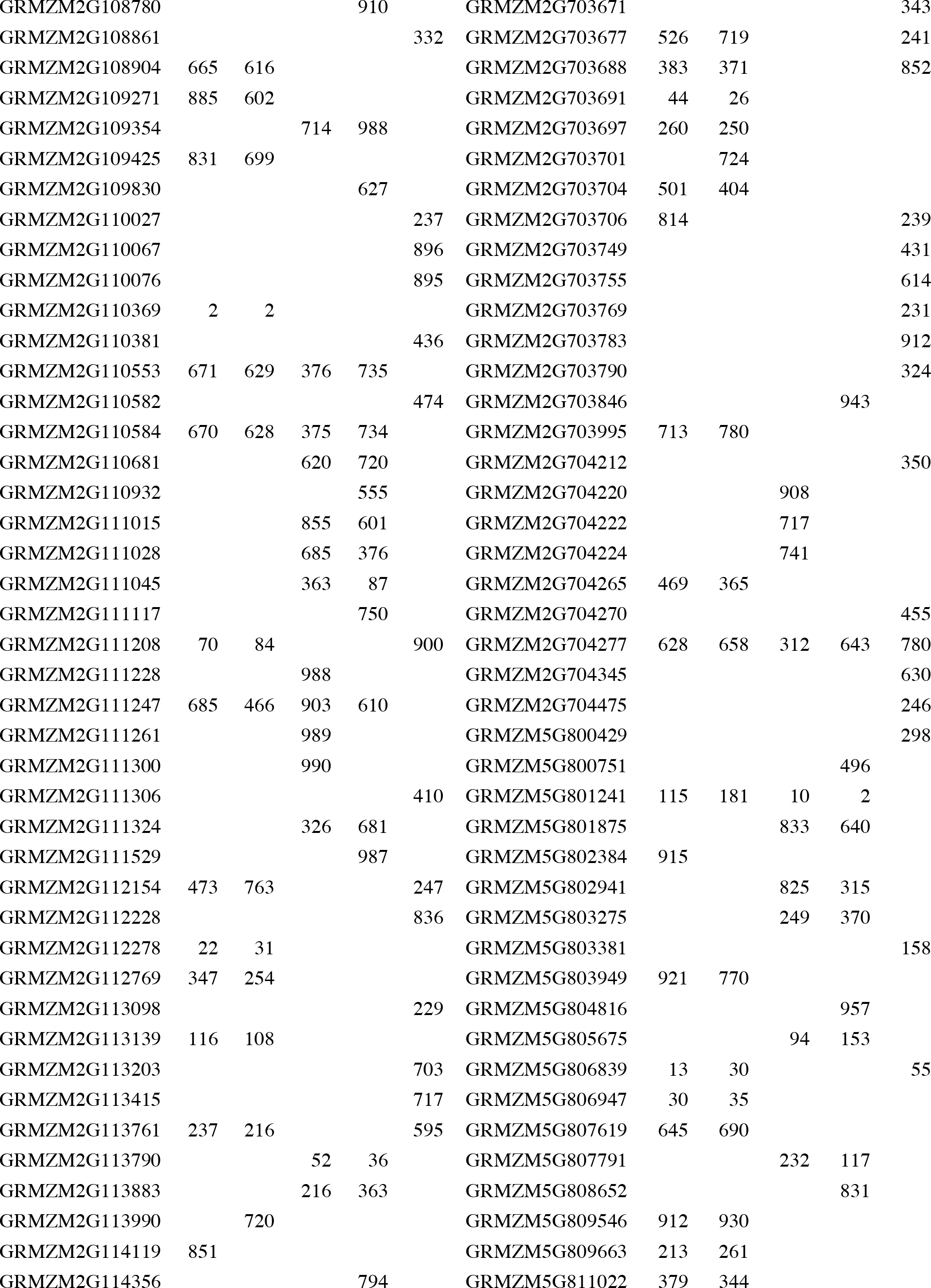
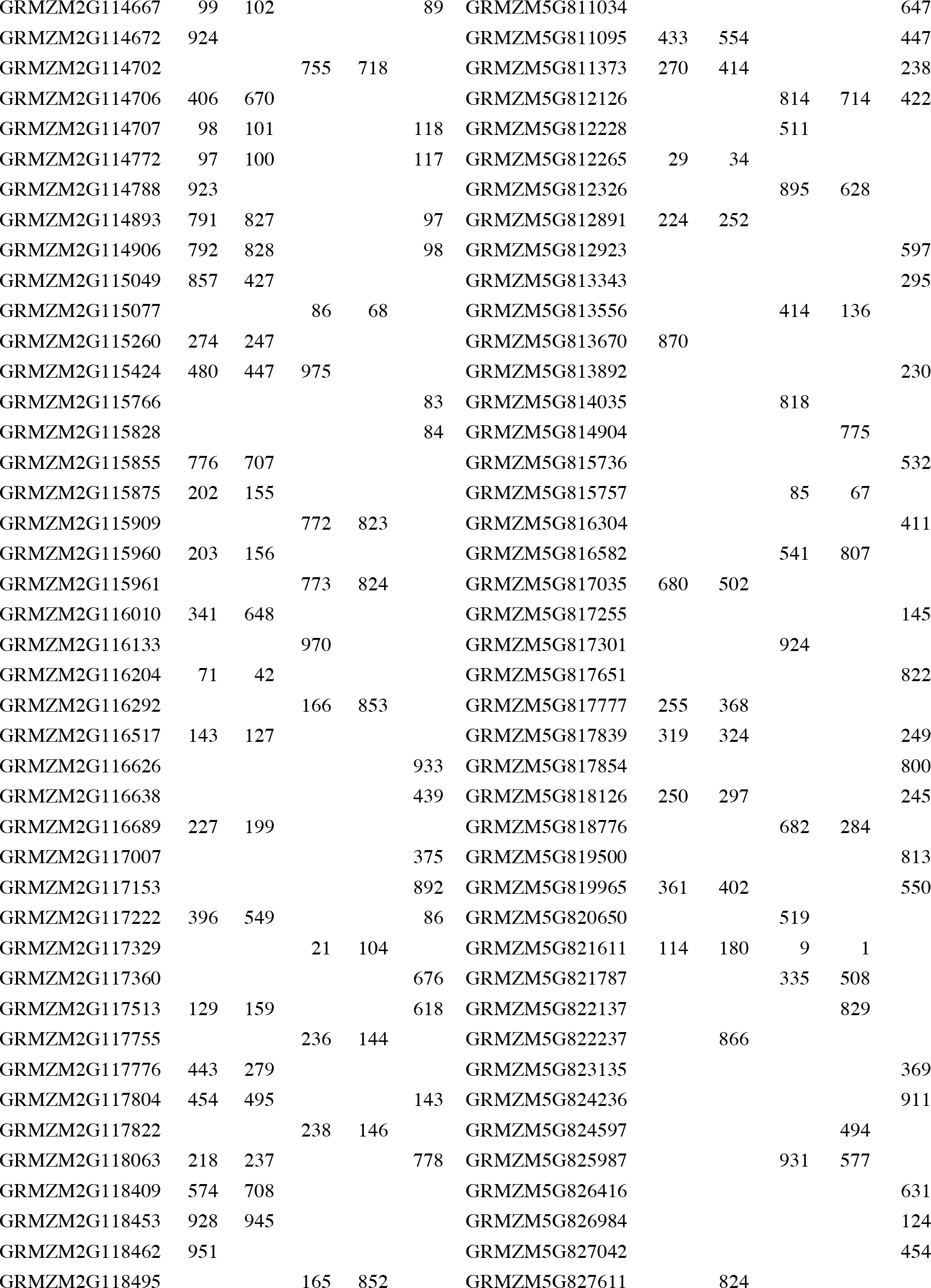
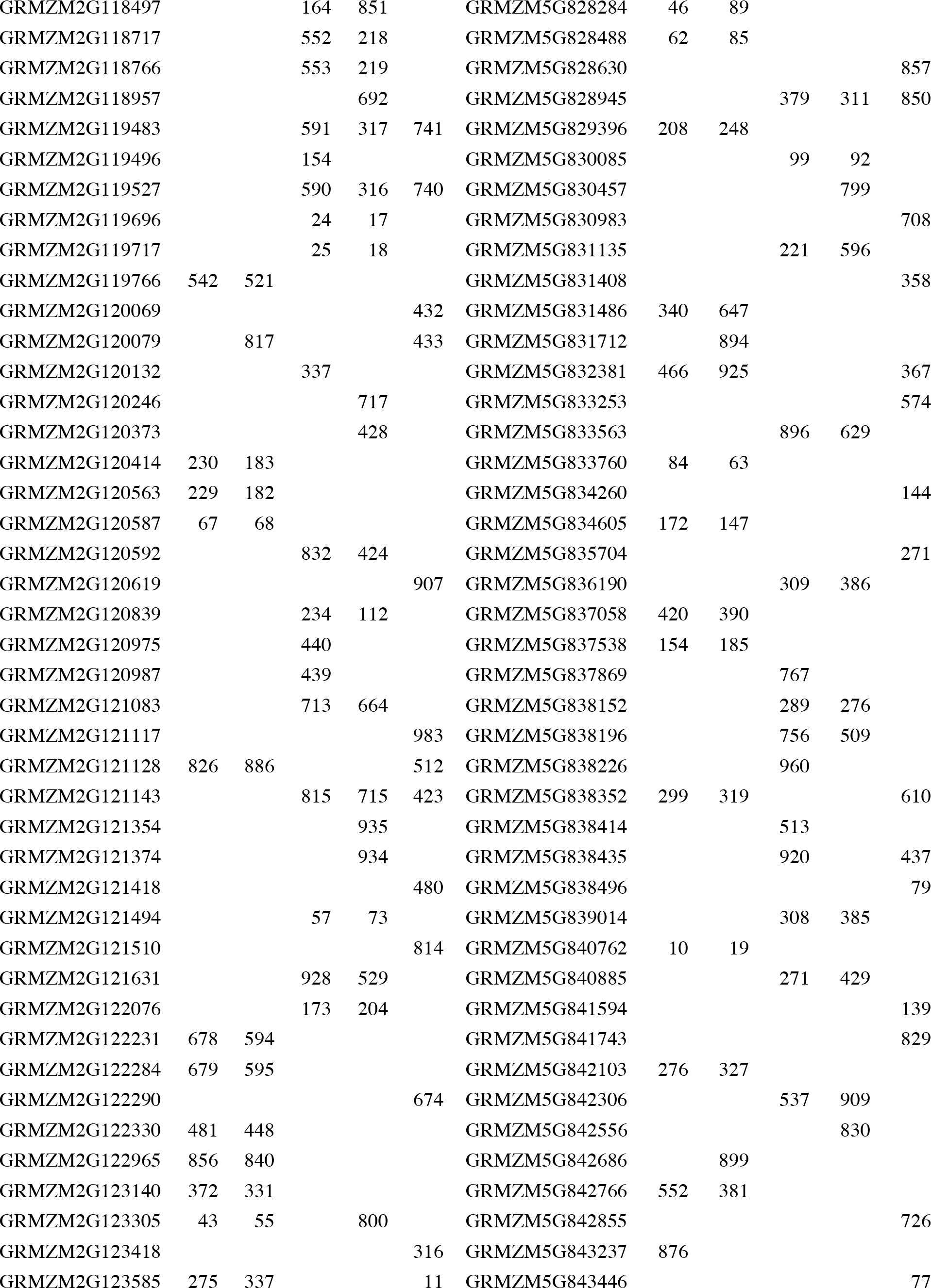
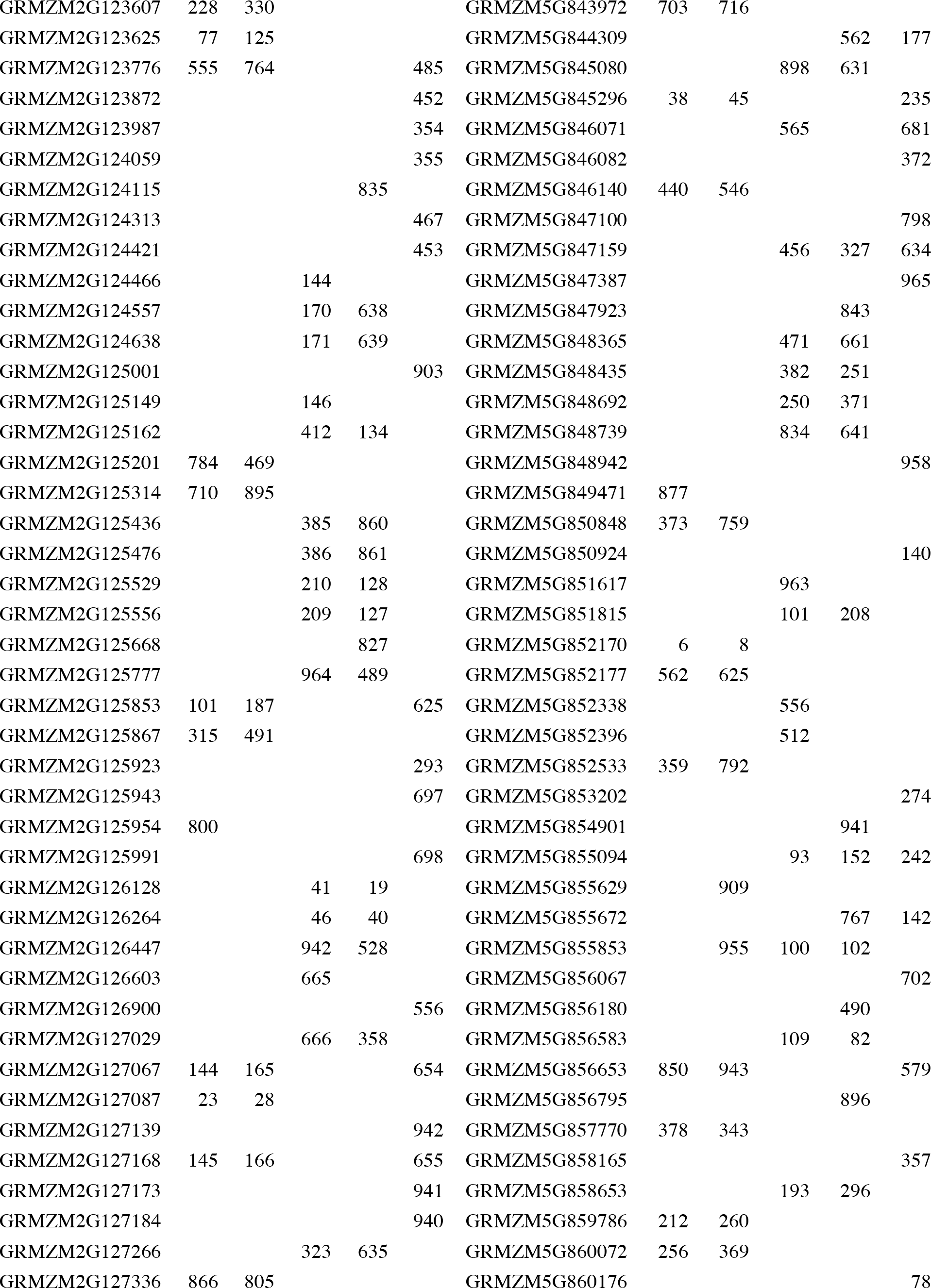
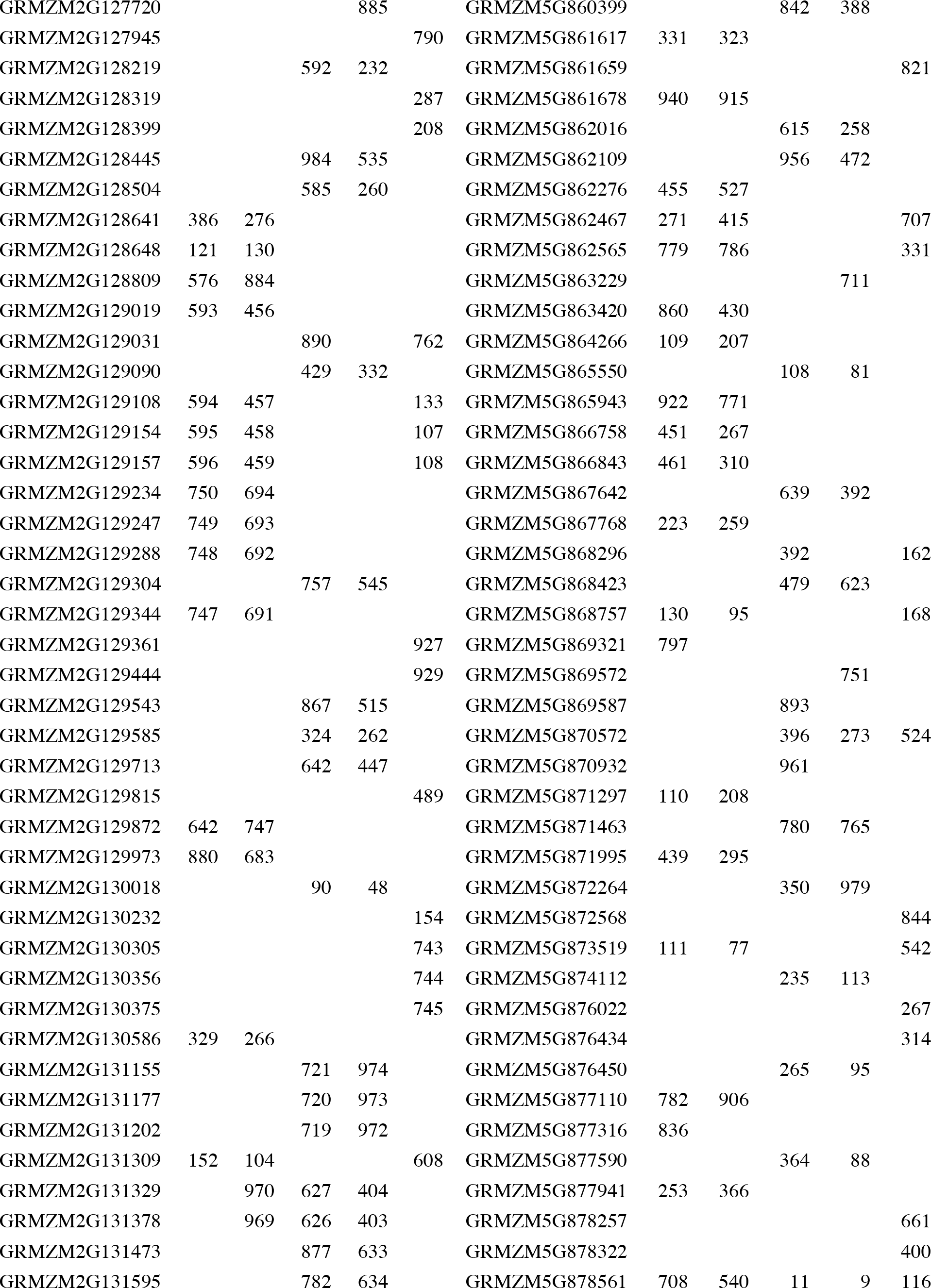
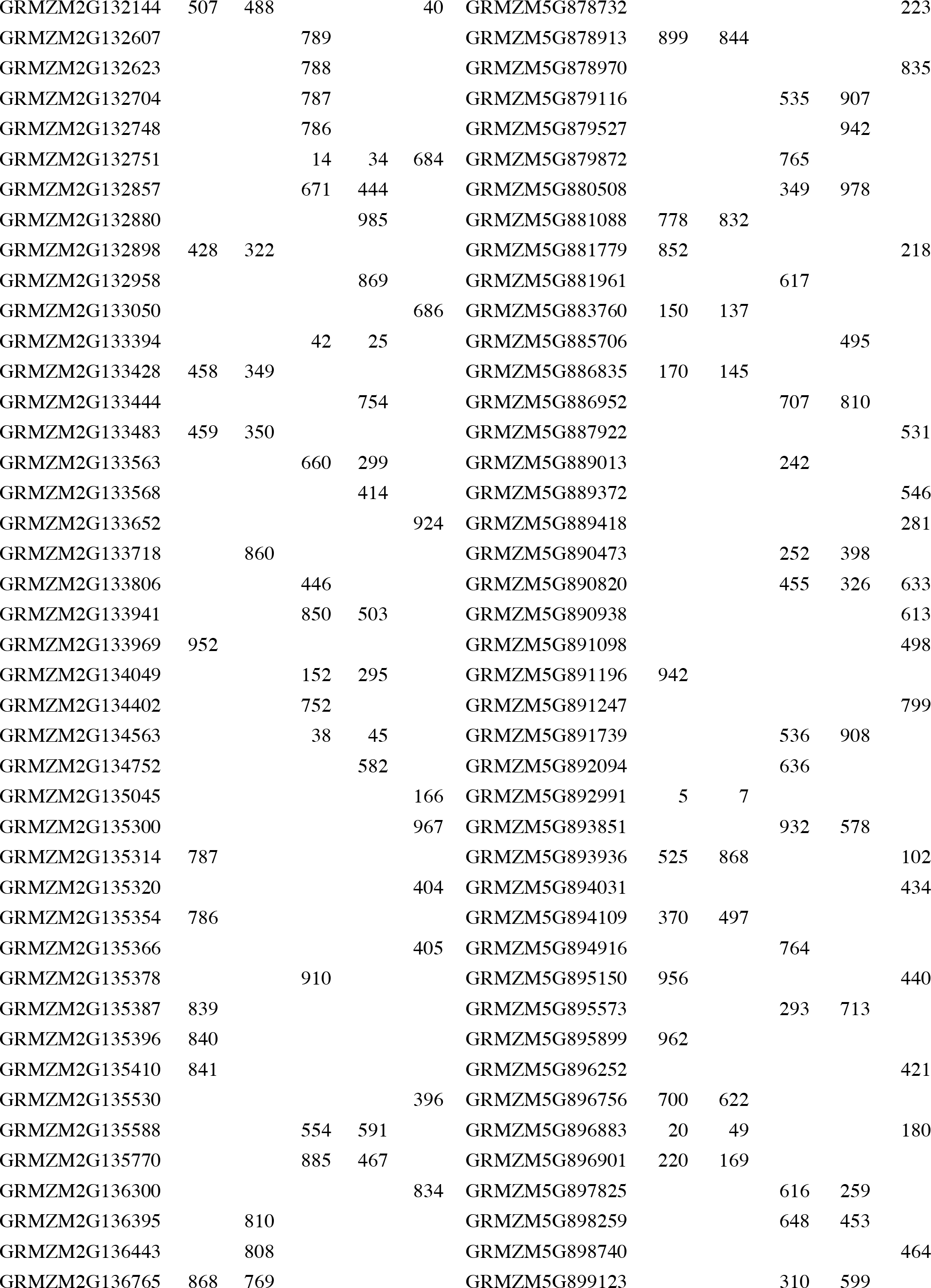
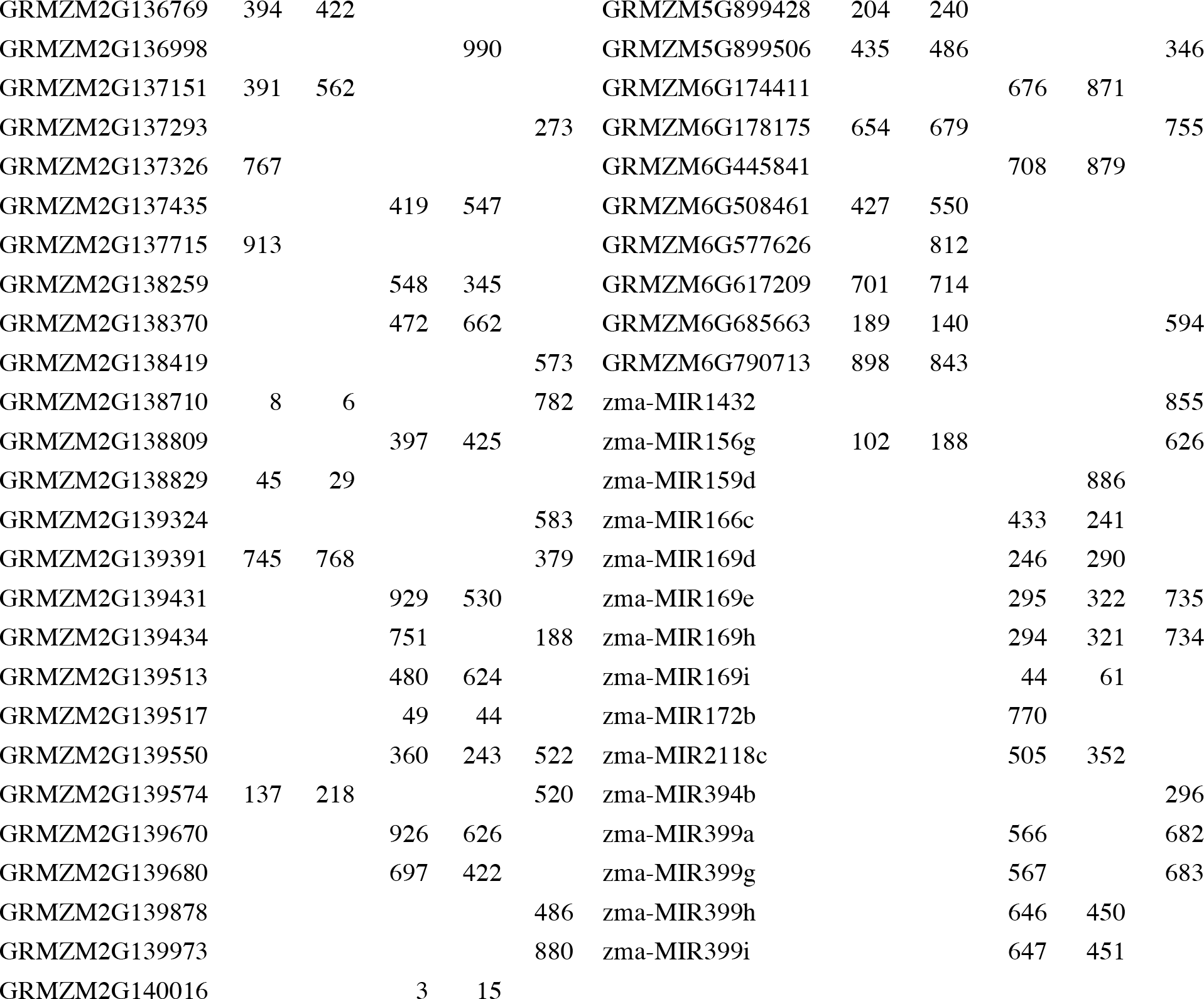
New candidate genes derived from Ames and combined GWAS analysis, based on genes found within 100kbp of the top GWAS hits sorted by additive p-value. The ranking within each subset is given. The first 79 by rank are used in comparison to the shorter Dong et al/ZCN list and the full 918 by rank for each results list are compared with Li et al.

**Table S3.**
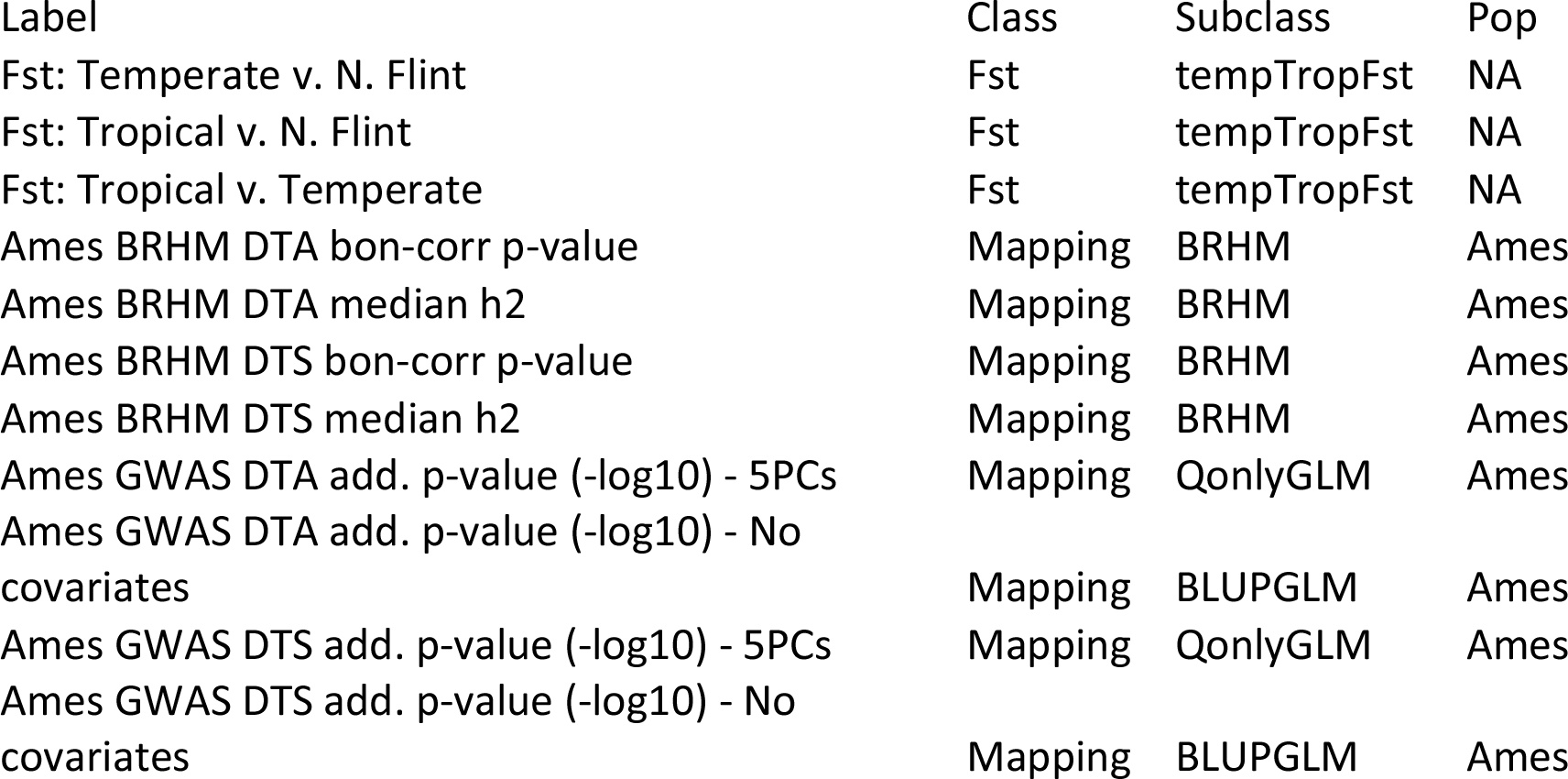
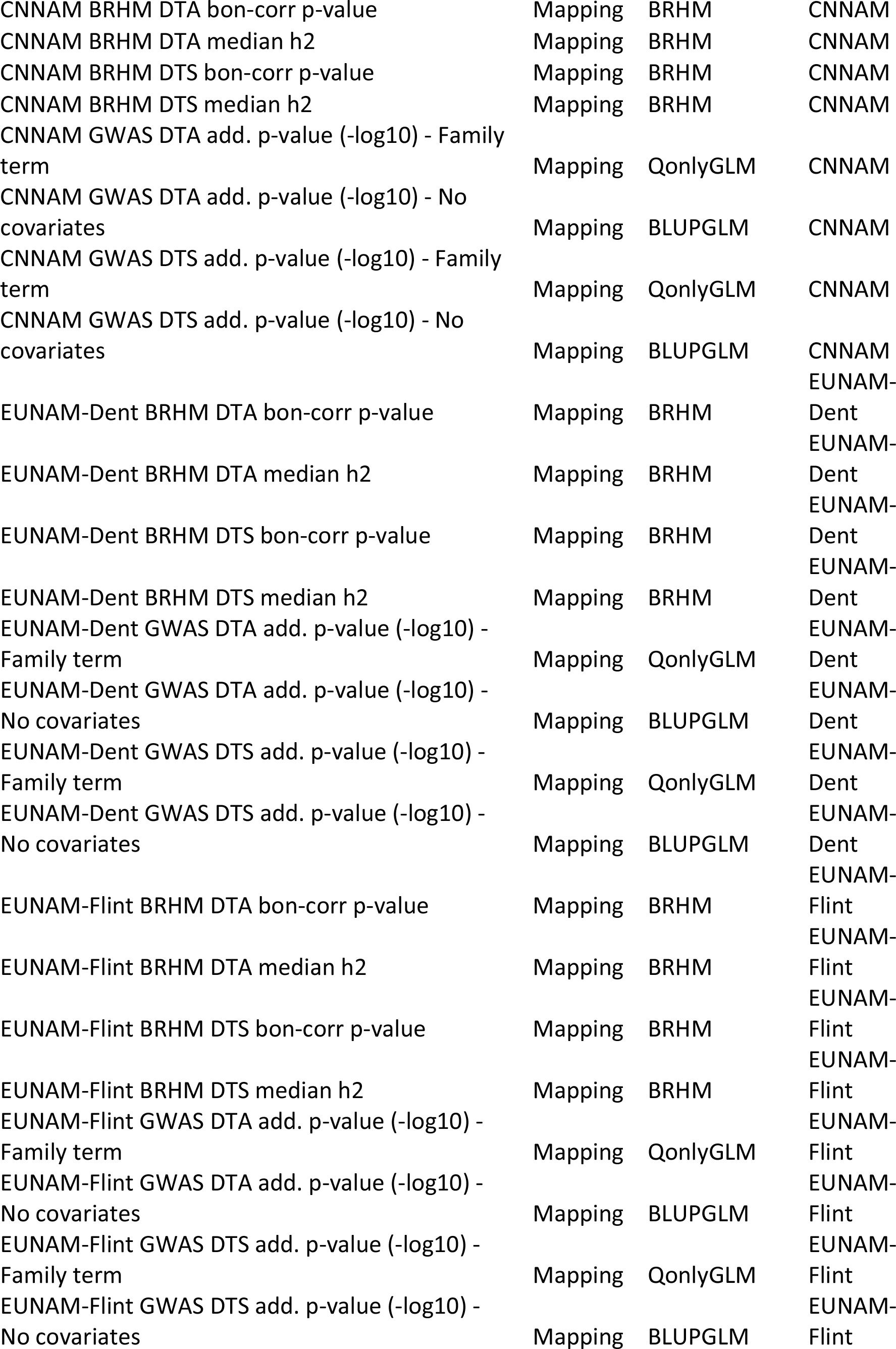
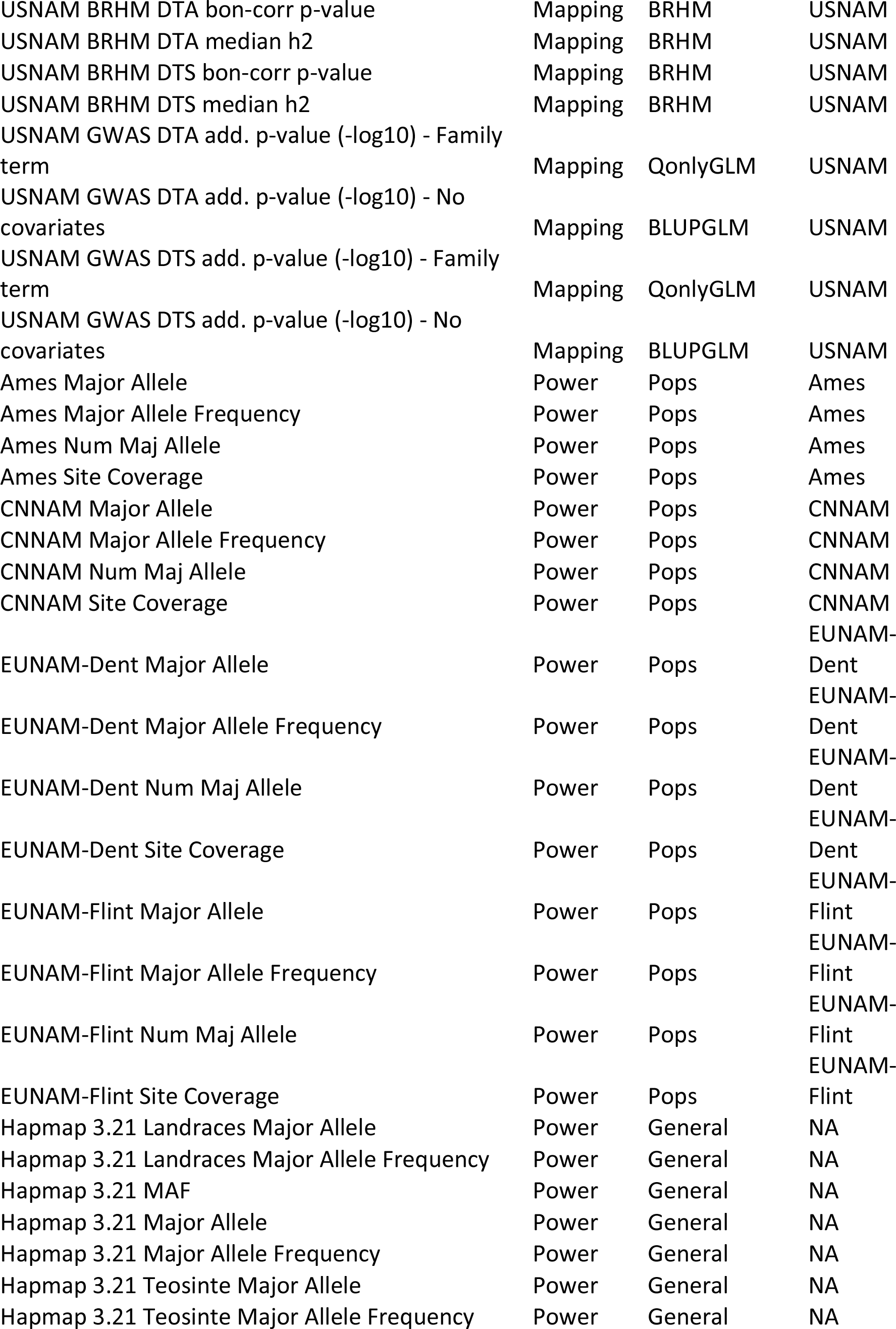
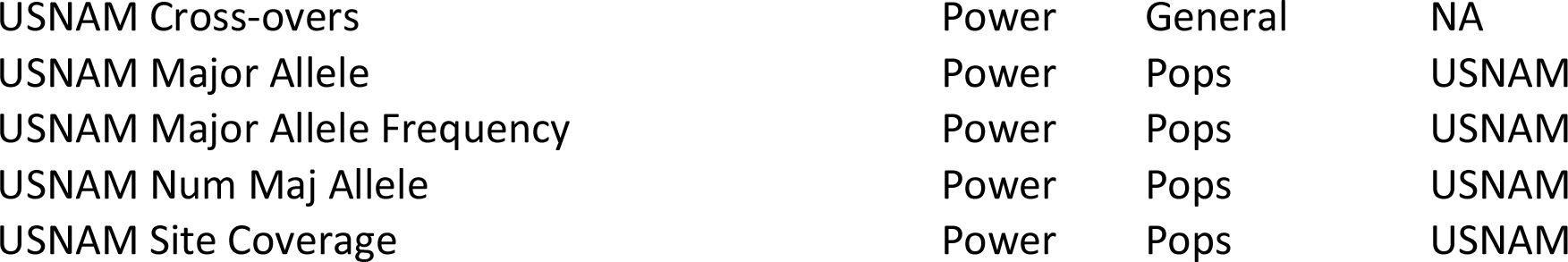
Complete list of non-redundant predictors using in machine learning

| Label | Class | Subclass | Pop |
| --- | --- | --- | --- |
| Fst: Temperate v. N. Flint | Fst | tempTropFst | NA |
| Fst: Tropical v. N. Flint | Fst | tempTropFst | NA |
| Fst: Tropical v. Temperate | Fst | tempTropFst | NA |
| Ames BRHM DTA bon-corr p-value | Mapping | BRHM | Ames |
| Ames BRHM DTA median h2 | Mapping | BRHM | Ames |
| Ames BRHM DTS bon-corr p-value | Mapping | BRHM | Ames |
| Ames BRHM DTS median h2 | Mapping | BRHM | Ames |
| Ames GWAS DTA add. p-value (-log10) - 5PCs | Mapping | QonlyGLM | Ames |
| Ames GWAS DTA add. p-value (-log10) - No covariates | Mapping | BLUPGLM | Ames |
| Ames GWAS DTS add. p-value (-log10) - 5PCs | Mapping | QonlyGLM | Ames |
| Ames GWAS DTS add. p-value (-log10) - No covariates | Mapping | BLUPGLM | Ames |
| CNNAM BRHM DTA bon-corr p-value | Mapping | BRHM | CNNAM |
| CNNAM BRHM DTA median h2 | Mapping | BRHM | CNNAM |
| CNNAM BRHM DTS bon-corr p-value | Mapping | BRHM | CNNAM |
| CNNAM BRHM DTS median h2 | Mapping | BRHM | CNNAM |
| CNNAM GWAS DTA add. p-value (-log10) - Family term | Mapping | QonlyGLM | CNNAM |
| CNNAM GWAS DTA add. p-value (-log10) - No covariates | Mapping | BLUPGLM | CNNAM |
| CNNAM GWAS DTS add. p-value (-log10) - Family term | Mapping | QonlyGLM | CNNAM |
| CNNAM GWAS DTS add. p-value (-log10) - No covariates | Mapping | BLUPGLM | CNNAM |
| EUNAM-Dent BRHM DTA bon-corr p-value | Mapping | BRHM | EUNAM-Dent |
| EUNAM-Dent BRHM DTA median h2 | Mapping | BRHM | EUNAM-Dent |
| EUNAM-Dent BRHM DTS bon-corr p-value | Mapping | BRHM | EUNAM-Dent |
| EUNAM-Dent BRHM DTS median h2 | Mapping | BRHM | EUNAM-Dent |
| EUNAM-Dent GWAS DTA add. p-value (-log10) - Family term | Mapping | QonlyGLM | EUNAM-Dent |
| EUNAM-Dent GWAS DTA add. p-value (-log10) - No covariates | Mapping | BLUPGLM | EUNAM-Dent |
| EUNAM-Dent GWAS DTS add. p-value (-log10) - Family term | Mapping | QonlyGLM | EUNAM-Dent |
| EUNAM-Dent GWAS DTS add. p-value (-log10) - No covariates | Mapping | BLUPGLM | EUNAM-Dent |
| EUNAM-Flint BRHM DTA bon-corr p-value | Mapping | BRHM | EUNAM-Flint |
| EUNAM-Flint BRHM DTA median h2 | Mapping | BRHM | EUNAM-Flint |
| EUNAM-Flint BRHM DTS bon-corr p-value | Mapping | BRHM | EUNAM-Flint |
| EUNAM-Flint BRHM DTS median h2 | Mapping | BRHM | EUNAM-Flint |
| EUNAM-Flint GWAS DTA add. p-value (-log10) - Family term | Mapping | QonlyGLM | EUNAM-Flint |
| EUNAM-Flint GWAS DTA add. p-value (-log10) - No covariates | Mapping | BLUPGLM | EUNAM-Flint |
| EUNAM-Flint GWAS DTS add. p-value (-log10) - Family term | Mapping | QonlyGLM | EUNAM-Flint |
| EUNAM-Flint GWAS DTS add. p-value (-log10) - No covariates | Mapping | BLUPGLM | EUNAM-Flint |
| USNAM BRHM DTA bon-corr p-value | Mapping | BRHM | USNAM |
| USNAM BRHM DTA median h2 | Mapping | BRHM | USNAM |
| USNAM BRHM DTS bon-corr p-value | Mapping | BRHM | USNAM |
| USNAM BRHM DTS median h2 | Mapping | BRHM | USNAM |
| USNAM GWAS DTA add. p-value (-log10) - Family term | Mapping | QonlyGLM | USNAM |
| USNAM GWAS DTA add. p-value (-log10) - No covariates | Mapping | BLUPGLM | USNAM |
| USNAM GWAS DTS add. p-value (-log10) - Family term | Mapping | QonlyGLM | USNAM |
| USNAM GWAS DTS add. p-value (-log10) - No covariates | Mapping | BLUPGLM | USNAM |
| Ames Major Allele | Power | Pops | Ames |
| Ames Major Allele Frequency | Power | Pops | Ames |
| Ames Num Maj Allele | Power | Pops | Ames |
| Ames Site Coverage | Power | Pops | Ames |
| CNNAM Major Allele | Power | Pops | CNNAM |
| CNNAM Major Allele Frequency | Power | Pops | CNNAM |
| CNNAM Num Maj Allele | Power | Pops | CNNAM |
| CNNAM Site Coverage | Power | Pops | CNNAM |
| EUNAM-Dent Major Allele | Power | Pops | EUNAM-Dent |
| EUNAM-Dent Major Allele Frequency | Power | Pops | EUNAM-Dent |
| EUNAM-Dent Num Maj Allele | Power | Pops | EUNAM-Dent |
| EUNAM-Dent Site Coverage | Power | Pops | EUNAM-Dent |
| EUNAM-Flint Major Allele | Power | Pops | EUNAM-Flint |
| EUNAM-Flint Major Allele Frequency | Power | Pops | EUNAM-Flint |
| EUNAM-Flint Num Maj Allele | Power | Pops | EUNAM-Flint |
| EUNAM-Flint Site Coverage | Power | Pops | EUNAM-Flint |
| Hapmap 3.21 Landraces Major Allele | Power | General | NA |
| Hapmap 3.21 Landraces Major Allele Frequency | Power | General | NA |
| Hapmap 3.21 MAF | Power | General | NA |
| Hapmap 3.21 Major Allele | Power | General | NA |
| Hapmap 3.21 Major Allele Frequency | Power | General | NA |
| Hapmap 3.21 Teosinte Major Allele | Power | General | NA |
| Hapmap 3.21 Teosinte Major Allele Frequency | Power | General | NA |
| USNAM Cross-overs | Power | General | NA |
| USNAM Major Allele | Power | Pops | USNAM |
| USNAM Major Allele Frequency | Power | Pops | USNAM |
| USNAM Num Maj Allele | Power | Pops | USNAM |
| USNAM Site Coverage | Power | Pops | USNAM |

**Table S4.**
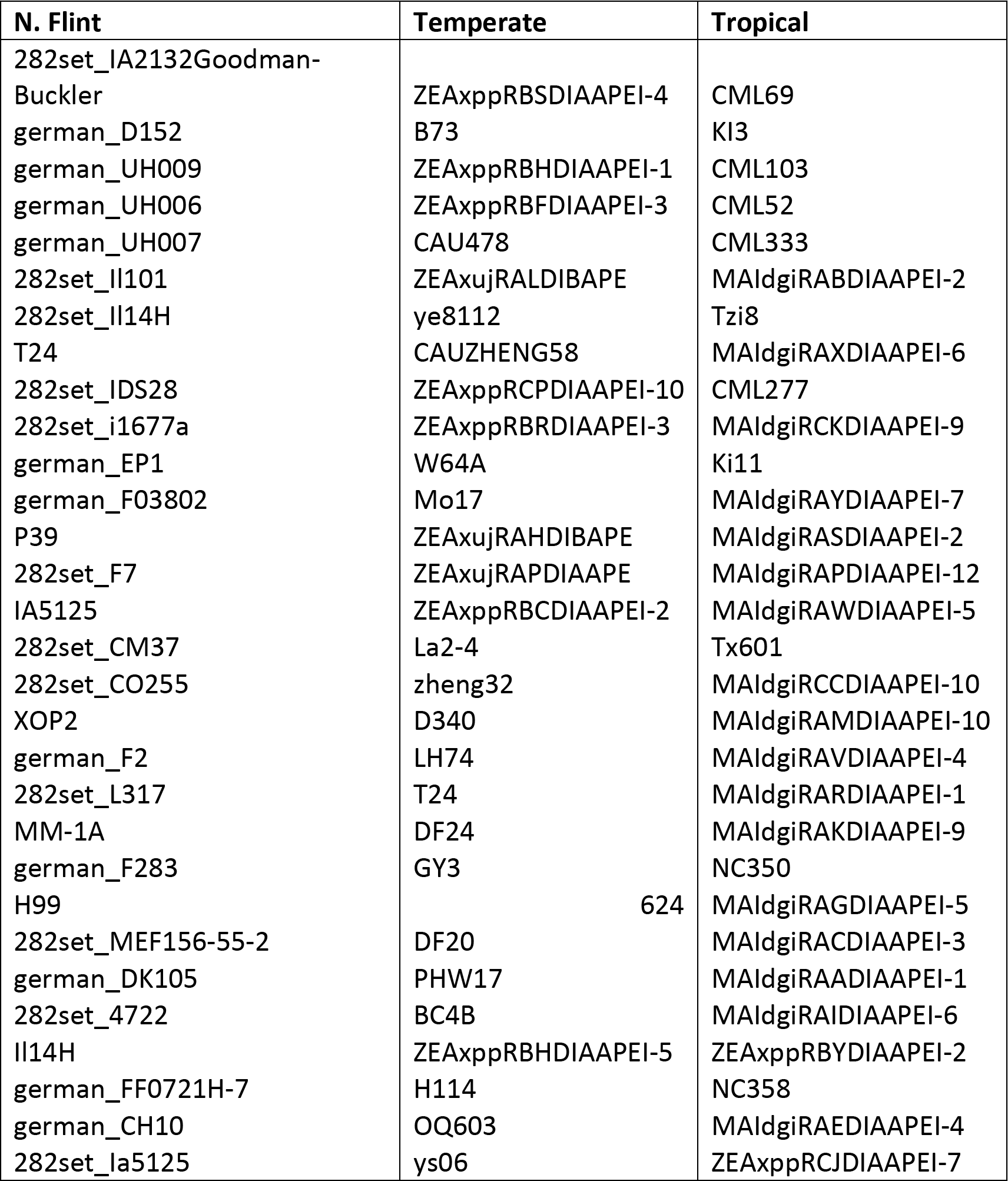
Hapmap 3.21 taxa included in the North American Fst subsets. Names are not necessarily representative of the country of origin (eg. german_X). For list of EUNAM parent origins see Bauer et al (*88*).

**Table S5.**
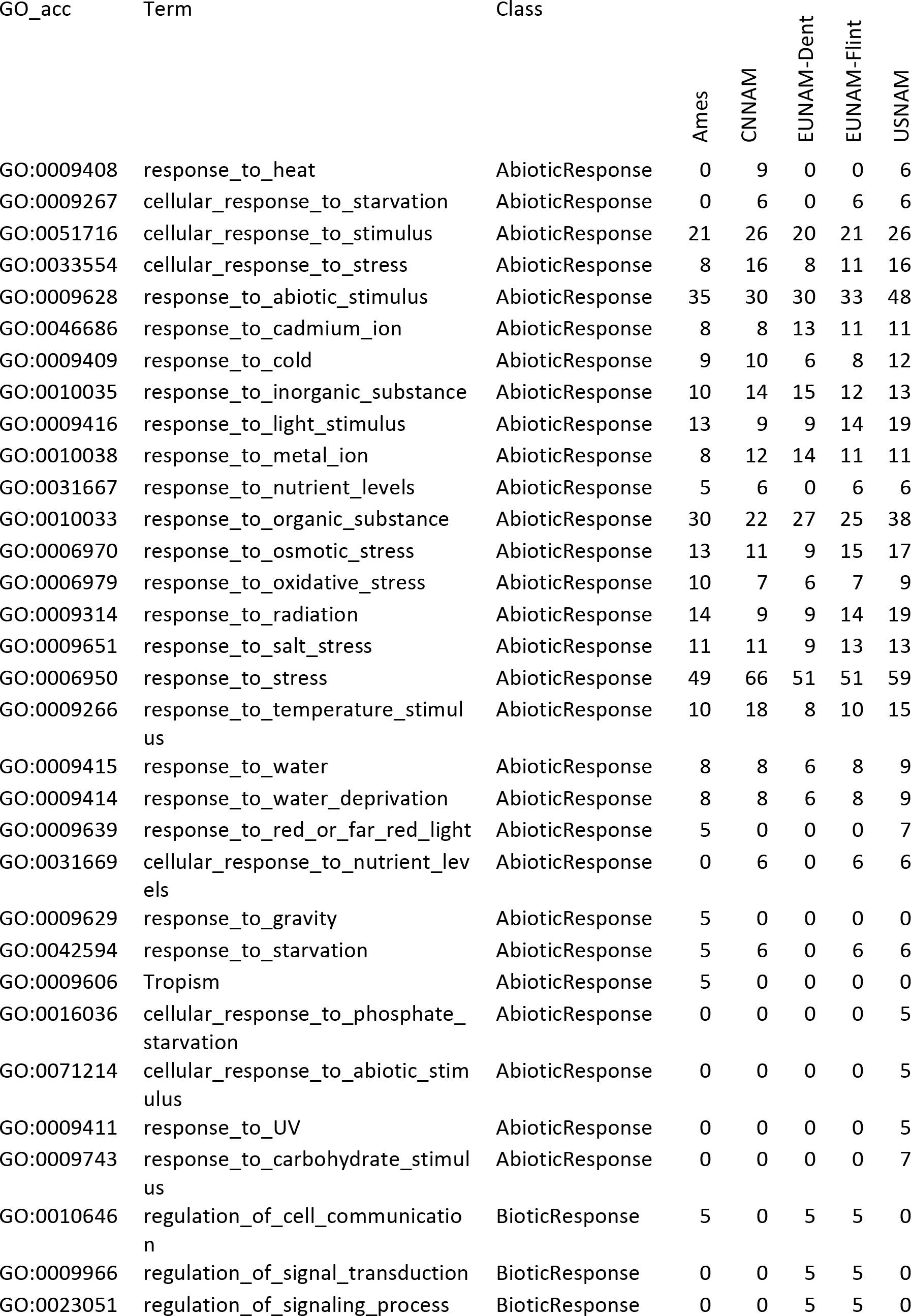
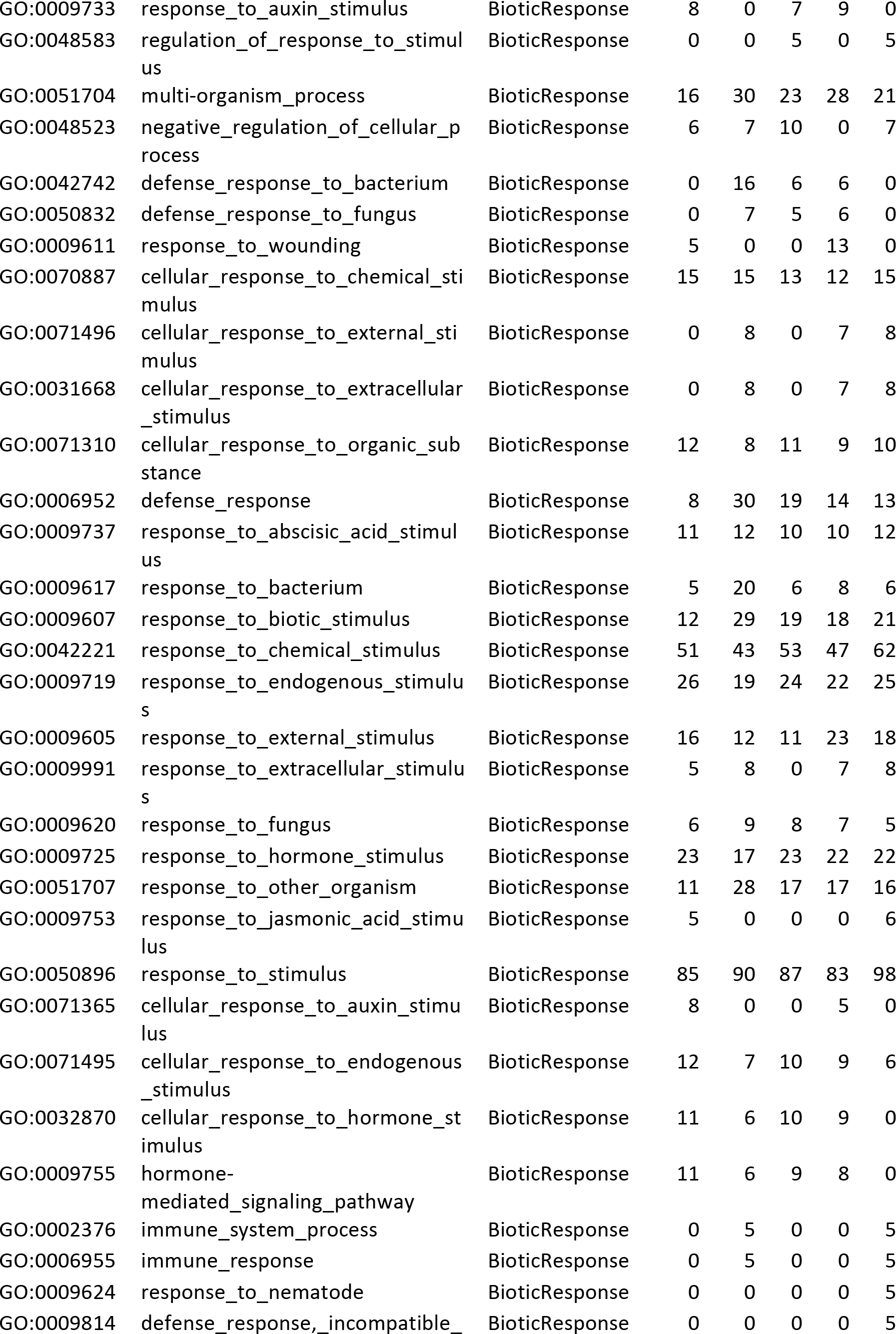
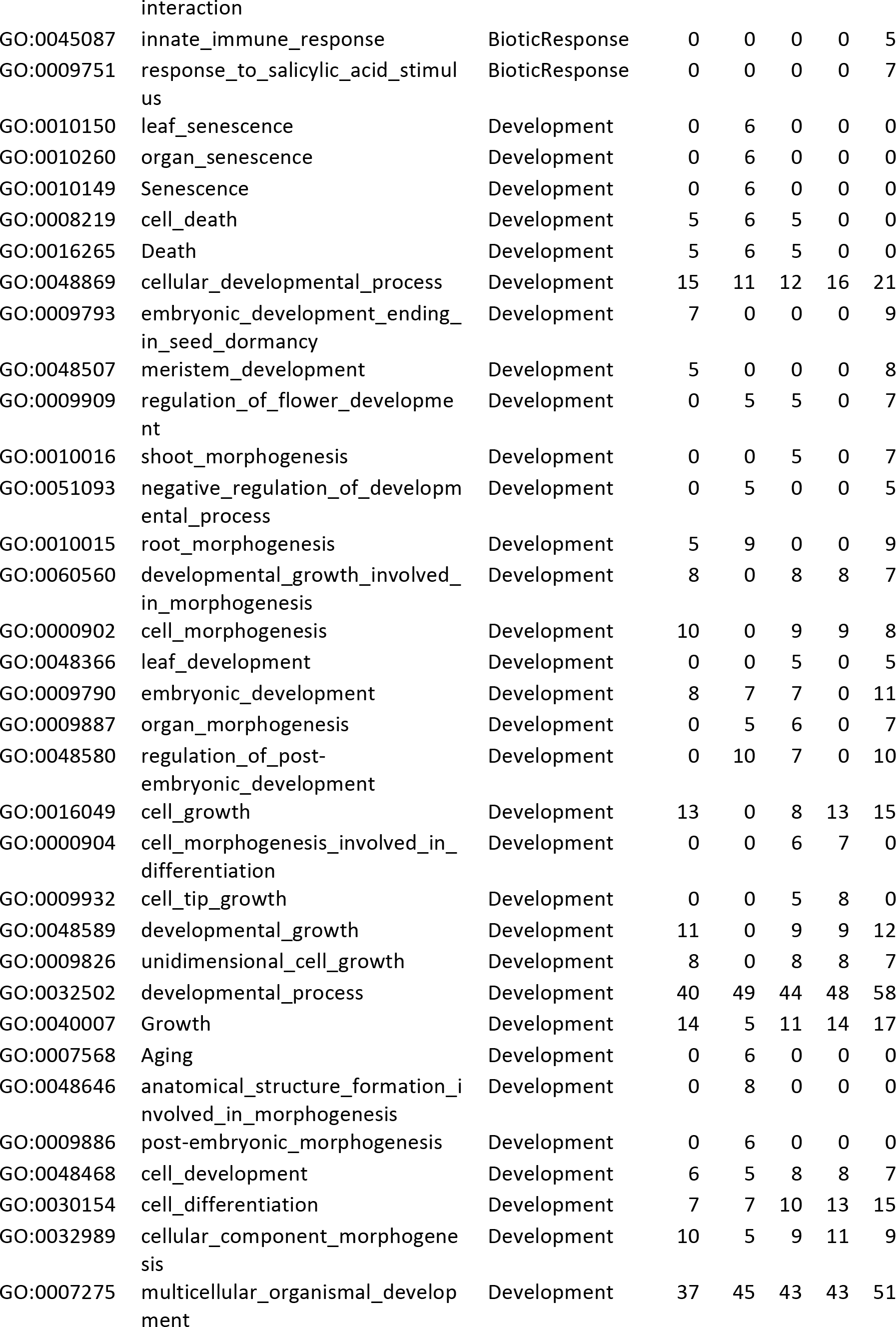
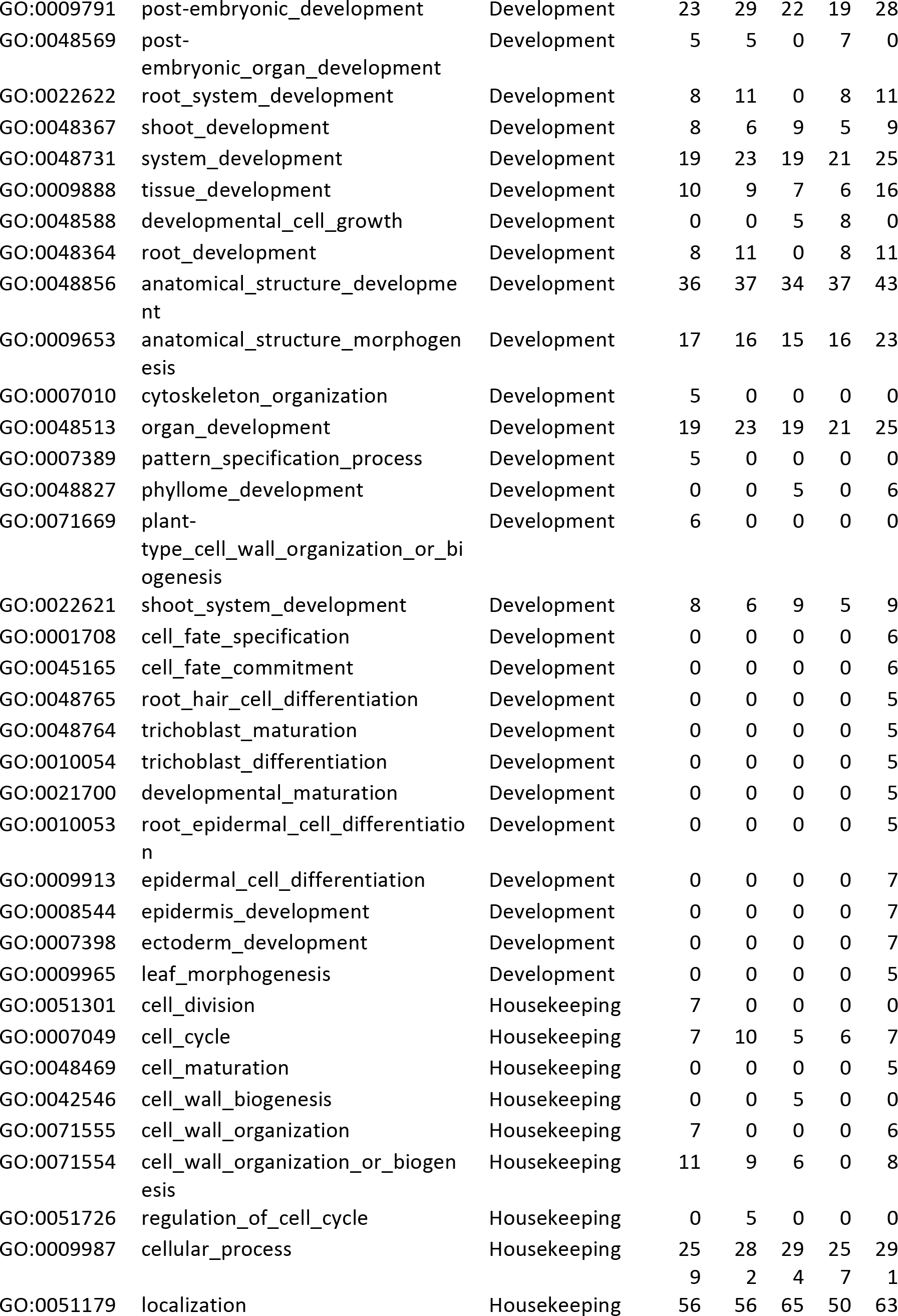
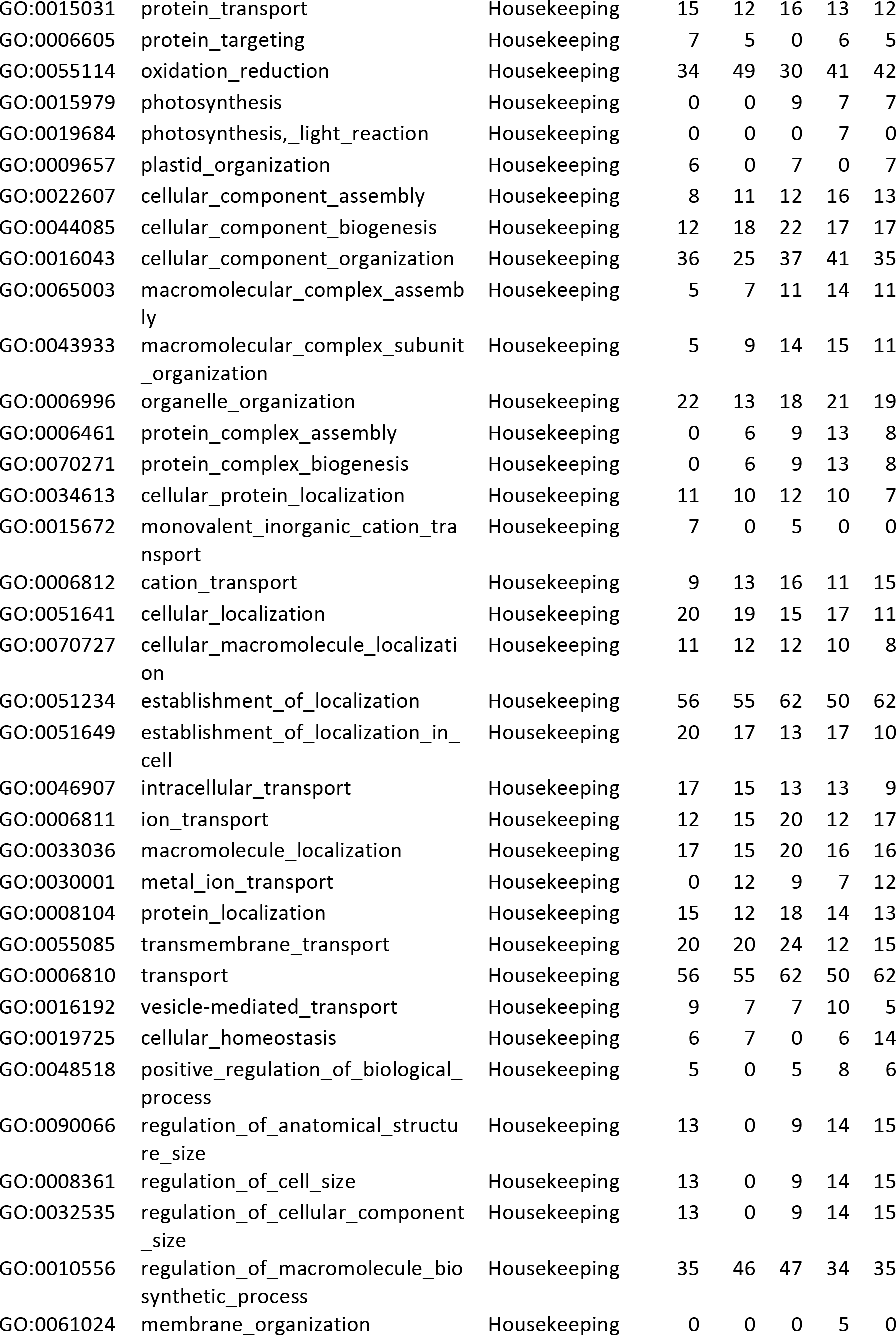
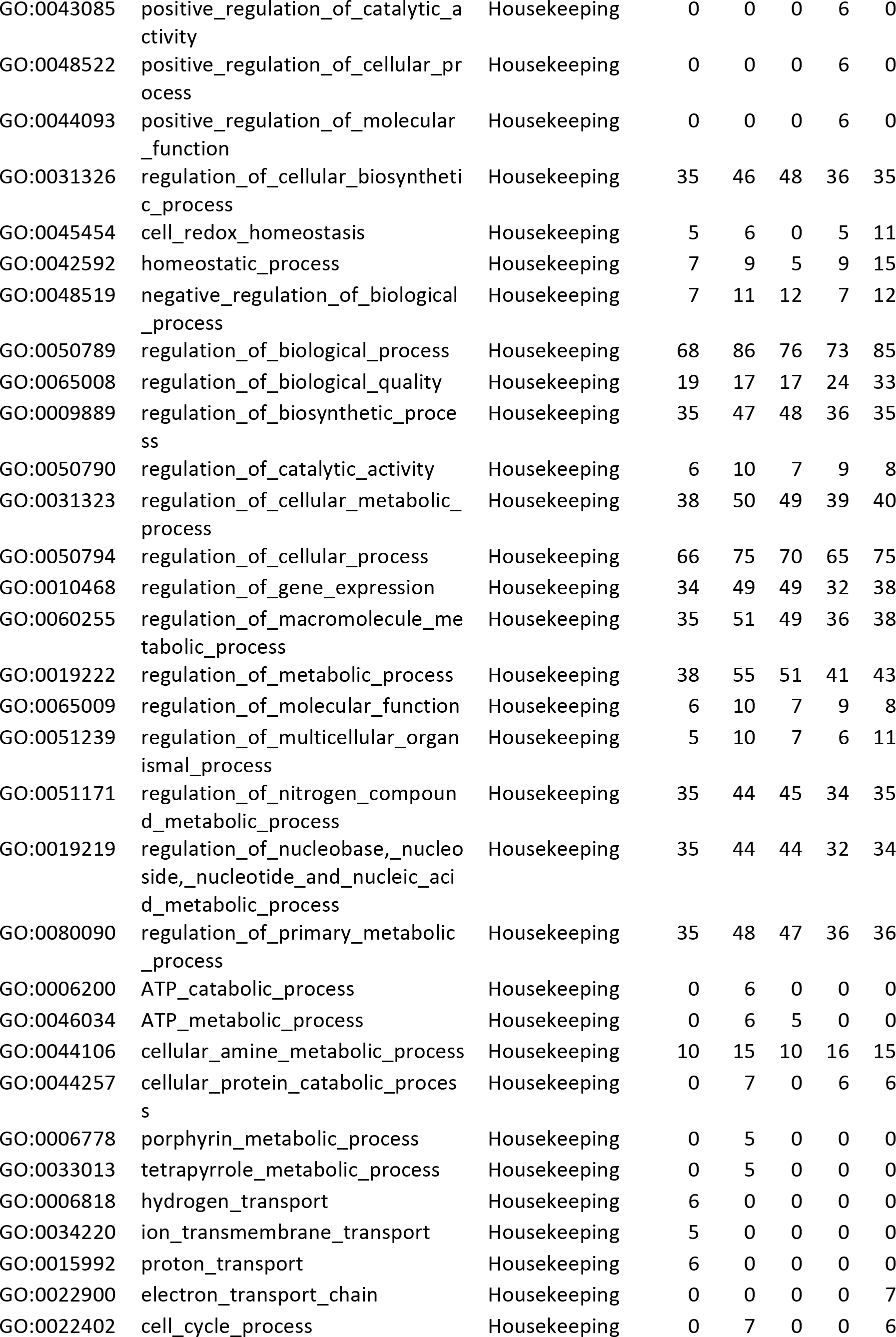
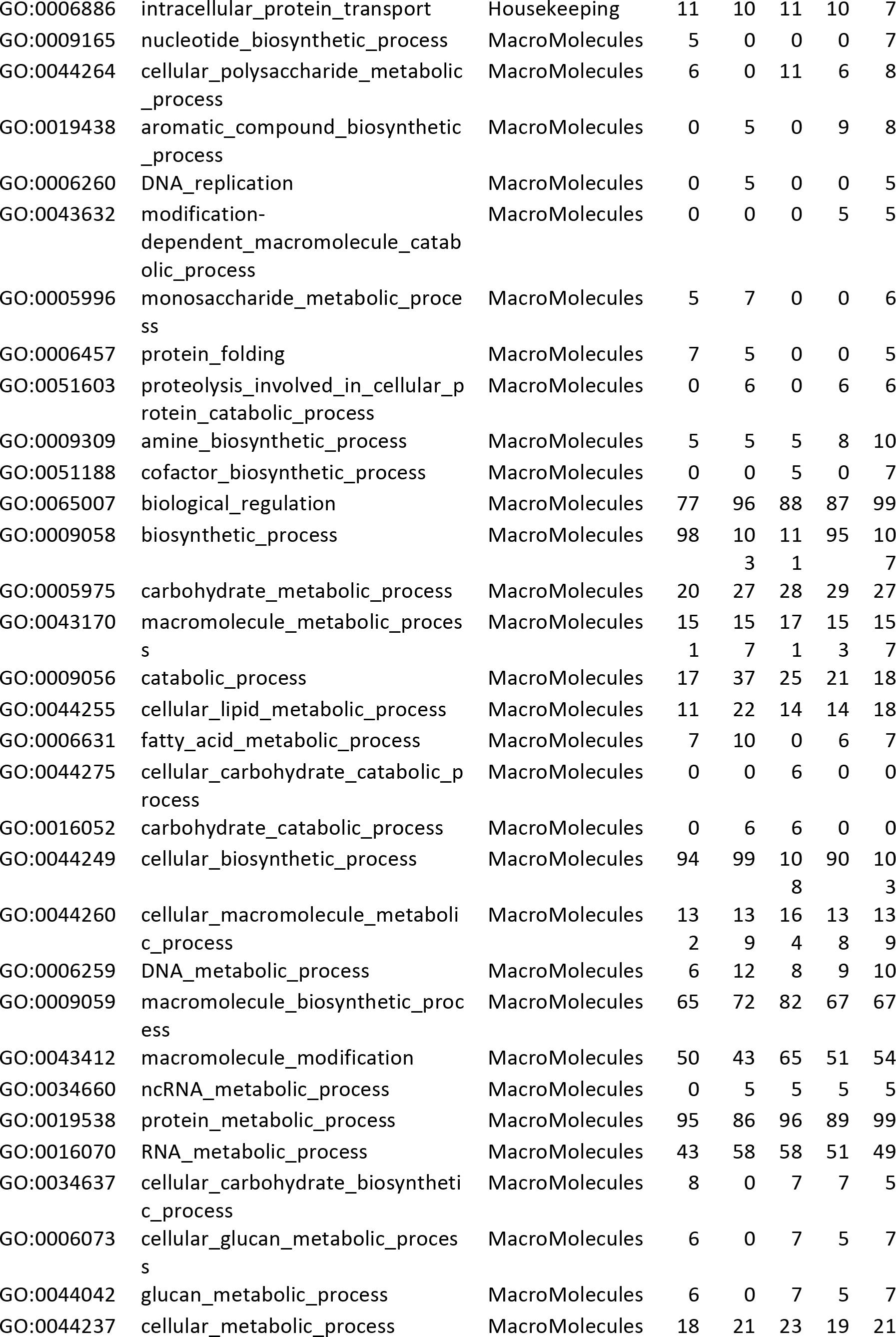
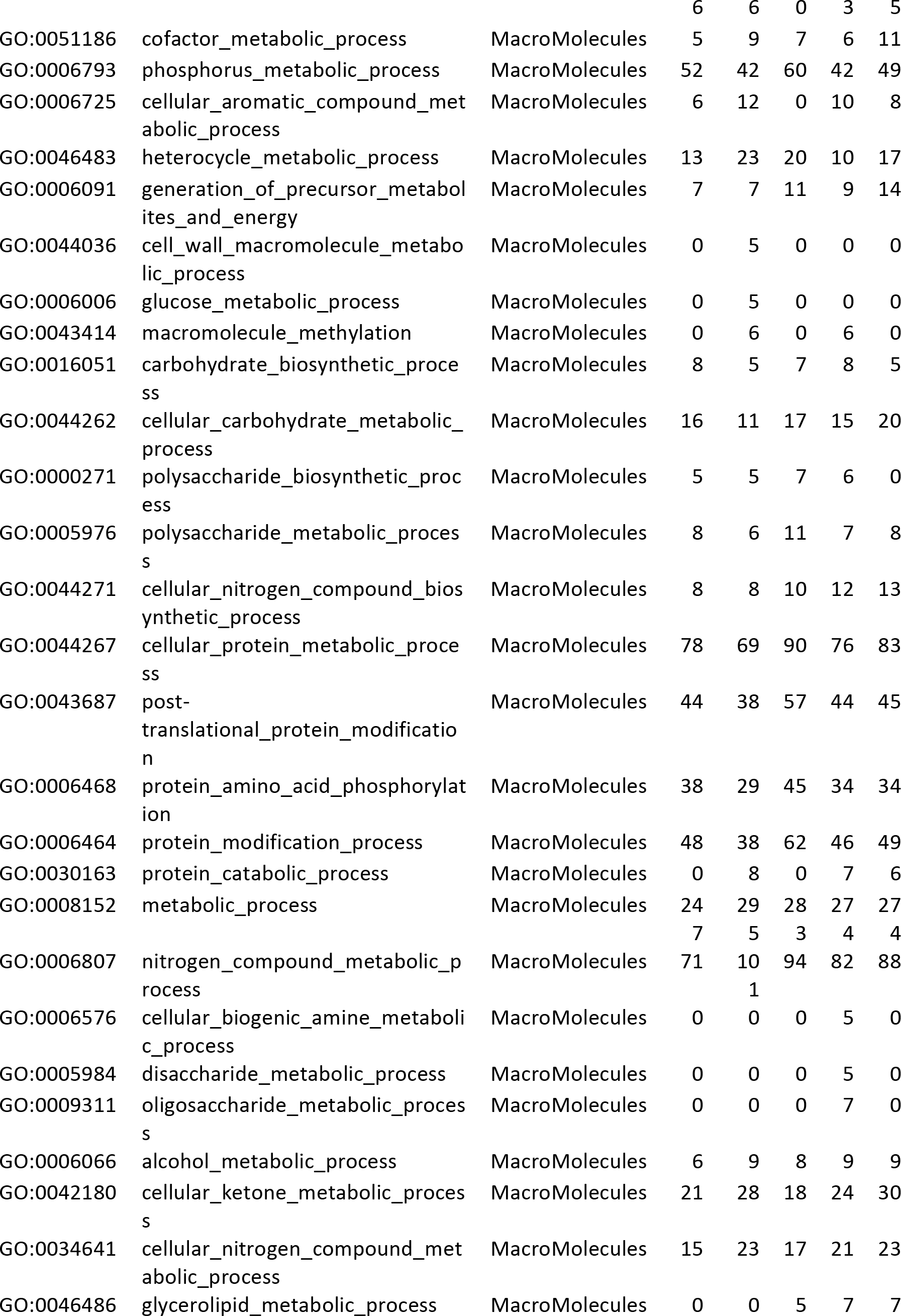
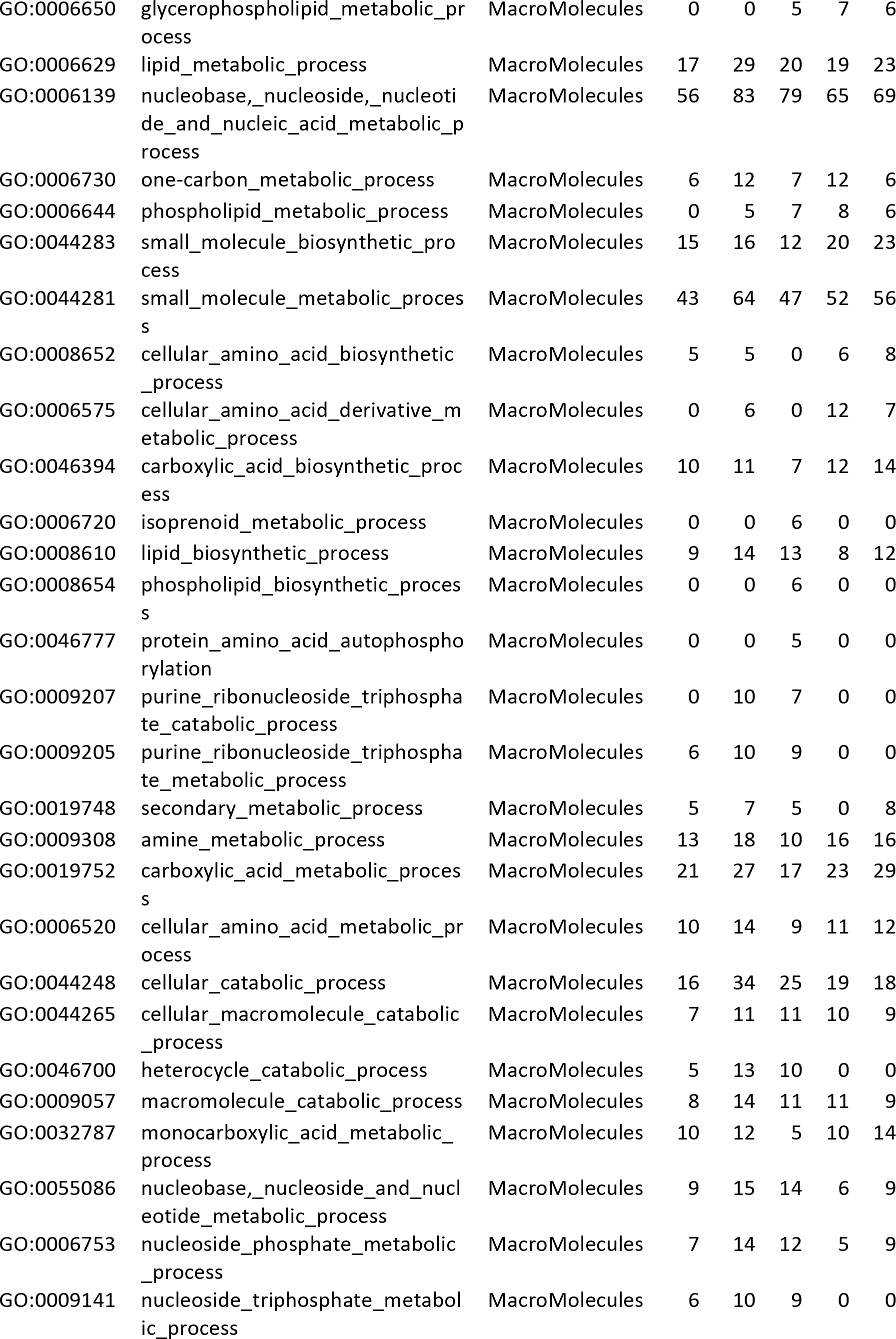
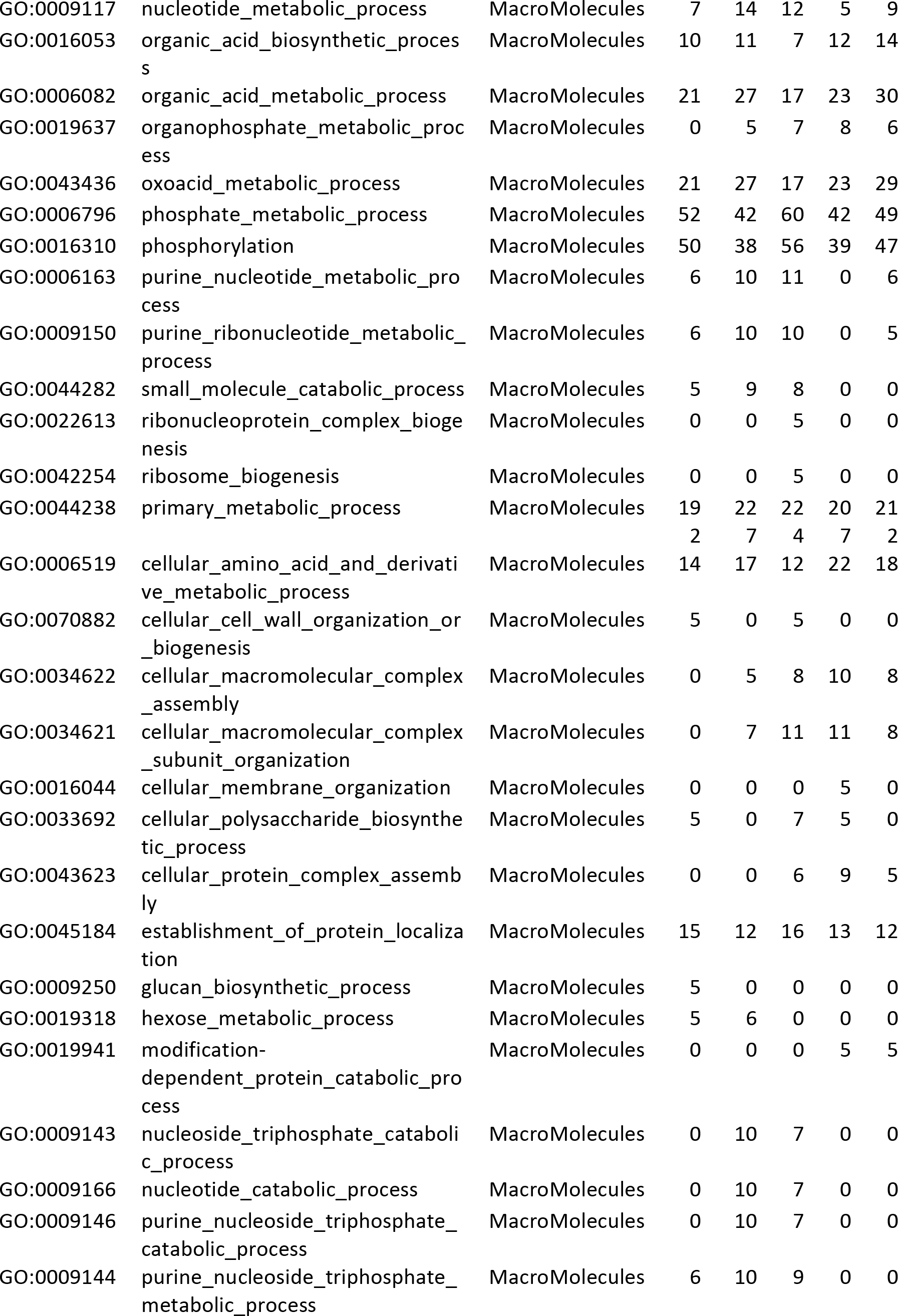
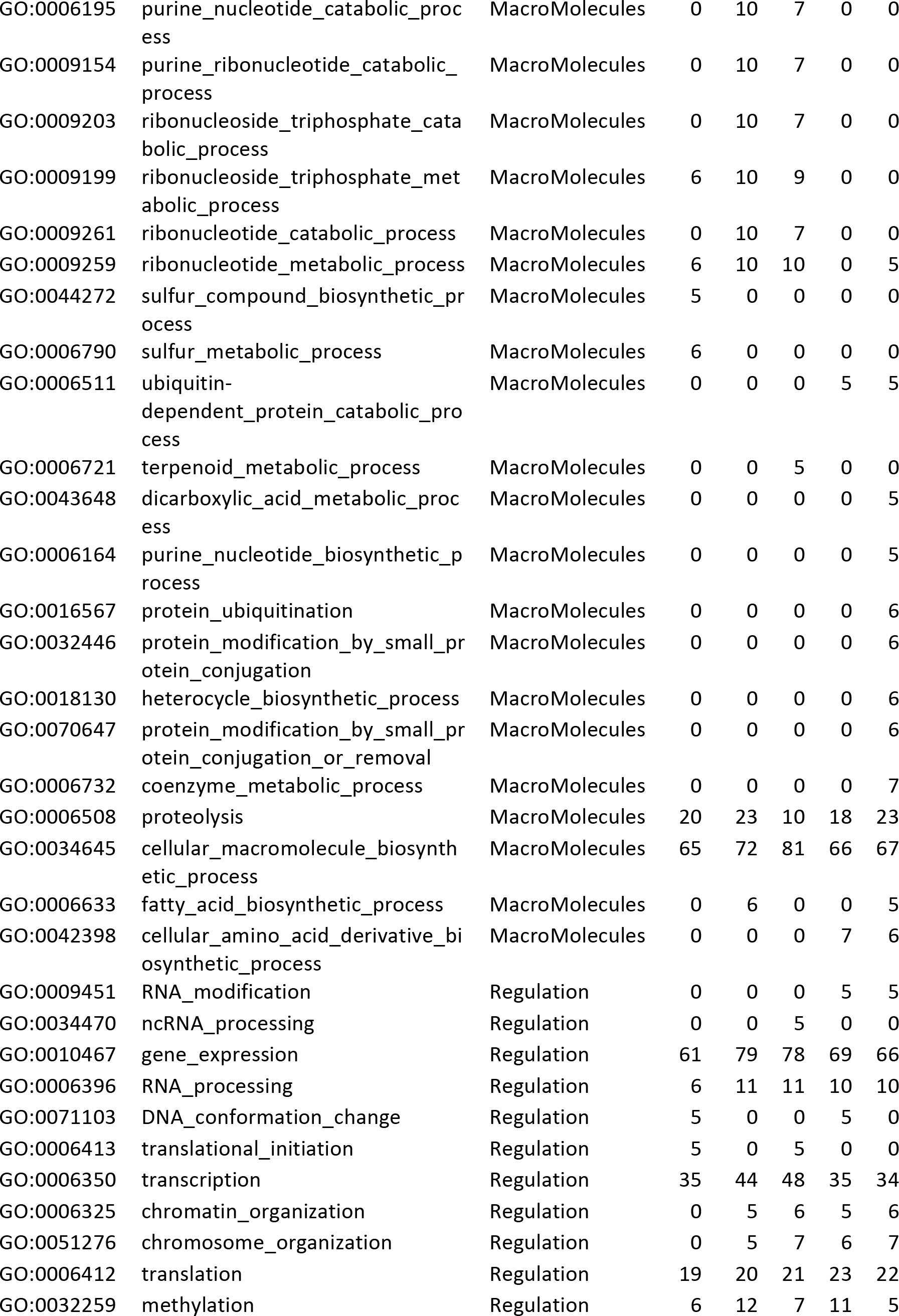
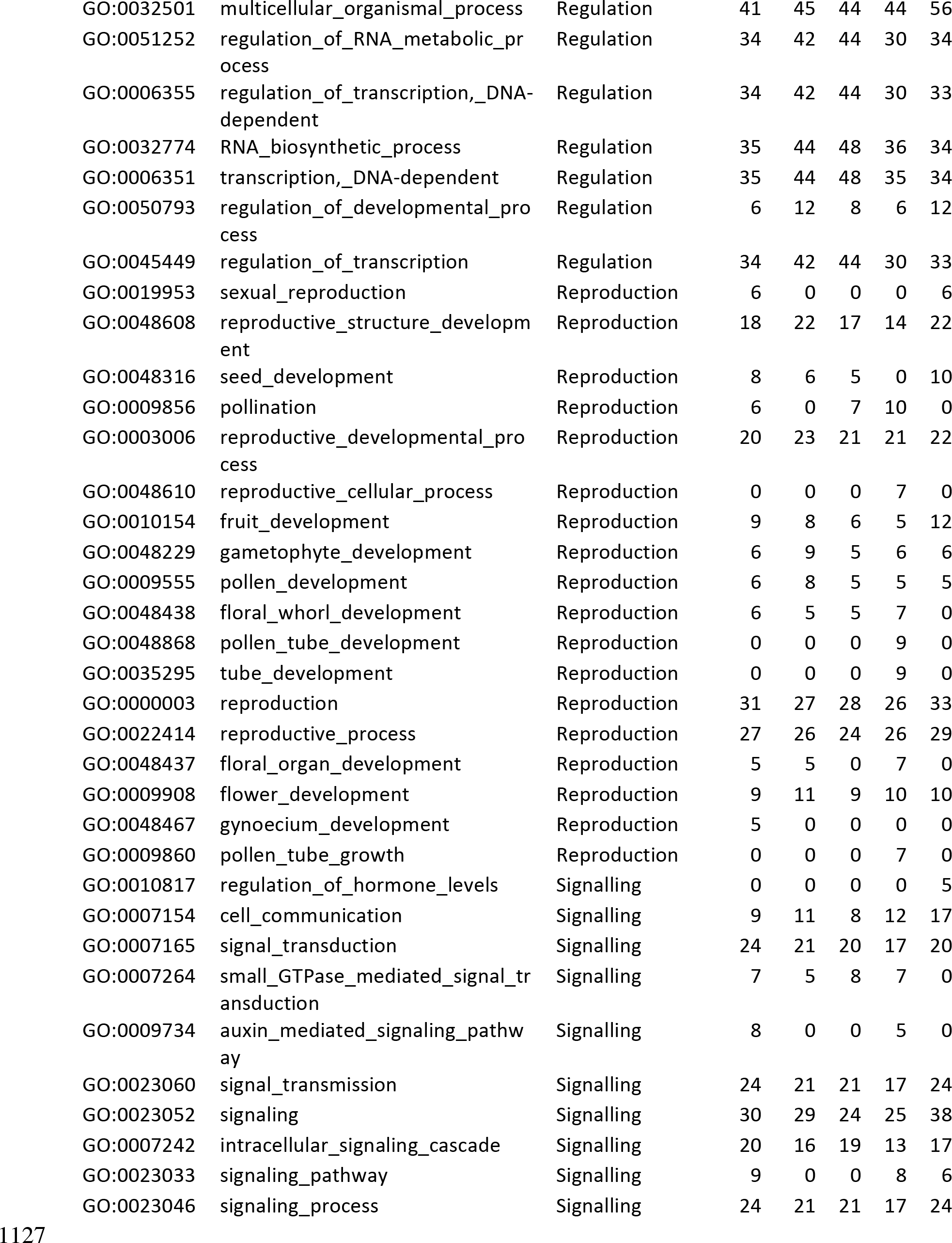
GO term classification f 1126 or top 1,000 gene models by BRHM

**Table S6.** Proportion of candidate genes in significant regions relative to random random genes within significant regions. Significance based on Bonferroni corrected significance for BRHM method, Benjamini-Hochberg for RHM, and for SNPs selected in at least a certain number of models for the RMIP method. Random genes in significant regions are the average of 100 samples. BRHM results duplicated from Table 5.

|  | <u>DTA</u> |  |  | <u>DTS</u> |  |  |
| --- | --- | --- | --- | --- | --- | --- |
| **RMIP** | 3 | 2 | 1 | 3 | 2 | 1 |
| Ames | 1.953 (5) | 1.759 (7) | 1.497 (14) | 1.24 (3) | 0.75 (3) | 1.086 (11) |
| CNNAM | 1.404 (4) | 1.931 (9) | 1.304 (17) | 2.214 (6) | 1.709 (8) | 1.239 (17) |
| EUNAM-Dent | 5.556 (4) | 4.167 (5) | 2.353 (8) | 2.02 (2) | 2 (2) | 2.141 (7) |
| EUNAM-Flint | 0 (0) | 0 (0) | 0.943 (3) | 0 (0) | 0 (0) | 0.356 (1) |
| USNAM | 1.339 (3) | 1.016 (5) | 0.84 (10) | 1.688 (4) | 0.899 (4) | 1.227 (14) |
| **RHM** | 0.001 | 0.01 | 0.05 | 0.001 | 0.01 | 0.05 |
| Ames | 12 (3) | 6.25 (3) | 2.013 (3) | 2.548 (8) | 1.224 (10) | 1.364 (16) |
| CNNAM | 0 (0) | 5.882 (4) | 2.844 (6) | 0 (0) | 1.235 (1) | 2.315 (5) |
| EUNAM.Dent | 0 (0) | 0 (0) | 0 (0) | 0 (0) | 0 (0) | 0 (0) |
| EUNAM.Flint | 2 (2) | 1.653 (2) | 1.117 (2) | 1.471 (1) | 1.19 (1) | 1.55 (2) |
| USNAM | 3.704 (3) | 2.5 (6) | 1.621 (16) | 1.432 (6) | 1.332 (23) | 1.166 (35) |
| **BRHM** | 0.001 | 0.01 | 0.05 | 0.001 | 0.01 | 0.05 |
| Ames | 0.968 (6) | 1.316 (10) | 1.22 (11) | 1.297 (8) | 1.282 (9) | 1.424 (13) |
| CNNAM | 1.579 (6) | 1.777 (11) | 1.31 (11) | 2.174 (8) | 2.609 (15) | 1.724 (16) |
| EUNAM-Dent | 1.019 (7) | 1.046 (8) | 1.012 (12) | 1.372 (9) | 1.171 (10) | 1.033 (14) |
| EUNAM-Flint | 1.156 (8) | 0.937 (8) | 0.915 (13) | 1.441 (10) | 1.202 (12) | 0.904 (13) |
| USNAM | 1.146 (28) | 1.095 (32) | 1.044 (38) | 1.139 (29) | 1.16 (36) | 1.164 (42) |

\*Candidates in significant regions/Average over 100 samples of non-candidate genes from the maize v3 gene models in significant regions (Number of candidates in significant regions)

